## supplemental file for "Amidase and Lysozyme Dual Functions in TseP Reveal a New Family of Chimeric Effectors in the Type VI Secretion System"

Zeng-Hang Wang *et al*.

* Tao Dong.

* Wenming Qin.

**Supplemental Figure S1. Staining SDS-PAGE of secretion of TseP, TseP$^N$, and TseP$^C$ in the SSU triple effector deletion mutant (Δ*3eff*).** The secretion proteins were visualized by SDS-PAGE and stained with Coomassie brilliant blue dye.

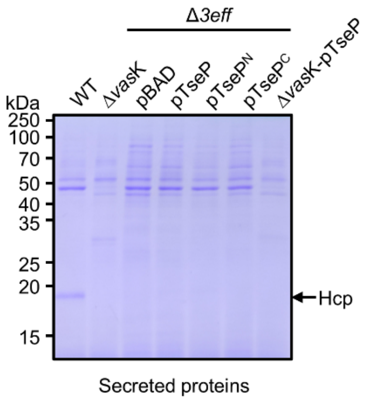

**Supplemental Figure S2. TsiP inhibits both the amidase and lysozyme activities of TseP.** The immunity protein TsiP was incubated with TseP or TseP$^{C}$ on ice for 12 h before being mixed with PG. Products were analyzed by UPLC-QTOF MASS spectrometry.

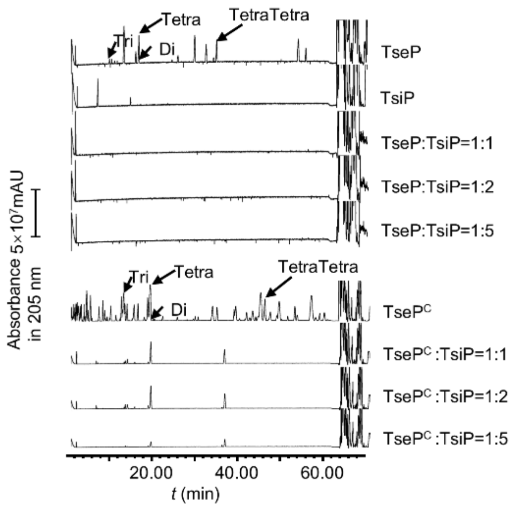

**Supplemental Figure S3. The amidase activity of TseP is not required for either T6SS assembly or lysozyme function. A,** Time-lapse imaging of VipA-sfGFP signals in the Δ*3eff* mutant complemented with different TseP amidase-inactive mutants. Each sample was captured every 10 s for 5 min and temporally color-coded. Color scale used to temporally color code the VipA-sfGFP signals is shown at the bottom. A 30 × 30 μm representative field of cells is shown. Scale bars, 5 μm. **B,** The maximum enzymatic activity of TseP and its variants. PG substrates (1.5 mg/ml) were treated with 10 nM TseP or its variants, respectively. **C,** Statistical analysis of T6SS sheath assemblies in the Δ*3eff* mutant complemented with different TseP variants. For **B** and **C**, error bars indicate the mean ± standard deviation of three biological replicates, and statistical significance was calculated using a two-tailed Student's *t*-test. ns, not significant; *, $P < 0.05$; **, $P < 0.01$.

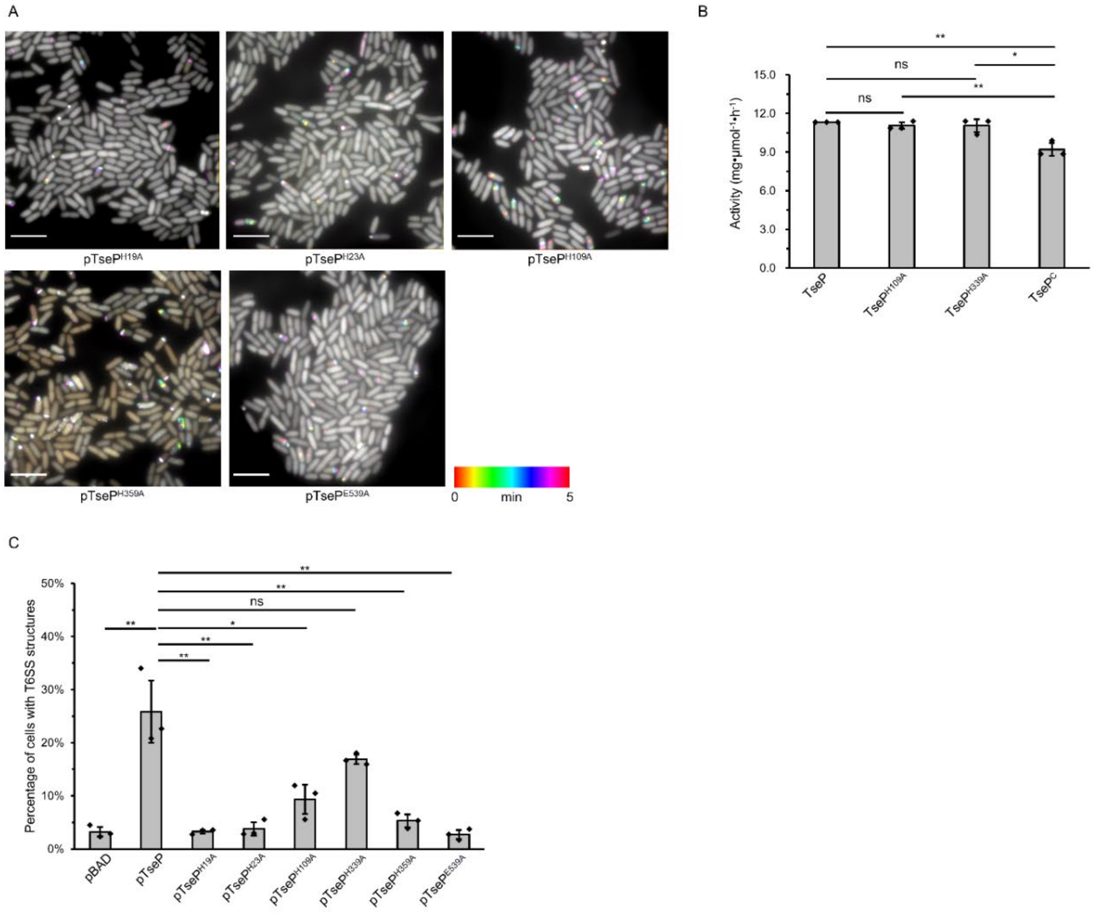

**Supplemental Figure S4. Protein sequence analysis of the TseP homologs.** Sequences were retrieved from UniProt and aligned using Jalview. Alignment view was generated using ESPript 3 with default settings. Proteins tested in this study are highlighted in red. The amidase active sites are marked with a black triangle at the bottom.

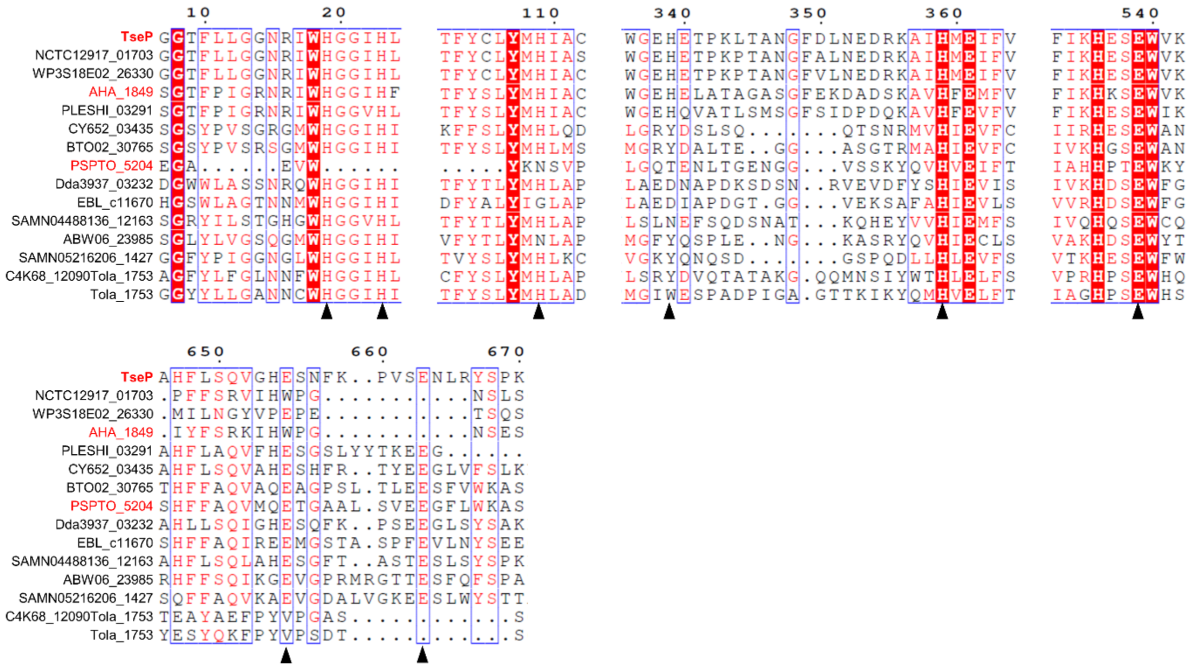

**Supplemental Figure S5. Structural and mutational analyses of the C-terminal domain. A,** Secondary structure analysis of TseP$^{C}$. Data was calculated using EMBL-EBI webs with PDBsum tool (https://www.ebi.ac.uk/thornton-srv/databases/pdbsum/). **B,** Electrostatic potential maps of the TseP$^{C}$ and lysozyme. The electrostatic surface potentials were colored red for negative charges, blue for positive charges, and white for neutral residues. **C,** Digestion of *B. subtilis* PG with different TseP variants. Exponential phase *B. subtilis* cells ($OD_{600}$~1.0) were used as substrates and the lysis percentage was calculated by detecting the changes in optical density at 600 nm. **D,** Statistical analysis of killer cells during competition assays for which the survival of prey cells is shown in Figure 6F. For **C** and **D**, error bars indicate the mean ± standard deviation of three biological replicates, and statistical significance was calculated using a two-tailed Student's *t*-test. ns, not significant; **, $P < 0.01$.

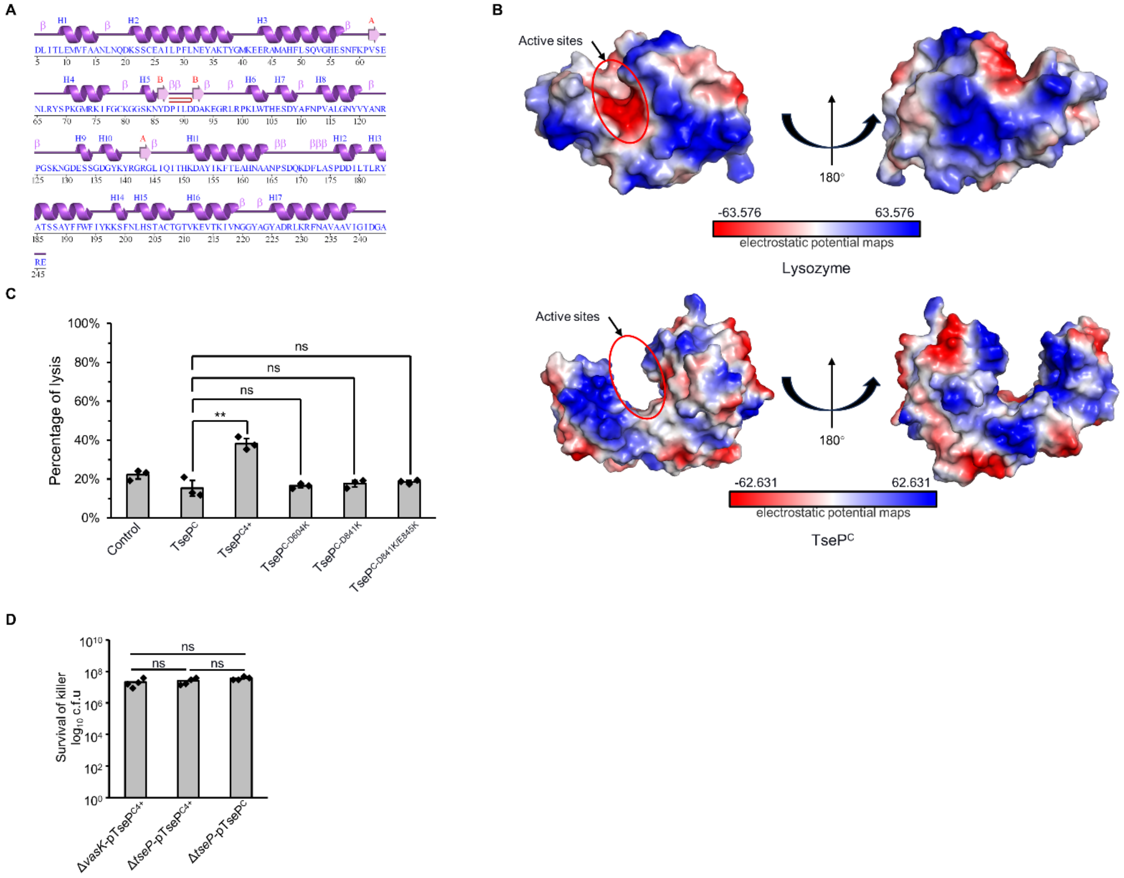

**Supplemental Figure S6. EF-hand domain has no effect on TseP activity. A,** Protein sequence alignment (top) and structure alignment (bottom) of the EF-hand domain of TseP to the known EF-hand domain-containing protein $_{CD}$EFhd2 (PDB ID: 5H0P) (59). **B,** Statistical analysis of T6SS sheath assemblies in the Δ*3eff* mutant complemented with different TseP variants. Error bars indicate the mean ± standard deviation of three biological replicates, and statistical significance was calculated using a two-tailed Student's *t*-test. ns, not significant; **, $P < 0.01$. **C,** Competition assays of the Δ*3eff* mutant complemented with different TseP variants against *E. coli* MG1655. Competition assays were repeated twice.

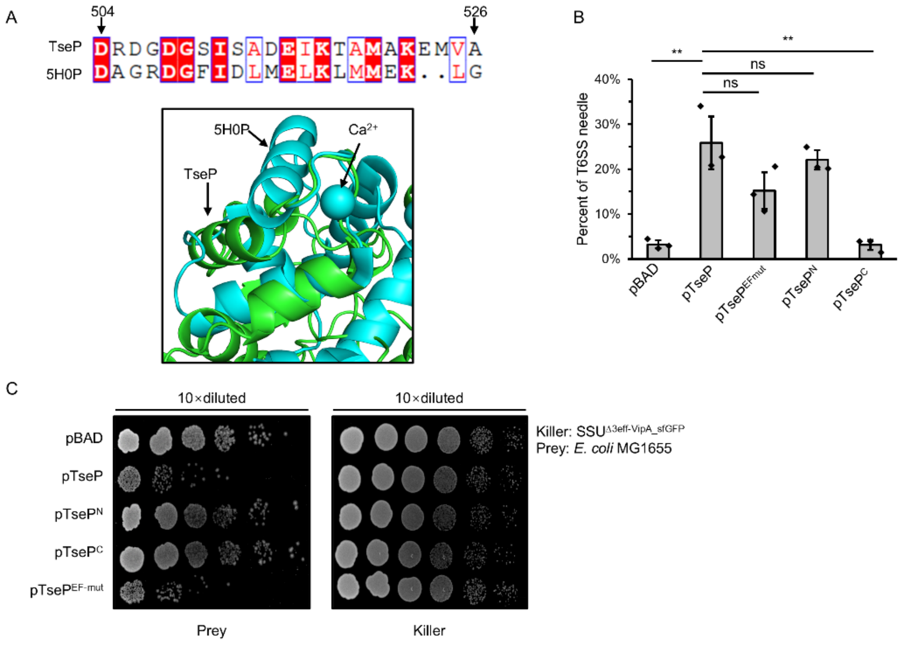

**Supplemental Figure S7. Expression and secretion of TseP variants. A,** Secretion analysis of Hcp in the Δ*3eff* mutant complemented with different TseP variants. RpoB serves as an equal loading and autolysis control. Hcp, RpoB, and 3V5-tagged TseP proteins were detected using specific antibodies. **B,** Protein expression of TseP and its variants in *E. coli* and SSU strains. RpoB serves as an equal loading control. RpoB and 3V5-tagged TseP proteins were detected using specific antibodies.

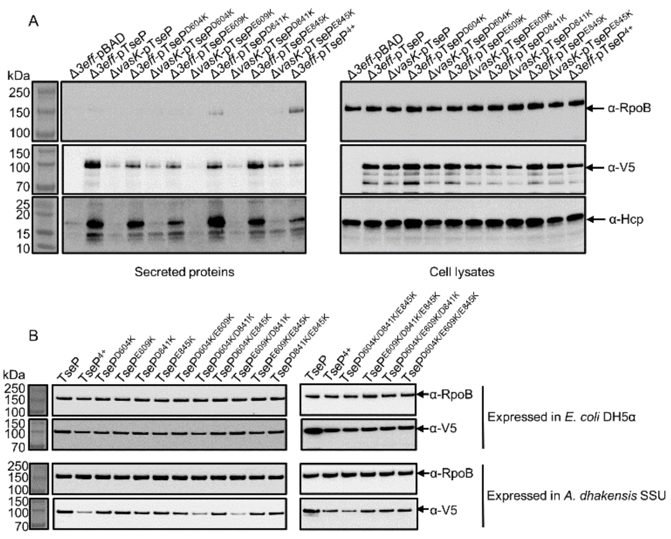

**Supplemental Figure S8. Purification of TseP variants and homologous proteins. A,** Purification of His-TseP, His-TseP$^{E663A}$, His-SUMO-TseP$^{N}$, His-SUMO-TseP$^{C}$, and His-SUMO-TseP$^{C\text{-}E663A}$. The His-SUMO tag was removed by SUMO protease. Proteins were used for PG-digestion analysis in Figure 2A, 2D, and 2E. **B,** Purification of TseP proteins with or without EDTA. Proteins were used for PG-digestion analysis in Figure 2E. **C,** Purification of TseP$^{N}$ variants with a His-SUMO tag. The His-SUMO tag was removed by SUMO protease. Proteins were used for PG-digestion analysis in Figure 2D. **D,** Purification of AHA_1849, PSPTO_5204, and their mutants. Proteins were used for PG-digestion analysis in Figure 4B and 4C. **E,** Purification of TseP$^{C}$ and TseP$^{C\text{-}E663D}$ with an His-SUMO tag. The His-SUMO tag was removed by SUMO protease. Proteins were used for PG-lysis in Figure 5E. **F,** Purification of TseP$^{C}$ variants with an His-SUMO tag. The His-SUMO tag was removed by SUMO protease. Proteins were used for PG-lysis in Figure 6C and Figure S4C. All these proteins were purified with Ni-NTA affinity chromatography column, eluted using imidazole, and analyzed via SDS-PAGE.

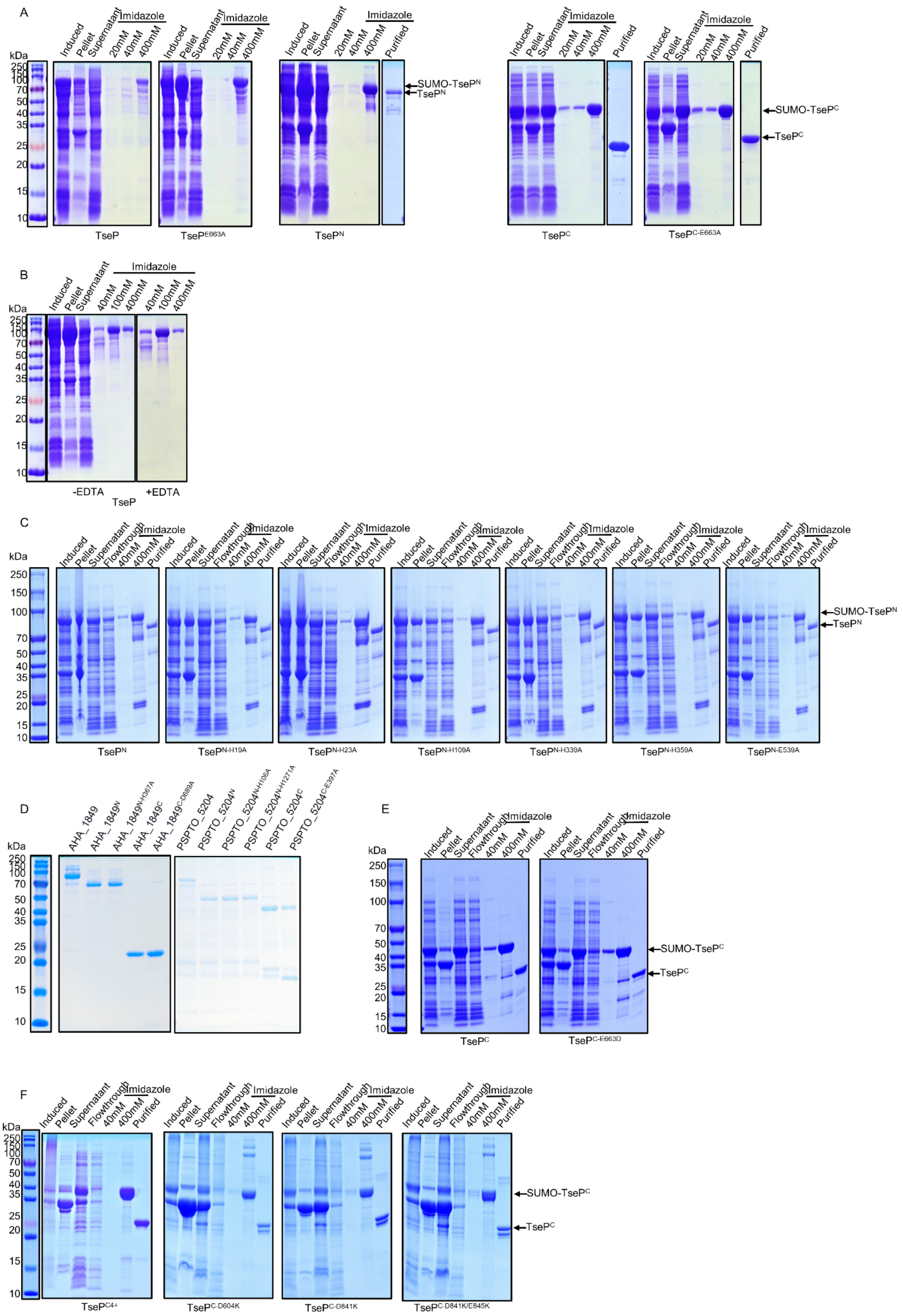
A
kDa
Induced
Pellet
Supernatant
Imidazole
20mM
40mM
400mM
Purified
250
150
100
70
50
40
35
25
20
15
10
SUMO-TseP^N
TseP^N
SUMO-TseP^C
TseP^C
TseP
TseP^E663A
TseP^N
TseP^C
TseP^C-E663A
B
100mM
-EDTA
+EDTA
TseP
C
Flowthrough
TseP^N-H19A
TseP^N-H23A
TseP^N-H109A
TseP^N-H339A
TseP^N-H359A
TseP^N-E539A
D
AHA_1849
AHA_1849^N
AHA_1849^N-H387A
AHA_1849^C
AHA_1849^C-D689A
PSPTO_5204
PSPTO_5204^N
PSPTO_5204^N-H106A
PSPTO_5204^N-H127A
PSPTO_5204^C
PSPTO_5204^C-E397A
E
TseP^C-E663D
F
TseP^C4+
TseP^C-D604K
TseP^C-D841K
TseP^C-D841K/E845K
