## Supplemental Table S1-3 for "Amidase and Lysozyme Dual Functions in TseP Reveal a New Family of Chimeric Effectors in the Type VI Secretion System"

**Supplementary Table S1.** Strains and plasmids used in this study.

| Strain | Genotype | Description | Source |
|---|---|---|---|
| *E. coli* | | | |
| BL21(DE3) | *F– ompT gal dcm lon hsdSB(rB–mB–) λ(DE3 [lacI lacUV5T7p07 ind1 sam7 nin5]) [malB+]K-12(λS)* | Strain used for protein expression | Lab stock |
| DH5α | *F- Φ80lacZΔM15 Δ(lacZYA-argF) U169 recA1 endA1 hsdR17 phoA supE44 thi-1 gyrA96 relA1 λ-* | Strain used for cloning | Invitrogen |
| T-fast | *F' proA+ B+ lacIq ΔlacZM15 / fhuA2 Δ(lac-proAB) glnV galK16 galE15 R(zgb-210::Tn10)TetR endA1 thi-1 Δ(hsdS-mcrB)5* | Strain used for cloning | TIANGEN |
| MG1655 | *F-, lambda-, rph-1* | K-12 wild-type strain used as prey for T6SS killing | Lab stock |
| *Aeromonas dhakensis* SSU | WT | Parental strain | [1] |
| | Δ*vasK* | T6SS null, in-frame deletion of *vasK* | [1] |
| | Δ*tseP* | In-frame deletion of *tseP* | [1] |
| | Δ*3eff*, *vipA-sfgfp* | In-frame deletion of *tseP*, *tseI*, and *tseC*, and fusion expression of sfGFP protein and VipA protein. | This study |

### Plasmid

| | Plasmid | Description | Source |
|---|---|---|---|
| pBAD24 | pBAD24Kan | Arabinose inducible expression plasmid, kanamycin resistance | Lab stock |
| | pBAD24kan-VgrG2-TseP-V5 | Arabinose inducible expression of VgrG2 and TseP | This study |
| | pBAD24kan-VgrG2-TseP$^{N}$-V5 | Arabinose inducible expression of VgrG2 and TseP$^{N}$ | This study |
| | pBAD24kan-VgrG2-TseP$^{C}$-V5 | Arabinose inducible expression of VgrG2 and TseP$^{C}$ | This study |

| | | | |
|---|---|---|---|
| | pBAD24kan-VgrG2-TseP$^{H19A}$-V5 | Arabinose inducible expression of VgrG2 and TseP $^{H19A}$ | This study |
| | pBAD24kan-VgrG2-TseP$^{H23A}$-V5 | Arabinose inducible expression of VgrG2 and TseP$^{H23A}$ | This study |
| | pBAD24kan-VgrG2-TseP$^{H109A}$-V5 | Arabinose inducible expression of VgrG2 and TseP$^{H109A}$ | This study |
| | pBAD24kan-VgrG2-TseP$^{H339A}$-V5 | Arabinose inducible expression of VgrG2 and TseP$^{H339A}$ | This study |
| | pBAD24kan-VgrG2-TseP$^{H359A}$-V5 | Arabinose inducible expression of VgrG2 and TseP$^{H359A}$ | This study |
| | pBAD24kan-VgrG2-TseP$^{E539A}$-V5 | Arabinose inducible expression of VgrG2 and TseP$^{E539A}$ | This study |
| | pBAD24kan-VgrG2-TseP$^{EFhandmut}$-V5 | Arabinose inducible expression of VgrG2 and TseP$^{EFhandmut}$ | This study |
| pBAD18 | pBAD18Cm | Arabinose inducible expression plasmid, choloramphenicol resistance | Lab stock |
| pET28a | | IPTG inducible expression plasmid, gentamycin resistant | Lab stock |
| | pET28a-His-SUMO-TseP | IPTG inducible expression of SUMO-TseP fusion with an N-terminal 6×His tag | Lab stock |
| | pET28a-His-SUMO-TseP$^{E663A}$ | IPTG inducible expression of SUMO-TseP$^{E663A}$ fusion with an N-terminal 6×His tag | [2] |
| | pET28a-His-SUMO-TseP$^{N}$ | IPTG inducible expression of SUMO-TseP fusion with an N-terminal 6×His tag | This study |
| | pET28a-His-SUMO-TseP$^{N-H19A}$ | IPTG inducible expression of SUMO-TseP$^{N-H19A}$ fusion with an N-terminal 6×His tag | This study |
| | pET28a-His-SUMO-TseP$^{N-H23A}$ | IPTG inducible expression of SUMO-TseP$^{N-H23A}$ fusion with an N-terminal 6×His tag | This study |
| | pET28a-His-SUMO-TseP$^{N-H109A}$ | IPTG inducible expression of SUMO-TseP$^{N-H109A}$ fusion with an N-terminal 6×His tag | This study |
| | pET28a-His-SUMO-TseP$^{N-H339A}$ | IPTG inducible expression of SUMO-TseP$^{N-H339A}$ fusion with an N-terminal 6×His tag | This study |
| | pET28a-His-SUMO-TseP$^{N-H359A}$ | IPTG inducible expression of SUMO-TseP$^{N-H359A}$ fusion with an N-terminal 6×His tag | This study |
| | pET28a-His-SUMO-TseP$^{N-E539A}$ | IPTG inducible expression of SUMO-TseP$^{N-E539A}$ fusion with an N-terminal 6×His tag | This study |
| | pET28a-His-SUMO-TseP$^{C}$ | IPTG inducible expression of SUMO-TseP$^{C}$ fusion with an N-terminal 6×His tag | This study |

| | | |
|---|---|---|
| pET28a-His-SUMO-TseP$^{C\text{-}E663D}$ | IPTG inducible expression of SUMO-TseP$^{C\text{-}E663D}$ fusion with an N-terminal 6×His tag | This study |
| pET28a-His-SUMO-TseP$^{C4+}$ | IPTG inducible expression of SUMO-TseP$^{C4+}$ fusion with an N-terminal 6×His tag | This study |
| pET28a-His-SUMO-TseP$^{C\text{-}D604K}$ | IPTG inducible expression of SUMO-TseP$^{C\text{-}D604K}$ fusion with an N-terminal 6×His tag | This study |
| pET28a-His-SUMO-TseP$^{C\text{-}D841K}$ | IPTG inducible expression of SUMO-TseP$^{C\text{-}D841K}$ fusion with an N-terminal 6×His tag | This study |
| pET28a-His-SUMO-TseP$^{C\text{-}D841K/E845K}$ | IPTG inducible expression of SUMO-TseP$^{C\text{-}D841K/E845K}$ fusion with an N-terminal 6×His tag | This study |
| pET28a-His-SUMO-TsiP | IPTG inducible expression of SUMO-TsiP fusion with an N-terminal 6×His tag | This study |
| pET28a-His-SUMO-VgrG2 | IPTG inducible expression of SUMO-VgrG2 fusion with an N-terminal 6×His tag | This study |
| pET28a-His-SUMO-AHA_1849 | IPTG inducible expression of SUMO-AHA_1849 fusion with an N-terminal 6×His tag | This study |
| pET28a-His-SUMO-AHA_1849$^{N}$ | IPTG inducible expression of SUMO-AHA_1849$^{N}$ fusion with an N-terminal 6×His tag | This study |
| pET28a-His-SUMO-AHA_1849$^{N\text{-}H369A}$ | IPTG inducible expression of SUMO- AHA_1849$^{N\text{-}H369A}$ fusion with an N-terminal 6×His tag | This study |
| pET28a-His-SUMO-AHA_1849$^{C}$ | IPTG inducible expression of SUMO-AHA_1849$^{C}$ fusion with an N-terminal 6×His tag | This study |
| pET28a-His-SUMO-AHA_1849$^{C\text{-}D689A}$ | IPTG inducible expression of SUMO-AHA_1849$^{C\text{-}D689A}$ fusion with an N-terminal 6×His tag | This study |
| pET28a-His-SUMO-PSPTO_5204 | IPTG inducible expression of SUMO-PSPTO_5204 fusion with an N-terminal 6×His tag | This study |
| pET28a-His-SUMO-PSPTO_5204$^{N}$ | IPTG inducible expression of SUMO-PSPTO_5204$^{N}$ fusion with an N-terminal 6×His tag | This study |
| pET28a-His-SUMO-PSPTO_5204$^{N\text{-}H106A}$ | IPTG inducible expression of SUMO-PSPTO_5204$^{N\text{-}H106A}$ fusion with an N-terminal 6×His tag | This study |
| pET28a-His-SUMO-PSPTO_5204$^{N\text{-}H271A}$ | IPTG inducible expression of SUMO-PSPTO_5204$^{N\text{-}H271A}$ fusion with an N-terminal 6×His tag | This study |
| pET28a-His-SUMO-PSPTO_5204$^{C}$ | IPTG inducible expression of SUMO-PSPTO_5204$^{C}$ fusion with an N-terminal 6×His tag | This study |
| pET28a-His-SUMO-PSPTO_5204$^{C\text{-}E397A}$ | IPTG inducible expression of SUMO-PSPTO_5204$^{C\text{-}E397A}$ fusion with an N-terminal 6×His tag | This study |

**Supplementary Table S2.** The sequence identity of the N-terminal of TseP and homologs. Sequence identity was calculated using ClustalW web server. Colors are assigned from blue to red indicating low to high sequence similarity, respectively.

| | 1 | 2 | 3 | 4 | 5 | 6 | 7 | 8 | 9 | 10 | 11 | 12 | 13 | 14 | 15 |
|---|---|---|---|---|---|---|---|---|---|---|---|---|---|---|---|
| **TseP** | | 41.8 | 88.2 | 88.7 | 47.1 | 12.4 | 14.3 | 12.1 | 13 | 12.1 | 13.3 | 12.6 | 12.3 | 12.5 | 16.7 |
| NCTC12917_01703 | 41.8 | | 42.9 | 41.1 | 60.6 | 14.9 | 13.7 | 13 | 14.7 | 13.9 | 13.3 | 10.9 | 10 | 14.4 | 17.4 |
| WP3S18E02_26330 | 88.2 | 42.9 | | 88.5 | 46.8 | 13 | 12.8 | 10.9 | 15.1 | 12 | 12.8 | 12.4 | 12 | 13.7 | 17 |
| AHA_1849 | 88.7 | 41.1 | 88.5 | | 47.4 | 12.5 | 11.6 | 12.6 | 13.3 | 11.7 | 13 | 13.1 | 15.7 | 13.1 | 17.1 |
| PLESHI_03291 | 47.1 | 60.6 | 46.8 | 47.4 | | 14.5 | 13.9 | 13.4 | 16.3 | 12.6 | 13.5 | 11.1 | 14.6 | 13.8 | 17.5 |
| CY652_03435 | 12.4 | 14.9 | 13 | 12.5 | 14.5 | | 15.6 | 12.8 | 13.5 | 42.1 | 13.2 | 17.4 | 13.5 | 12.2 | 14.9 |
| BTO02_30765 | 14.3 | 13.7 | 12.8 | 11.6 | 13.9 | 15.6 | | 14 | 13.7 | 13.3 | 18.4 | 13.3 | 13.1 | 34.9 | 12.8 |
| PSPTO_5204 | 12.1 | 13 | 10.9 | 12.6 | 13.4 | 12.8 | 14 | | 12.3 | 10.7 | 13.2 | 13.2 | 15.3 | 13.4 | 14.6 |
| Dda3937_03232 | 13 | 14.7 | 15.1 | 13.3 | 16.3 | 13.5 | 13.7 | 12.3 | | 13.3 | 17.2 | 14.2 | 14.4 | 17.4 | 18.1 |
| EBL_c11670 | 12.1 | 13.9 | 12 | 11.7 | 12.6 | 42.1 | 13.3 | 10.7 | 13.3 | | 13.7 | 12.3 | 10.7 | 11.6 | 12.8 |
| SAMN04488136_12163 | 13.3 | 13.3 | 12.8 | 13 | 13.5 | 13.2 | 18.4 | 13.2 | 17.2 | 13.7 | | 11.8 | 13.5 | 17.3 | 14.9 |
| ABW06_23985 | 12.6 | 10.9 | 12.4 | 13.1 | 11.1 | 17.4 | 13.3 | 13.2 | 14.2 | 12.3 | 11.8 | | 11.6 | 12.4 | 12.2 |
| SAMN05216206_1427 | 12.3 | 10 | 12 | 15.7 | 14.6 | 13.5 | 13.1 | 15.3 | 14.4 | 10.7 | 13.5 | 11.6 | | 15.9 | 16.3 |
| C4K68_12090Tola_1753 | 12.5 | 14.4 | 13.7 | 13.1 | 13.8 | 12.2 | 34.9 | 13.4 | 17.4 | 11.6 | 17.3 | 12.4 | 15.9 | | 13.8 |
| Tola_1753 | 16.7 | 17.4 | 17 | 17.1 | 17.5 | 14.9 | 12.8 | 14.6 | 18.1 | 12.8 | 14.9 | 12.2 | 16.3 | 13.8 | |

**Supplementary Table S3.** The sequence identity of the C-terminal of TseP and homologs. Sequence identity was calculated using ClustalW web server. Colors are assigned from blue to red indicating low to high sequence similarity, respectively.

| | 1 | 2 | 3 | 4 | 5 | 6 | 7 | 8 | 9 | 10 | 11 | 12 | 13 | 14 | 15 |
|---|---|---|---|---|---|---|---|---|---|---|---|---|---|---|---|
| **TseP** | | 10 | 11.8 | 12.4 | 13.6 | 48.1 | 48.9 | 49.8 | 20.7 | 24.2 | 19.8 | 17.4 | 10.1 | 19.3 | 12.1 |
| NCTC12917_01703 | 10 | | 69.2 | 18.7 | 11.4 | 9.5 | 11.4 | 11.4 | 11.8 | 9.5 | 11.4 | 11.4 | 14.7 | 12.3 | 17.1 |
| WP3S18E02_26330 | 11.8 | 69.2 | | 17.1 | 11.3 | 12.2 | 10.9 | 11.8 | 11.3 | 10.9 | 11.8 | 10 | 14.9 | 11.8 | 15.4 |
| AHA_1849 | 12.4 | 18.7 | 17.1 | | 12.4 | 10.9 | 10.4 | 14 | 11.4 | 11.4 | 10.9 | 12.4 | 15 | 12.4 | 13.5 |
| PLESHI_03291 | 13.6 | 11.4 | 11.3 | 12.4 | | 14.2 | 14 | 14.1 | 13.9 | 13.1 | 13.2 | 13 | 13.6 | 14.6 | 11.6 |
| CY652_03435 | 48.1 | 9.5 | 12.2 | 10.9 | 14.2 | | 46.4 | 49.8 | 21 | 23.6 | 19.8 | 20.6 | 11.4 | 21.5 | 11.6 |
| BTO02_30765 | 48.9 | 11.4 | 10.9 | 10.4 | 14 | 46.4 | | 50.6 | 21.3 | 22.6 | 20.7 | 20.4 | 10.1 | 22.7 | 9.8 |
| PSPTO_5204 | 49.8 | 11.4 | 11.8 | 14 | 14.1 | 49.8 | 50.6 | | 22.4 | 24.6 | 20.3 | 17 | 10.5 | 21 | 10.3 |
| Dda3937_03232 | 20.7 | 11.8 | 11.3 | 11.4 | 13.9 | 21 | 21.3 | 22.4 | | 58.5 | 26.4 | 21.3 | 10.1 | 33.9 | 9.4 |
| EBL_c11670 | 24.2 | 9.5 | 10.9 | 11.4 | 13.1 | 23.6 | 22.6 | 24.6 | 58.5 | | 25.1 | 23.3 | 10.1 | 35.2 | 8.5 |
| SAMN04488136_12163 | 19.8 | 11.4 | 11.8 | 10.9 | 13.2 | 19.8 | 20.7 | 20.3 | 26.4 | 25.1 | | 21.1 | 9.7 | 25.1 | 10.3 |
| ABW06_23985 | 17.4 | 11.4 | 10 | 12.4 | 13 | 20.6 | 20.4 | 17 | 21.3 | 23.3 | 21.1 | | 12.7 | 18.9 | 10.7 |
| SAMN05216206_1427 | 10.1 | 14.7 | 14.9 | 15 | 13.6 | 11.4 | 10.1 | 10.5 | 10.1 | 10.1 | 9.7 | 12.7 | | 11 | 50.4 |
| C4K68_12090Tola_1753 | 19.3 | 12.3 | 11.8 | 12.4 | 14.6 | 21.5 | 22.7 | 21 | 33.9 | 35.2 | 25.1 | 18.9 | 11 | | 11.6 |
| Tola_1753 | 12.1 | 17.1 | 15.4 | 13.5 | 11.6 | 11.6 | 9.8 | 10.3 | 9.4 | 8.5 | 10.3 | 10.7 | 50.4 | 11.6 | |
