## Supplemental Table S4 for "Amidase and Lysozyme Dual Functions in TseP Reveal a New Family of Chimeric Effectors in the Type VI Secretion System"

| Number | Target | Sequence identity | E-value | Scientific Name |
|---|---|---|---|---|
| 1 | AF-K1JFN7-F1-model_v4 EF-hand domain-containing protein | 100 | 0.00E+00 | Aeromonas dhakensis |
| 2 | AF-A0A3S4T080-F1-model_v4 Phage-related lysozyme (Muraminidase) | 88.6 | 0.00E+00 | Aeromonas encheleia |
| 3 | AF-A0A2P1VPN0-F1-model_v4 Uncharacterized protein | 48.2 | 0.00E+00 | Plesiomonas shigelloides |
| 4 | AF-A0A379CLM5-F1-model_v4 Predicted chitinase | 49.3 | 0.00E+00 | Plesiomonas shigelloides |
| 5 | AF-A0A7Z2UK12-F1-model_v4 Uncharacterized protein | 92.1 | 0.00E+00 | Aeromonas hydrophila |
| 6 | AF-A0A2S3XNF5-F1-model_v4 Glyco_hydro_19_cat domain-containing protein | 45.9 | 0.00E+00 | Aeromonas veronii |
| 7 | AF-A0A7D5T027-F1-model_v4 Pesticin domain-containing protein | 47.2 | 0.00E+00 | Aeromonas veronii |
| 8 | AF-A0A0S2SIZ0-F1-model_v4 Uncharacterized protein | 47.1 | 0.00E+00 | Aeromonas schubertii |
| 9 | AF-A0A2H9U6E9-F1-model_v4 Muramidase domain-containing protein | 41.8 | 0.00E+00 | Aeromonas cavernicola |
| 10 | AF-A0A806XLZ8-F1-model_v4 Uncharacterized protein | 93 | 0.00E+00 | Aeromonas hydrophila |
| 11 | AF-A0A327QPZ1-F1-model_v4 Uncharacterized protein DUF3380 | 22.3 | 0.00E+00 | Aeromonas salmonicida |
| 12 | AF-A0A2R7QGU1-F1-model_v4 EF-hand domain-containing protein | 21.7 | 0.00E+00 | Aeromonas sp. HMWF015 |
| 13 | AF-A0A653KIA7-F1-model_v4 EF-hand domain-containing protein | 22.7 | 0.00E+00 | Aeromonas salmonicida |
| 14 | AF-A0A1Z1NNX9-F1-model_v4 EF-hand domain-containing protein | 22.4 | 0.00E+00 | Aeromonas salmonicida |
| 15 | AF-A0A7G1LC66-F1-model_v4 EF-hand domain-containing protein | 21.8 | 0.00E+00 | Aeromonas hydrophila |
| 16 | AF-A0A6M4XMF1-F1-model_v4 EF-hand domain-containing protein | 21.9 | 0.00E+00 | Aeromonas sp. 2692-1 |
| 17 | AF-A0A833JXN3-F1-model_v4 Uncharacterized protein | 22 | 0.00E+00 | Aeromonas veronii |
| 18 | AF-Q0PZF6-F1-model_v4 Uncharacterized protein | 100 | 0.00E+00 | Aeromonas hydrophila |
| 19 | AF-A0A0D5XR60-F1-model_v4 EF-hand domain-containing protein | 21.9 | 0.00E+00 | Pseudomonas chlororaphis |
| 20 | AF-A0A653L6J0-F1-model_v4 Predicted chitinase (Modular protein) | 21.5 | 0.00E+00 | Aeromonas veronii |
| 21 | AF-A0A1J0EL78-F1-model_v4 NlpC/P60 domain-containing protein | 21.4 | 0.00E+00 | Pseudomonas frederiksbergensis |
| 22 | AF-A0A2N1GRP9-F1-model_v4 Peptidase_M23 domain-containing protein | 21.2 | 0.00E+00 | Pseudomonas sp. Choline-02u-1 |
| 23 | AF-A0A5C5Q324-F1-model_v4 Peptidase_M23 domain-containing protein | 20.8 | 0.00E+00 | Pseudomonas extremaustralis |
| 24 | AF-A0A2R7SQD1-F1-model_v4 Uncharacterized protein | 19.2 | 0.00E+00 | Pseudomonas sp. HMWF010 |
| 25 | AF-A0A7Y9WFN0-F1-model_v4 Putative chitinase | 22.6 | 0.00E+00 | Paraburkholderia bryophila |
| 26 | AF-J2YI06-F1-model_v4 Putative chitinase | 22.7 | 0.00E+00 | Pseudomonas sp. GM24 |
| 27 | AF-A0A2S8HLX4-F1-model_v4 Uncharacterized protein | 22.6 | 0.00E+00 | Pseudomonas frederiksbergensis |
| 28 | AF-A0A0Q0AGB6-F1-model_v4 EF hand domain protein | 22.1 | 0.00E+00 | Pseudomonas syringae pv. solidagae |
| 29 | AF-A0A2G0VN88-F1-model_v4 Chitinase | 23 | 0.00E+00 | Pseudomonas sp. ICMP 460 |
| 30 | AF-A0A7Y9VVZ1-F1-model_v4 NlpC/P60 domain-containing protein | 22.3 | 0.00E+00 | Pseudomonas moraviensis |
| 31 | AF-A0A4Y8H7G8-F1-model_v4 EF-hand domain-containing protein | 21.3 | 0.00E+00 | Pseudomonas sp. LAIL14HWK12:I2 |
| 32 | AF-M4WVA5-F1-model_v4 NlpC/P60 domain-containing protein | 21.7 | 0.00E+00 | Pseudomonas sp. ATCC 13867 |
| 33 | AF-W6VFM9-F1-model_v4 Calcium-binding EF-hand-containing protein | 21.1 | 0.00E+00 | Pseudomonas sp. GM30 |
| 34 | AF-A0A0P9QD83-F1-model_v4 EF hand domain protein | 22.3 | 0.00E+00 | Pseudomonas syringae pv. delphinii |

| | | | | |
|---|---|---|---|---|
| 35 | AF-A0A654AQ01-F1-model_v4 Uncharacterized protein | 19 | 0.00E+00 | Pseudomonas sp. 9AZ |
| 36 | AF-A0A3M4VTT8-F1-model_v4 EF-hand domain-containing protein | 22 | 0.00E+00 | Pseudomonas cichorii |
| 37 | AF-A0A5H2XRQ8-F1-model_v4 Uncharacterized protein | 21.6 | 0.00E+00 | Pseudomonas sp. KUIN-1 |
| 38 | AF-A0A5N7JSJ8-F1-model_v4 DUF3380 domain-containing protein | 22.3 | 0.00E+00 | Pseudomonas kitaguniensis |
| 39 | AF-A0A1H7NW51-F1-model_v4 NlpC/P60 domain-containing protein | 19.3 | 0.00E+00 | Atopomonas hussainii |
| 40 | AF-C4LFJ6-F1-model_v4 Uncharacterized protein | 19.8 | 0.00E+00 | Tolumonas auensis DSM 9187 |
| 41 | AF-A0A162AU51-F1-model_v4 Uncharacterized protein | 21.1 | 0.00E+00 | Pseudomonas fluorescens |
| 42 | AF-A0A3M4FRA9-F1-model_v4 Peptidase_M23 domain-containing protein | 21.6 | 0.00E+00 | Pseudomonas syringae pv. berberidis |
| 43 | AF-A0A1G3E770-F1-model_v4 Uncharacterized protein | 19.6 | 0.00E+00 | Pseudomonadales bacterium RIFCSPHIGHO2_02_FULL_60_43 |
| 44 | AF-A0A5C7W1M1-F1-model_v4 NlpC/P60 domain-containing protein | 17.9 | 0.00E+00 | Pseudomonas alcaligenes |
| 45 | AF-A0A653Q6X3-F1-model_v4 Hydrolase_2 domain-containing protein | 19.8 | 0.00E+00 | Pseudomonas sp. 8BK |
| 46 | AF-A0A7G7ETT0-F1-model_v4 Uncharacterized protein | 86.1 | 0.00E+00 | Aeromonas jandaei |
| 47 | AF-A0A4S3YGT8-F1-model_v4 Chitinase | 21.6 | 0.00E+00 | Pseudomonas atacamensis |
| 48 | AF-A0A2S5KSA6-F1-model_v4 EF-hand domain-containing protein | 15.5 | 0.00E+00 | Proteobacteria bacterium 228 |
| 49 | AF-A0A2R7NF23-F1-model_v4 Peptidase_M23 domain-containing protein | 21.8 | 0.00E+00 | Pseudomonas sp. HMWF021 |
| 50 | AF-A0A419N388-F1-model_v4 Uncharacterized protein | 21.4 | 0.00E+00 | Rahnella woolbedingensis |
| 51 | AF-A0A7T8M4B8-F1-model_v4 N-acetylmuramoyl-L-alanine amidase | 16.1 | 0.00E+00 | Entomomonas asaccharolytica |
| 52 | AF-A0A318C115-F1-model_v4 EF-hand domain-containing protein | 16.1 | 0.00E+00 | Pokkaliibacter plantistimulans |
| 53 | AF-A0A7A3AZZ1-F1-model_v4 Lysozyme | 19.9 | 0.00E+00 | Escherichia coli |
| 54 | AF-A0A859WGH6-F1-model_v4 DUF3380 domain-containing protein | 20.7 | 0.00E+00 | Klebsiella grimontii |
| 55 | AF-A0A4C3GAJ5-F1-model_v4 Uncharacterized protein | 20 | 0.00E+00 | Escherichia coli |
| 56 | AF-A0A0J5KRC0-F1-model_v4 Uncharacterized protein | 20.4 | 0.00E+00 | Pluralibacter gergoviae |
| 57 | AF-A0A4R6DPZ0-F1-model_v4 Phage lysozyme-like predicted toxin | 20.3 | 0.00E+00 | Buttiauxella sp. JUb87 |
| 58 | AF-C7C507-F1-model_v4 Lysozyme | 18.8 | 0.00E+00 | Siccibacter turicensis |
| 59 | AF-A0A479JZR6-F1-model_v4 Pesticin domain-containing protein | 20.4 | 0.00E+00 | Escherichia coli |
| 60 | AF-A0A2U3ES87-F1-model_v4 Uncharacterized protein | 17.6 | 0.00E+00 | Enterobacter sp. CGMCC 5087 |
| 61 | AF-I2B6W7-F1-model_v4 EF-hand domain-containing protein | 17.5 | 0.00E+00 | Shimwellia blattae DSM 4481 = NBRC 105725 |
| 62 | AF-A0A3R8YYI2-F1-model_v4 EF-hand domain-containing protein | 17.1 | 0.00E+00 | Enterobacter cloacae |
| 63 | AF-A0A8B3UVL6-F1-model_v4 Phage-related lysozyme (Muraminidase) | 18.4 | 0.00E+00 | Enterobacter hormaechei |
| 64 | AF-A0A3S4JC07-F1-model_v4 Predicted chitinase | 17.4 | 0.00E+00 | Cedecea lapagei |
| 65 | AF-W0BP28-F1-model_v4 Chitinase | 16.4 | 0.00E+00 | Enterobacter ludwigii |
| 66 | AF-A0A2K8W0N2-F1-model_v4 Uncharacterized protein | 16.7 | 0.00E+00 | Dickeya solani RNS 08.23.3.1.A |
| 67 | AF-I2B6V6-F1-model_v4 EF-hand domain-containing protein | 17.5 | 0.00E+00 | Shimwellia blattae DSM 4481 = NBRC 105725 |
| 68 | AF-I2B6W3-F1-model_v4 EF-hand domain-containing protein | 17.3 | 0.00E+00 | Shimwellia blattae DSM 4481 = NBRC 105725 |
| 69 | AF-A0A5F0JQZ1-F1-model_v4 EF-hand domain-containing protein | 17.6 | 0.00E+00 | Enterobacter sp. A11 |

| | | | | |
|---|---|---|---|---|
| 70 | AF-A0A3N4P755-F1-model_v4 EF-hand domain-containing protein | 17.4 | 0.00E+00 | Pantoea sp. RIT388 |
| 71 | AF-W0BUM4-F1-model_v4 LYZ2 domain-containing protein | 17.8 | 0.00E+00 | Enterobacter ludwigii |
| 72 | AF-A0A7T8M297-F1-model_v4 Peptidoglycan DD-metalloendopeptidase family protein | 16.4 | 0.00E+00 | Entomomonas asaccharolytica |
| 73 | AF-A0A379SC66-F1-model_v4 Phage-related lysozyme (Muraminidase) | 18.2 | 0.00E+00 | Salmonella enterica |
| 74 | AF-A0A3S7DM35-F1-model_v4 Peptidase_M23 domain-containing protein | 18.6 | 0.00E+00 | Aeromonas sp. ASNIH4 |
| 75 | AF-A0A2H4ZHK7-F1-model_v4 Phage-encoded peptidoglycan binding protein | 16.4 | 0.00E+00 | Raoultella ornithinolytica |
| 76 | AF-A0A5C7BYA2-F1-model_v4 Uncharacterized protein | 18.3 | 0.00E+00 | Serratia marcescens |
| 77 | AF-A0A433SF78-F1-model_v4 N-acetylmuramoyl-L-alanine amidase | 16.6 | 0.00E+00 | Saezia sanguinis |
| 78 | AF-W1J536-F1-model_v4 Muramidase domain-containing protein | 17.8 | 0.00E+00 | Xenorhabdus szentirmaii DSM 16338 |
| 79 | AF-A0A3T0ZYT9-F1-model_v4 Peptidase_M23 domain-containing protein | 18.7 | 0.00E+00 | Aeromonas hydrophila |
| 80 | AF-A0A0A0CQD6-F1-model_v4 EF-hand domain-containing protein | 17.2 | 0.00E+00 | Photorhabdus luminescens |
| 81 | AF-A0A7T8THQ7-F1-model_v4 CHAP domain-containing protein | 18.9 | 0.00E+00 | Klebsiella pneumoniae |
| 82 | AF-A0A6G6JN88-F1-model_v4 AAA_lid_9 domain-containing protein | 17 | 0.00E+00 | Pantoea stewartii |
| 83 | AF-A0A0U1KJV7-F1-model_v4 Phage-related lysozyme (Muraminidase) | 15.2 | 0.00E+00 | Yersinia mollaretii |
| 84 | AF-A0A4R6E4I7-F1-model_v4 EF-hand domain-containing protein | 17.9 | 0.00E+00 | Buttiauxella sp. JUb87 |
| 85 | AF-A0A2K8QIQ6-F1-model_v4 Uncharacterized protein | 18.5 | 0.00E+00 | Dickeya fangzhongdai |
| 86 | AF-A0A379T132-F1-model_v4 Lytic enzyme | 16.4 | 0.00E+00 | Salmonella enterica subsp. arizonae |
| 87 | AF-Q58PV0-F1-model_v4 Uncharacterized protein | 16.7 | 0.00E+00 | Erwinia amylovora |
| 88 | AF-A0A0U1HZC6-F1-model_v4 Uncharacterized protein | 16.1 | 0.00E+00 | Yersinia mollaretii |
| 89 | AF-A0A7G5CV67-F1-model_v4 Uncharacterized protein | 17.9 | 0.00E+00 | Ewingella americana |
| 90 | AF-A0A377N8F4-F1-model_v4 Phage-related lysozyme (Muraminidase) | 18.3 | 0.00E+00 | Ewingella americana |
| 91 | AF-A0A085G4I9-F1-model_v4 EF-hand domain-containing protein | 17.9 | 0.00E+00 | Ewingella americana ATCC 33852 |
| 92 | AF-A0A4R1NF17-F1-model_v4 Putative chitinase | 17.1 | 0.00E+00 | Sodalis ligni |
| 93 | AF-A0A1Y0KXH2-F1-model_v4 Pesticin domain-containing protein | 17.5 | 0.00E+00 | Pseudomonas sp. M30-35 |
| 94 | AF-A0A7R8RD58-F1-model_v4 EF-hand domain-containing protein | 18.6 | 0.00E+00 | Enterobacter cancerogenus |
| 95 | AF-A4JG68-F1-model_v4 PG_binding_3 domain-containing protein | 18.2 | 0.00E+00 | Burkholderia vietnamiensis G4 |
| 96 | AF-A0A0D5J4V0-F1-model_v4 Bacterial SH3 domain protein | 18 | 0.00E+00 | Burkholderia dolosa AU0158 |
| 97 | AF-A0A6I6IKF0-F1-model_v4 Glyco_hydro_19_cat domain-containing protein | 19.2 | 0.00E+00 | Tatumella sp. TA1 |
| 98 | AF-A0A379SD19-F1-model_v4 Predicted chitinase | 18.7 | 0.00E+00 | Salmonella enterica |
| 99 | AF-A0A379QFF5-F1-model_v4 Phage-related lysozyme (Muraminidase) | 18.7 | 0.00E+00 | Salmonella enterica |
| 100 | AF-A0A3G2IKU8-F1-model_v4 EF-hand domain-containing protein | 17.1 | 0.00E+00 | Buttiauxella sp. 3AFRM03 |
| 101 | AF-A0A5J6R8G5-F1-model_v4 Uncharacterized protein | 17.2 | 0.00E+00 | Pseudomonas denitrificans (nom. rej.) |
| 102 | AF-A0A2J9E7X6-F1-model_v4 EF-hand domain-containing protein | 17.5 | 0.00E+00 | Pantoea sp. FDAARGOS_194 |

| | | | | |
|---|---|---|---|---|
| 103 | AF-C9Y3H6-F1-model_v4 EF-hand domain-containing protein | 19.4 | 0.00E+00 | Cronobacter turicensis z3032 |
| 104 | AF-A0A246F5G2-F1-model_v4 Pesticin domain-containing protein | 17.6 | 0.00E+00 | Pseudomonas nitroreducens |
| 105 | AF-A0A1I5NKC8-F1-model_v4 Predicted chitinase | 16.4 | 0.00E+00 | Pseudomonas sagittaria |
| 106 | AF-A0A828S2D7-F1-model_v4 Chitinase class I family protein | 18.5 | 0.00E+00 | Escherichia coli STEC_7v |
| 107 | AF-A0A1G7UVY5-F1-model_v4 Predicted chitinase | 18.3 | 0.00E+00 | Paraburkholderia phenazinium |
| 108 | AF-A0A1C6YXU7-F1-model_v4 Mannosyl-glycoprotein endo-beta-N-acetylglucosaminidase | 20.5 | 0.00E+00 | Hafnia alvei |
| 109 | AF-A0A0U1KI05-F1-model_v4 Putative endolysin | 16.2 | 0.00E+00 | Yersinia mollaretii |
| 110 | AF-A0A7Y8YN20-F1-model_v4 Glycoside hydrolase family 73 protein | 18.1 | 0.00E+00 | Enterobacter bugandensis |
| 111 | AF-A0A702L6W9-F1-model_v4 Uncharacterized protein | 18.7 | 0.00E+00 | Salmonella enterica subsp. salamae |
| 112 | AF-A0A381KQ07-F1-model_v4 Peptidoglycan hydrolase flgJ | 18.3 | 0.00E+00 | Buttiauxella agrestis |
| 113 | AF-A0A1H1LQD9-F1-model_v4 Uncharacterized protein | 16.1 | 0.00E+00 | Pseudomonas oryzae |
| 114 | AF-J3DH08-F1-model_v4 Putative chitinase | 18.7 | 0.00E+00 | Pantoea sp. GM01 |
| 115 | AF-A0A1E4WMW2-F1-model_v4 Uncharacterized protein | 17.7 | 0.00E+00 | Pseudomonas sp. ENNP23 |
| 116 | AF-A0A7Y8YKH4-F1-model_v4 Glycoside hydrolase family 104 protein | 18.5 | 0.00E+00 | Enterobacter bugandensis |
| 117 | AF-A0A2K1Q4G0-F1-model_v4 EF-hand domain-containing protein | 17.6 | 0.00E+00 | Mixta theicola |
| 118 | AF-A0A2A7TB69-F1-model_v4 Uncharacterized protein | 19.5 | 0.00E+00 | Yersinia kristensenii |
| 119 | AF-A0A7Z9CSA0-F1-model_v4 Phage-related lysozyme (Muraminidase) | 15.7 | 0.00E+00 | Raoultella terrigena |
| 120 | AF-A0A379QQR5-F1-model_v4 Phage-related lysozyme (Muraminidase) | 19.9 | 0.00E+00 | Salmonella enterica |
| 121 | AF-A0A0D5IZJ8-F1-model_v4 Bacterial SH3 domain protein | 18 | 0.00E+00 | Burkholderia dolosa AU0158 |
| 122 | AF-A0A1G6IK64-F1-model_v4 Predicted chitinase | 16.3 | 0.00E+00 | Pseudomonas chengduensis |
| 123 | AF-A0A071MRH4-F1-model_v4 Calcium-binding protein | 18.5 | 0.00E+00 | Burkholderia cenocepacia |
| 124 | AF-A0A5J6QZL7-F1-model_v4 Uncharacterized protein | 18.3 | 0.00E+00 | Pseudomonas denitrificans (nom. rej.) |
| 125 | AF-A0A1P9Y714-F1-model_v4 Pesticin domain-containing protein | 19.2 | 0.00E+00 | Burkholderia sp. KK1 |
| 126 | AF-A0A4V2Q2X0-F1-model_v4 Putative chitinase | 17 | 0.00E+00 | Sodalis ligni |
| 127 | AF-A0A8B2FA03-F1-model_v4 Uncharacterized protein | 17.9 | 0.00E+00 | Providencia stuartii |
| 128 | AF-A0A7L5V640-F1-model_v4 Lysozyme | 18.1 | 0.00E+00 | Escherichia coli |
| 129 | AF-A0A1B4BU39-F1-model_v4 Uncharacterized protein | 18.2 | 0.00E+00 | Burkholderia diffusa |
| 130 | AF-A0A7T8M2F1-F1-model_v4 N-acetylmuramoyl-L-alanine amidase | 15.2 | 0.00E+00 | Entomomonas asaccharolytica |
| 131 | AF-A0A2X4WAZ9-F1-model_v4 Phage-related lysozyme (Muraminidase) | 15.9 | 0.00E+00 | Salmonella enterica subsp. arizonae |
| 132 | AF-A0A2A2E430-F1-model_v4 Uncharacterized protein | 18 | 0.00E+00 | Pseudomonas sp. PIC25 |
| 133 | AF-A0A560URU2-F1-model_v4 Cell wall hydrolase | 17.7 | 0.00E+00 | Burkholderia sp. SJZ115 |
| 134 | AF-A0A7Y7WS97-F1-model_v4 Uncharacterized protein | 17.9 | 0.00E+00 | Pseudomonas gingeri |
| 135 | AF-A0A8A8MKJ9-F1-model_v4 Uncharacterized protein | 18.3 | 0.00E+00 | Burkholderia glumae |
| 136 | AF-A0A2S8RTK3-F1-model_v4 EF hand domain-containing protein | 17.5 | 0.00E+00 | Paraburkholderia sp. BL21I4N1 |
| 137 | AF-A0A3M2H916-F1-model_v4 EF-hand domain-containing protein | 16.6 | 0.00E+00 | Pseudomonas sp. AOB-7 |

| | | | | |
|---|---|---|---|---|
| 138 | AF-A0A1R1J8H9-F1-model_v4 Uncharacterized protein | 18.3 | 0.00E+00 | Burkholderia ubonensis |
| 139 | AF-A0A3A6QN54-F1-model_v4 Glyco_hydro_19_cat domain-containing protein | 19.5 | 0.00E+00 | Pseudomonas sp. LS-2 |
| 140 | AF-W0BWA6-F1-model_v4 Uncharacterized protein | 18.5 | 0.00E+00 | Enterobacter ludwigii |
| 141 | AF-A0A423K9E3-F1-model_v4 Uncharacterized protein | 18.3 | 0.00E+00 | Pseudomonas frederiksbergensis |
| 142 | AF-A0A6S5RPI3-F1-model_v4 Pesticin domain-containing protein | 17.2 | 0.00E+00 | Pseudomonas otitidis |
| 143 | AF-A0A391NEB5-F1-model_v4 Uncharacterized protein | 16 | 0.00E+00 | Pseudomonas sp. SCT |
| 144 | AF-A0A1V4LXQ2-F1-model_v4 Uncharacterized protein | 16.8 | 0.00E+00 | Pseudomonas sp. VI4.1 |
| 145 | AF-A0A0D6SA67-F1-model_v4 Uncharacterized protein | 17.6 | 0.00E+00 | Pseudomonas sp. FeS53a |
| 146 | AF-A0A0D9AI65-F1-model_v4 Glyco_hydro_19_cat domain-containing protein | 18 | 0.00E+00 | Stutzerimonas stutzeri |
| 147 | AF-A0A1V9USM0-F1-model_v4 Uncharacterized protein | 17 | 0.00E+00 | Pseudomonas sp. T |
| 148 | AF-A0A3D3M8H8-F1-model_v4 Uncharacterized protein | 18.1 | 0.00E+00 | Pseudomonas sp. |
| 149 | AF-A0A2W5CSX5-F1-model_v4 Muramidase domain-containing protein | 18.9 | 0.00E+00 | Pseudomonas kuykendallii |
| 150 | AF-A0A679GDI2-F1-model_v4 Uncharacterized protein | 17.1 | 0.00E+00 | Pseudomonas otitidis |
| 151 | AF-A0A210XXD3-F1-model_v4 Uncharacterized protein | 17.4 | 0.00E+00 | Stutzerimonas stutzeri |
| 152 | AF-C5AP31-F1-model_v4 EF hand domain-containing protein | 17.7 | 0.00E+00 | Burkholderia glumae BGR1 |
| 153 | AF-A0A8B3VH40-F1-model_v4 Chitinase class I family protein | 18.4 | 0.00E+00 | Burkholderia multivorans |
| 154 | AF-A0A5E7AHE3-F1-model_v4 EF-hand domain-containing protein | 18.3 | 0.00E+00 | Pseudomonas fluorescens |
| 155 | AF-A0A2N7W9Y3-F1-model_v4 Uncharacterized protein | 18.4 | 0.00E+00 | Trinickia soli |
| 156 | AF-A0A5E6ZGV0-F1-model_v4 EF-hand domain-containing protein | 17.5 | 0.00E+00 | Pseudomonas fluorescens |
| 157 | AF-S6JGB7-F1-model_v4 Uncharacterized protein | 16.8 | 0.00E+00 | Stutzerimonas stutzeri B1SMN1 |
| 158 | AF-A0A2T0HPH8-F1-model_v4 Uncharacterized protein | 18.8 | 0.00E+00 | Pseudomonas simiae |
| 159 | AF-A0A024H9M8-F1-model_v4 EF-hand domain-containing protein | 17.6 | 0.00E+00 | Pseudomonas knackmussii B13 |
| 160 | AF-A0A4U3I8H5-F1-model_v4 DUF3380 domain-containing protein | 19.5 | 0.00E+00 | Pseudomonas sp. CFBP13528 |
| 161 | AF-A0A1V9ULV9-F1-model_v4 Uncharacterized protein | 16.9 | 0.00E+00 | Pseudomonas sp. T |
| 162 | AF-A0A1H5NKN3-F1-model_v4 Uncharacterized protein | 19 | 0.00E+00 | Pseudomonas migulae |
| 163 | AF-A0A1G8U540-F1-model_v4 Uncharacterized protein | 18.3 | 0.00E+00 | Pseudomonas indica |
| 164 | AF-A0A1V9UI10-F1-model_v4 EF-hand domain-containing protein | 17.8 | 0.00E+00 | Pseudomonas sp. Bc-h |
| 165 | AF-A0A5E7FBZ7-F1-model_v4 EF-hand domain-containing protein | 17.4 | 0.00E+00 | Pseudomonas fluorescens |
| 166 | AF-A0A7Y1A897-F1-model_v4 EF-hand domain-containing protein | 18.6 | 0.00E+00 | Pseudomonas veronii |
| 167 | AF-B1JAM9-F1-model_v4 EF-hand domain-containing protein | 18.2 | 0.00E+00 | Pseudomonas putida W619 |
| 168 | AF-A0A4R1PK41-F1-model_v4 Phage lysozyme-like predicted toxin | 18.5 | 0.00E+00 | Azotobacter chroococcum |
| 169 | AF-A0A5E7G4Z9-F1-model_v4 Pesticin domain-containing protein | 17.6 | 0.00E+00 | Pseudomonas fluorescens |
| 170 | AF-A0A4R8M1H9-F1-model_v4 Uncharacterized protein | 17.9 | 0.00E+00 | Paraburkholderia rhizosphaerae |
| 171 | AF-A0A1G8IEF5-F1-model_v4 Predicted chitinase | 18.7 | 0.00E+00 | Pseudomonas panipatensis |
| 172 | AF-A0A2N7VP26-F1-model_v4 Uncharacterized protein | 18.6 | 0.00E+00 | Trinickia dabaoshanensis |

| | | | | |
|---|---|---|---|---|
| 173 | AF-A0A0U4P8L8-F1-model_v4 Uncharacterized protein | 17 | 0.00E+00 | Pseudomonas oryzihabitans |
| 174 | AF-A0A4V2KQ09-F1-model_v4 Uncharacterized protein | 18.6 | 0.00E+00 | Azotobacter chroococcum subsp. isscasi |
| 175 | AF-A0A2X4WTC5-F1-model_v4 Phage-related lysozyme (Muraminidase) | 16.1 | 0.00E+00 | Salmonella enterica subsp. arizonae |
| 176 | AF-A0A1H2B4C6-F1-model_v4 Pesticin domain-containing protein | 17.6 | 0.00E+00 | Pseudomonas umsongensis |
| 177 | AF-A0A1H1X442-F1-model_v4 Predicted chitinase | 17.9 | 0.00E+00 | Pseudomonas umsongensis |
| 178 | AF-A0A2Y9U1D3-F1-model_v4 Uncharacterized protein | 15.9 | 0.00E+00 | Limnobaculum parvum |
| 179 | AF-A0A3T0SBJ4-F1-model_v4 Uncharacterized protein | 14.9 | 0.00E+00 | Pseudomonadaceae bacterium SI-3 |
| 180 | AF-E5B1N7-F1-model_v4 Uncharacterized protein | 15.4 | 0.00E+00 | Erwinia amylovora ATCC BAA-2158 |
| 181 | AF-A0A7Y0ZFU7-F1-model_v4 Uncharacterized protein | 18.8 | 0.00E+00 | Pseudomonas sp. WS 5010 |
| 182 | AF-A0A6P2ZV10-F1-model_v4 Calcium-binding protein | 17.3 | 0.00E+00 | Burkholderia contaminans |
| 183 | AF-A0A5E6SYA6-F1-model_v4 Uncharacterized protein | 17.9 | 0.00E+00 | Pseudomonas fluorescens |
| 184 | AF-A0A5E6XXC3-F1-model_v4 Uncharacterized protein | 16.9 | 0.00E+00 | Pseudomonas fluorescens |
| 185 | AF-A0A0S8ZK03-F1-model_v4 Uncharacterized protein | 17.9 | 0.00E+00 | Pseudomonas sp. Leaf48 |
| 186 | AF-A0A3G2IG66-F1-model_v4 EF-hand domain-containing protein | 17.8 | 0.00E+00 | Buttiauxella sp. 3AFRM03 |
| 187 | AF-A0A329I0V7-F1-model_v4 SH3b domain-containing protein | 15.9 | 0.00E+00 | Pseudomonas sp. RIT 411 |
| 188 | AF-A0A379DT67-F1-model_v4 Predicted chitinase | 15.3 | 0.00E+00 | Pragia fontium |
| 189 | AF-A0A2R7PQD8-F1-model_v4 Uncharacterized protein | 20.8 | 0.00E+00 | Aeromonas sp. HMWF014 |
| 190 | AF-A0A6J4EDA1-F1-model_v4 Uncharacterized protein | 18 | 0.00E+00 | Pseudomonas tohonis |
| 191 | AF-A0A7U6E5V9-F1-model_v4 Lytic transglycosylase domain-containing protein | 17.3 | 0.00E+00 | Burkholderia multivorans |
| 192 | AF-A0A4Q1CRB3-F1-model_v4 EF-hand domain-containing protein | 17.4 | 0.00E+00 | Stenotrophomonas sp. MA5 |
| 193 | AF-A0A653B4F2-F1-model_v4 Predicted chitinase (Modular protein) | 14.4 | 0.00E+00 | Pseudomonas oleovorans |
| 194 | AF-A0A1B8VBW7-F1-model_v4 Uncharacterized protein | 17.8 | 0.00E+00 | Pseudomonas sp. AU11447 |
| 195 | AF-A0A242ND96-F1-model_v4 Uncharacterized protein | 13.6 | 0.00E+00 | Gilliamella apicola |
| 196 | AF-A0A856CLC3-F1-model_v4 Lytic transglycosylase domain-containing protein | 16.5 | 0.00E+00 | Xanthomonas oryzae |
| 197 | AF-A0A1K2HS13-F1-model_v4 Pesticin domain-containing protein | 15.8 | 0.00E+00 | Chitinimonas taiwanensis DSM 18899 |
| 198 | AF-A0A1G7ZEX0-F1-model_v4 Putative peptidoglycan binding domain-containing protein | 13.8 | 0.00E+00 | Vibrio xiamenensis |
| 199 | AF-A0A379SP72-F1-model_v4 Phage-related lysozyme (Muraminidase) | 16.8 | 0.00E+00 | Salmonella enterica subsp. arizonae |
| 200 | AF-A0A4R3I7M2-F1-model_v4 EF-hand domain-containing protein | 14.2 | 0.00E+00 | Reinekea marinisedimentorum |
| 201 | AF-Q8KR50-F1-model_v4 Uncharacterized protein | 16.6 | 0.00E+00 | Escherichia fergusonii |
| 202 | AF-A0A2E9AMK4-F1-model_v4 EF-hand domain-containing protein | 17.9 | 0.00E+00 | Salinisphaera sp. |
| 203 | AF-A0A1T5C8Y0-F1-model_v4 Muramidase domain-containing protein | 13.1 | 0.00E+00 | Luteibacter sp. 22Crub2.1 |
| 204 | AF-A0A118KC83-F1-model_v4 SH3b domain-containing protein | 14.5 | 0.00E+00 | Burkholderia cepacia |
| 205 | AF-A0A0U4CZN5-F1-model_v4 SH3b domain-containing protein | 14.5 | 0.00E+00 | Burkholderia cepacia JBK9 |
| 206 | AF-A0A1B9L7U1-F1-model_v4 Uncharacterized protein | 13.6 | 0.00E+00 | Gilliamella apicola |

| | | | | |
|---|---|---|---|---|
| 207 | AF-A0A2G5NA41-F1-model_v4 EF-hand domain-containing protein | 17.2 | 0.00E+00 | Pseudomonas sp. 2822-17 |
| 208 | AF-A0A4P6GBD5-F1-model_v4 EF-hand domain-containing protein | 18.2 | 0.00E+00 | Pseudomonas arsenicoxydans |
| 209 | AF-A0A423ES90-F1-model_v4 EF-hand domain-containing protein | 15.7 | 0.00E+00 | Pseudomonas poae |
| 210 | AF-C9QG57-F1-model_v4 Uncharacterized protein | 16.7 | 0.00E+00 | Vibrio orientalis CIP 102891 = ATCC 33934 |
| 211 | AF-A0A423FH36-F1-model_v4 Uncharacterized protein | 17.3 | 0.00E+00 | Pseudomonas canadensis |
| 212 | AF-A0A4S3GQW9-F1-model_v4 Cell wall hydrolase | 14.7 | 0.00E+00 | Burkholderia cepacia |
| 213 | AF-A0A484GEZ8-F1-model_v4 N-acetylmuramoyl-L-alanine amidase | 13.1 | 0.00E+00 | Candidatus Schmidhempelia bombi str. Bimp |
| 214 | AF-A0A2I7SIG5-F1-model_v4 EF-hand domain-containing protein | 15.3 | 0.00E+00 | Tamlana carrageenivorans |
| 215 | AF-A0A7U1G1C4-F1-model_v4 EF-hand domain-containing protein | 13.8 | 0.00E+00 | Tamlana sp. s12 |
| 216 | AF-A0A1B9LD41-F1-model_v4 Uncharacterized protein | 14.2 | 0.00E+00 | Gilliamella apicola |
| 217 | AF-A0A1G7IL54-F1-model_v4 Muramidase domain-containing protein | 14.9 | 0.00E+00 | Dyella sp. 333MFSha |
| 218 | AF-A0A841GG37-F1-model_v4 Uncharacterized protein | 16.5 | 0.00E+00 | Tolumonas osonensis |
| 219 | AF-F8H458-F1-model_v4 SH3b domain-containing protein | 15.9 | 0.00E+00 | Stutzerimonas stutzeri ATCC 17588 = LMG 11199 |
| 220 | AF-A0A418YD95-F1-model_v4 Uncharacterized protein | 13.8 | 0.00E+00 | Motilimonas pumila |
| 221 | AF-A0A556RUH6-F1-model_v4 Uncharacterized protein | 12.8 | 0.00E+00 | Gilliamella apicola |
| 222 | AF-A0A1B9KHW1-F1-model_v4 Peptidase_M23 domain-containing protein | 13.6 | 0.00E+00 | Gilliamella apicola |
| 223 | AF-A0A1S2CYG0-F1-model_v4 Lysozyme | 25 | 0.00E+00 | Aeromonas sobria |
| 224 | AF-A0A5C5QKE5-F1-model_v4 Uncharacterized protein | 17.3 | 0.00E+00 | Pseudomonas extremaustralis |
| 225 | AF-A0A286NTW3-F1-model_v4 EF-hand domain-containing protein | 13.2 | 0.00E+00 | Capnocytophaga cynodegmi |
| 226 | AF-A0A2A2DRY7-F1-model_v4 Calcium-binding protein | 17.3 | 0.00E+00 | Pseudomonas sp. PICF141 |
| 227 | AF-Q87UT8-F1-model_v4 EF hand domain protein | 19.8 | 0.00E+00 | Pseudomonas syringae pv. tomato str. DC3000 |
| 228 | AF-A0A3N6UZI5-F1-model_v4 EF-hand domain-containing protein | 22.1 | 0.00E+00 | Aeromonas jandaei |
| 229 | AF-A0A2D5VUQ5-F1-model_v4 Uncharacterized protein | 15.5 | 0.00E+00 | Pseudomonas sp. |
| 230 | AF-A0A5C5VU41-F1-model_v4 N-acetylmuramoyl-L-alanine amidase | 15.7 | 0.00E+00 | Phycisphaerae bacterium RAS1 |
| 231 | AF-A0A7Z1PR97-F1-model_v4 Uncharacterized protein | 20.4 | 0.00E+00 | Salmonella enterica subsp. enterica |
| 232 | AF-A0A6S4ZVB6-F1-model_v4 EF-hand domain-containing protein | 24.3 | 0.00E+00 | Aeromonas caviae |
| 233 | AF-A0A7U9F7W2-F1-model_v4 Uncharacterized protein | 17.8 | 0.00E+00 | Pseudomonas alcaligenes OT 69 |
| 234 | AF-S3IX33-F1-model_v4 EF-hand domain-containing protein | 15.5 | 0.00E+00 | Cedecea davisae DSM 4568 |
| 235 | AF-A0A4Q2QTF0-F1-model_v4 Uncharacterized protein | 17.9 | 0.00E+00 | Enterobacter cloacae complex sp. 743-2DZ2F-22B |
| 236 | AF-A0A379S026-F1-model_v4 EF-hand domain-containing protein | 22.4 | 0.00E+00 | Salmonella enterica subsp. arizonae |
| 237 | AF-A0A1B9LC22-F1-model_v4 Uncharacterized protein | 12.8 | 0.00E+00 | Gilliamella apicola |
| 238 | AF-A0A2N1XA44-F1-model_v4 Uncharacterized protein | 15.2 | 0.00E+00 | Gammaproteobacteria bacterium HGW-Gammaproteobacteria-5 |
| 239 | AF-A0A7X3HDR3-F1-model_v4 Uncharacterized protein | 16.5 | 0.00E+00 | Pseudomonas otitidis |
| 240 | AF-A0A497WKJ2-F1-model_v4 SH3b domain-containing protein | 19.4 | 0.00E+00 | Pseudomonas asplenii |
| 241 | AF-H2J162-F1-model_v4 EF-hand domain-containing protein | 19.8 | 0.00E+00 | Rahnella aquatilis CIP 78.65 = ATCC 33071 |

| | | | | |
|---|---|---|---|---|
| 242 | AF-S9QU80-F1-model_v4 Uncharacterized protein | 12.9 | 0.00E+00 | Cystobacter fuscus DSM 2262 |
| 243 | AF-A0A2N8EL20-F1-model_v4 Uncharacterized protein | 16.3 | 0.00E+00 | Pseudomonas sp. FW305-131 |
| 244 | AF-A0A3S5YGX0-F1-model_v4 Chitinase | 15.4 | 0.00E+00 | Salmonella enterica subsp. arizonae serovar 18:z4,z23:- str. CVM N26626 |
| 245 | AF-A0A379QKR5-F1-model_v4 Phage endolysin | 17.3 | 0.00E+00 | Salmonella enterica |
| 246 | AF-A0A7Y8K5P0-F1-model_v4 Uncharacterized protein | 18.6 | 0.00E+00 | Pseudomonas yamanorum |
| 247 | AF-A0A3W1E7B9-F1-model_v4 Uncharacterized protein | 18.9 | 0.00E+00 | Salmonella enterica |
| 248 | AF-A0A379SG48-F1-model_v4 Phage-related lysozyme (Muraminidase) | 16.7 | 0.00E+00 | Salmonella enterica |
| 249 | AF-A0A3S5YIP0-F1-model_v4 Uncharacterized protein | 15 | 0.00E+00 | Salmonella enterica subsp. arizonae serovar 18:z4,z23:- str. CVM N26626 |
| 250 | AF-A0A7X7I980-F1-model_v4 Peptidoglycan DD-metalloendopeptidase family protein | 12.5 | 0.00E+00 | Fibrobacter sp. |
| 251 | AF-K8B3G2-F1-model_v4 EF-hand domain-containing protein | 20.3 | 0.00E+00 | Cronobacter dublinensis 1210 |
| 252 | AF-B6VN60-F1-model_v4 EF-hand domain-containing protein | 14 | 0.00E+00 | Photorhabdus asymbiotica subsp. asymbiotica ATCC 43949 |
| 253 | AF-A0A2R7Q9V7-F1-model_v4 Uncharacterized protein | 64.9 | 0.00E+00 | Aeromonas sp. HMWF014 |
| 254 | AF-A0A2N7W1U0-F1-model_v4 Uncharacterized protein | 14.6 | 0.00E+00 | Trinickia soli |
| 255 | AF-A0A0U1KGM0-F1-model_v4 Predicted lysozyme (DUF847) | 13.6 | 0.00E+00 | Yersinia mollaretii |
| 256 | AF-A0A1B7HR25-F1-model_v4 Uncharacterized protein | 19.1 | 0.00E+00 | Buttiauxella noackiae ATCC 51607 |
| 257 | AF-A0A6D2GAN1-F1-model_v4 Predicted lysozyme (DUF847) | 18 | 0.00E+00 | Salmonella enterica subsp. salamae |
| 258 | AF-A0A8B4T9D7-F1-model_v4 Phage-related lysozyme (Muraminidase) | 16.8 | 0.00E+00 | Klebsiella pneumoniae |
| 259 | AF-A0A4U2UEC6-F1-model_v4 Pesticin domain-containing protein | 17.7 | 0.00E+00 | Martelella alba |
| 260 | AF-A0A7G6V2K4-F1-model_v4 Uncharacterized protein | 14.4 | 0.00E+00 | Burkholderia cenocepacia |
| 261 | AF-A0A7Z2Z4D7-F1-model_v4 EF-hand domain-containing protein | 87.1 | 0.00E+00 | Aeromonas hydrophila |
| 262 | AF-A0A6P2R1T4-F1-model_v4 Calcium-binding protein | 18 | 0.00E+00 | Burkholderia lata |
| 263 | AF-A0A379S8V1-F1-model_v4 Phage lysozyme protein | 20.3 | 0.00E+00 | Salmonella enterica subsp. arizonae |
| 264 | AF-A0A4Q3ZSL4-F1-model_v4 Hydroxyethylthiazole kinase | 13.1 | 0.00E+00 | Alcaligenaceae bacterium |
| 265 | AF-A0A1Q9QX61-F1-model_v4 Uncharacterized protein | 13.5 | 0.00E+00 | Pseudomonas putida |
| 266 | AF-A0A843SHU8-F1-model_v4 Uncharacterized protein | 12.4 | 0.00E+00 | Rugamonas rivuli |
| 267 | AF-A0A4S0IZA3-F1-model_v4 N-acetylmuramidase family protein | 14.6 | 0.00E+00 | bacterium M00.F.Ca.ET.191.01.1.1 |
| 268 | AF-A0A0Q4MRZ0-F1-model_v4 EF-hand domain-containing protein | 24.5 | 0.00E+00 | Serratia sp. Leaf51 |
| 269 | AF-A0A2G6X112-F1-model_v4 Putative chitinase | 14.2 | 0.00E+00 | Variovorax sp. 54 |
| 270 | AF-G6F3J2-F1-model_v4 Uncharacterized protein | 14.4 | 0.00E+00 | Commensalibacter intestini A911 |
| 271 | AF-A0A0G2ZIR2-F1-model_v4 Muramidase (Phage lysozyme) | 13.1 | 0.00E+00 | Archangium gephyra |
| 272 | AF-A0A515EUQ2-F1-model_v4 M23 family metallopeptidase | 10.5 | 0.00E+00 | Rhodoferax sediminis |
| 273 | AF-A0A4U2U902-F1-model_v4 Uncharacterized protein | 16.8 | 0.00E+00 | Martelella alba |
| 274 | AF-A0A0N9VWZ7-F1-model_v4 Uncharacterized protein | 14.4 | 0.00E+00 | Acinetobacter equi |
| 275 | AF-A0A2Z6U981-F1-model_v4 Uncharacterized protein | 26.4 | 0.00E+00 | Burkholderia vietnamiensis |

| | | | | |
|---|---|---|---|---|
| 276 | AF-A0A379S8W2-F1-model_v4 Phage-related lysozyme (Muraminidase) | 20.2 | 0.00E+00 | Salmonella enterica subsp. arizonae |
| 277 | AF-A0A808K2M3-F1-model_v4 Uncharacterized protein | 17.2 | 0.00E+00 | Paraburkholderia caribensis |
| 278 | AF-A0A158DUE5-F1-model_v4 EF hand domain-containing protein | 22.4 | 0.00E+00 | Caballeronia catudaia |
| 279 | AF-A0A3A9C7C1-F1-model_v4 Uncharacterized protein | 11.8 | 0.00E+00 | bacterium C-53 |
| 280 | AF-A0A2V8Q1G9-F1-model_v4 SH3b domain-containing protein | 18.6 | 0.00E+00 | Acidobacteria bacterium |
| 281 | AF-A0A2V8PTD7-F1-model_v4 Non-specific serine/threonine protein kinase | 16.2 | 0.00E+00 | Acidobacteria bacterium |
| 282 | AF-A0A2V8PFP6-F1-model_v4 SH3b domain-containing protein | 17.4 | 0.00E+00 | Acidobacteria bacterium |
| 283 | AF-A0A2H1XN51-F1-model_v4 SH3b domain-containing protein | 10.5 | 0.00E+00 | Tenacibaculum dicentrarchi |
| 284 | AF-A0A7T9J536-F1-model_v4 Non-specific serine/threonine protein kinase | 17.4 | 0.00E+00 | Acidobacteria bacterium |
| 285 | AF-A0A7W1RAB2-F1-model_v4 SH3 domain-containing protein | 17.4 | 0.00E+00 | Acidobacteria bacterium |
| 286 | AF-A0A0Q7X798-F1-model_v4 Uncharacterized protein | 11.8 | 0.00E+00 | Duganella sp. Root1480D1 |
| 287 | AF-A0A2T4U7M3-F1-model_v4 Uncharacterized protein | 10.5 | 0.00E+00 | Alkalicoccus saliphilus |
| 288 | AF-A0A136KBV8-F1-model_v4 SH3b domain-containing protein | 15.1 | 0.00E+00 | Acidobacteria bacterium OLB17 |
| 289 | AF-A0A740VIV4-F1-model_v4 Uncharacterized protein | 17.9 | 0.00E+00 | Salmonella enterica subsp. enterica serovar 6,7:c:1,5 |
| 290 | AF-A0A7V9GC02-F1-model_v4 SH3 domain-containing protein | 15.5 | 0.00E+00 | Acidobacteria bacterium |
| 291 | AF-A0A7W1ZYT2-F1-model_v4 SH3 domain-containing protein | 16.6 | 0.00E+00 | Acidobacteria bacterium |
| 292 | AF-A0A7Z8PA17-F1-model_v4 SH3 domain-containing protein | 14.4 | 0.00E+00 | Rhodobacteraceae bacterium |
| 293 | AF-A0A060BSK8-F1-model_v4 CAZy families GH73 protein | 13.9 | 0.00E+00 | uncultured Bacillus sp. |
| 294 | AF-A0A2M9NTE7-F1-model_v4 D-alanyl-D-alanine carboxypeptidase | 11.6 | 0.00E+00 | Bacillus sp. mrc49 |
| 295 | AF-A0A6J5J4C6-F1-model_v4 Calcium-binding protein | 25.5 | 0.00E+00 | Burkholderia aenigmatica |
| 296 | AF-A0A7V8VXG5-F1-model_v4 SH3 domain-containing protein | 13.5 | 0.00E+00 | Chloroflexia bacterium |
| 297 | AF-A0A7W1CDA2-F1-model_v4 SH3 domain-containing protein | 11 | 0.00E+00 | Acidobacteria bacterium |
| 298 | AF-A0A0A1FH76-F1-model_v4 Secretion activator protein | 23.5 | 0.00E+00 | Collimonas arenae |
| 299 | AF-A0A2V8RNX3-F1-model_v4 Non-specific serine/threonine protein kinase | 13.3 | 0.00E+00 | Acidobacteria bacterium |
| 300 | AF-A0A2V8QMK2-F1-model_v4 SH3b domain-containing protein | 13.8 | 0.00E+00 | Acidobacteria bacterium |
| 301 | AF-A0A0Q4ND89-F1-model_v4 Lysozyme | 21.5 | 0.00E+00 | Serratia sp. Leaf51 |
| 302 | AF-A3IWE1-F1-model_v4 SH3b domain-containing protein | 14.2 | 0.00E+00 | Crocosphaera chwakensis CCY0110 |
| 303 | AF-T2IMB4-F1-model_v4 Uncharacterized protein | 11.9 | 0.00E+00 | Crocosphaera watsonii WH 0005 |
| 304 | AF-A0A831T5N2-F1-model_v4 Uncharacterized protein | 13.4 | 0.00E+00 | Blastocatellia bacterium |
| 305 | AF-A0A737C652-F1-model_v4 Uncharacterized protein | 14 | 0.00E+00 | Salmonella enterica subsp. arizonae serovar 18:z4,z32:- |
| 306 | AF-M0BGV4-F1-model_v4 N-acetylmuramyl-L-alanine amidase, negative regulator of AmpC, AmpD | 10.8 | 0.00E+00 | Halovivax asiaticus JCM 14624 |
| 307 | AF-A0A126ZF08-F1-model_v4 Uncharacterized protein | 15.5 | 0.00E+00 | Variovorax sp. PAMC 28711 |
| 308 | AF-A0A5C7JTL5-F1-model_v4 SH3b domain-containing protein | 13.7 | 0.00E+00 | Thermomicrobiales bacterium |
| 309 | AF-A0A2V7L197-F1-model_v4 Uncharacterized protein | 11.3 | 0.00E+00 | Gemmatimonadetes bacterium |
| 310 | AF-A0A2G7F7F9-F1-model_v4 Uncharacterized protein | 6.4 | 0.00E+00 | Streptomyces sp. 70 |

| | | | | |
|---|---|---|---|---|
| 311 | AF-A0A3M1SQI4-F1-model_v4 SH3b domain-containing protein | 19.7 | 0.00E+00 | Acidobacteria bacterium |
| 312 | AF-L0M8C9-F1-model_v4 Putative chitinase | 25.9 | 0.00E+00 | Enterobacteriaceae bacterium strain FGI 57 |
| 313 | AF-A0A850H2M4-F1-model_v4 Uncharacterized protein | 14.7 | 0.00E+00 | Altererythrobacter lutimaris |
| 314 | AF-A0A6J4XKJ1-F1-model_v4 SH3b domain-containing protein | 12.7 | 0.00E+00 | Olavius sp. associated proteobacterium Delta 1 |
| 315 | AF-A0A838GH99-F1-model_v4 Protein kinase | 11.7 | 0.00E+00 | Chloroflexia bacterium |
| 316 | AF-A0A6L9IUS5-F1-model_v4 Serine/threonine-protein phosphatase | 10.7 | 0.00E+00 | Chloroflexi bacterium |
| 317 | AF-A0A831WSS0-F1-model_v4 SH3 domain-containing protein | 13.9 | 0.00E+00 | Blastocatellia bacterium |
| 318 | AF-A0A4P6XGS0-F1-model_v4 SH3 domain-containing protein | 16.6 | 0.00E+00 | Sphingomonas sp. AAP5 |
| 319 | AF-A0A6J4H3C9-F1-model_v4 Protein kinase domain-containing protein | 14.1 | 0.00E+00 | uncultured Chloroflexia bacterium |
| 320 | AF-A0A3J4QS28-F1-model_v4 Uncharacterized protein | 16.5 | 0.00E+00 | Salmonella enterica |
| 321 | AF-A0A4P6YLD8-F1-model_v4 Uncharacterized protein | 13.9 | 0.00E+00 | Xanthomonas oryzae pv. oryzae |
| 322 | AF-A0A897MXE4-F1-model_v4 N-acetylmuramoyl-L-alanine amidase | 10.7 | 0.00E+00 | Natranaeroarchaeum sulfidigenes |
| 323 | AF-A0A2N6FQ02-F1-model_v4 EF-hand domain-containing protein | 19.2 | 0.00E+00 | Arcobacter sp. |
| 324 | AF-A0A104ZBZ9-F1-model_v4 Uncharacterized protein | 11.8 | 0.00E+00 | Burkholderia ubonensis |
| 325 | AF-A0A7W4RFV1-F1-model_v4 SH3-like domain-containing protein | 10.2 | 0.00E+00 | Comamonas terrigena |
| 326 | AF-A0A7H8Q6D0-F1-model_v4 SH3 domain-containing protein | 13.7 | 0.00E+00 | Planococcus glaciei |
| 327 | AF-A0A6B1DSB6-F1-model_v4 SH3 domain-containing protein | 12.4 | 0.00E+00 | Caldilineaceae bacterium SB0662 bin 9 |
| 328 | AF-A0A7X1TUM3-F1-model_v4 SH3 domain-containing protein | 9.7 | 0.00E+00 | Betaproteobacteria bacterium |
| 329 | AF-A0A3M1RLR5-F1-model_v4 SH3 domain-containing protein | 16 | 0.00E+00 | Nitrospinae bacterium |
| 330 | AF-A0A497YEF5-F1-model_v4 D-alanyl-D-alanine carboxypeptidase | 16 | 0.00E+00 | Planococcus citreus |
| 331 | AF-A0A7X9J708-F1-model_v4 Protein kinase domain-containing protein | 13.6 | 0.00E+00 | Anaerolineaceae bacterium |
| 332 | AF-A0A840S397-F1-model_v4 Uncharacterized protein | 13.3 | 0.00E+00 | Inhella inkyongensis |
| 333 | AF-A0A1L8P9F6-F1-model_v4 Uncharacterized protein | 11.9 | 0.00E+00 | Alkalibacterium sp. 20 |
| 334 | AF-A0A3B1CD62-F1-model_v4 SH3b domain-containing protein | 9 | 0.00E+00 | hydrothermal vent metagenome |
| 335 | AF-A0A351EVR3-F1-model_v4 LysM domain-containing protein | 13.5 | 0.00E+00 | Acidimicrobiaceae bacterium |
| 336 | AF-A0A6L7XKT4-F1-model_v4 SH3 domain-containing protein | 9 | 0.00E+00 | Rhodospirillales bacterium |
| 337 | AF-A0A7W3P7G9-F1-model_v4 Uncharacterized protein | 11.7 | 0.00E+00 | Microlunatus kandeliicorticis |
| 338 | AF-A0A2W5HBB7-F1-model_v4 EF-hand domain-containing protein | 19.1 | 0.00E+00 | Stenotrophomonas maltophilia |
| 339 | AF-A0A4R5KRP8-F1-model_v4 N-acetylmuramoyl-L-alanine amidase | 12.3 | 0.00E+00 | Paenibacillus piri |
| 340 | AF-A0A838U5I5-F1-model_v4 Serine/threonine protein kinase | 16.4 | 0.00E+00 | Blastocatellia bacterium |
| 341 | AF-A0A2Z6G869-F1-model_v4 Uncharacterized protein | 13.8 | 0.00E+00 | Ferriphaselus amnicola |
| 342 | AF-A0A447ISC7-F1-model_v4 Bacterial SH3 domain protein | 17.7 | 0.00E+00 | Paracoccus haematequi |
| 343 | AF-A0A0A1FJT4-F1-model_v4 Miscellaneous hypothetical | 12 | 0.00E+00 | Collimonas arenae |
| 344 | AF-K1W4A3-F1-model_v4 NLP/P60 protein | 15.4 | 0.00E+00 | Arthrospira platensis C1 |
| 345 | AF-A0A0S8HUB4-F1-model_v4 SH3b domain-containing protein | 15.2 | 0.00E+00 | bacterium SM23_31 |
| 346 | AF-A0A1C6H8P4-F1-model_v4 Bifunctional autolysin | 11.4 | 0.00E+00 | uncultured Clostridium sp. |

| | | | | |
|---|---|---|---|---|
| 347 | AF-A0A6G9W9E4-F1-model_v4 SH3b domain-containing protein | 15.4 | 0.00E+00 | Paracoccus sp. AK26 |
| 348 | AF-T2NAP0-F1-model_v4 SH3 domain protein | 12.1 | 0.00E+00 | Porphyromonas gingivalis JCVI SC001 |
| 349 | AF-A0A6J4UZI8-F1-model_v4 Uncharacterized protein | 11.2 | 0.00E+00 | uncultured Thermomicrobiales bacterium |
| 350 | AF-A0A3D2IPT8-F1-model_v4 Uncharacterized protein | 12.9 | 0.00E+00 | Lachnospiraceae bacterium |
| 351 | AF-A0A644W4E3-F1-model_v4 Uncharacterized protein | 12.5 | 0.00E+00 | bioreactor metagenome |
| 352 | AF-A0A2Z6BXI2-F1-model_v4 SH3b domain-containing protein | 8.3 | 0.00E+00 | Enterococcus faecalis |
| 353 | AF-A0A7C3JTG4-F1-model_v4 SH3b domain-containing protein | 15 | 0.00E+00 | Deltaproteobacteria bacterium |
| 354 | AF-A0A351AYS8-F1-model_v4 SH3 domain-containing protein | 8.1 | 0.00E+00 | Desulfobacterales bacterium |
| 355 | AF-A0A7V9N0Q8-F1-model_v4 Peptidoglycan-binding protein | 11.4 | 0.00E+00 | Alphaproteobacteria bacterium |
| 356 | AF-A0A7C4UVG6-F1-model_v4 Uncharacterized protein | 12.1 | 0.00E+00 | Anaerolineae bacterium |
| 357 | AF-A0A2G3Q2J4-F1-model_v4 Uncharacterized protein | 13.6 | 0.00E+00 | Lachnospiraceae bacterium |
| 358 | AF-A0A356V4C7-F1-model_v4 SH3b domain-containing protein | 13 | 0.00E+00 | Rhodobacteraceae bacterium |
| 359 | AF-A0A5P3MRM2-F1-model_v4 SH3 domain-containing protein | 9.1 | 0.00E+00 | Neisseria animalis |
| 360 | AF-A0A353CUY2-F1-model_v4 Peptide-binding protein | 10.8 | 0.00E+00 | Elusimicrobia bacterium |
| 361 | AF-A0A1Q3RFP0-F1-model_v4 SH3b domain-containing protein | 15.9 | 0.00E+00 | Clostridiales bacterium 38-18 |
| 362 | AF-A0A1I5GLS9-F1-model_v4 NlpC/P60 family protein | 8.2 | 0.00E+00 | Anaerocolumna aminovalerica |
| 363 | AF-I8J3B9-F1-model_v4 N-acetylmuramoyl-L-alanine amidase | 12 | 0.00E+00 | Fictibacillus macauensis ZFHKF-1 |
| 364 | AF-A0A2N6ATY6-F1-model_v4 SH3b domain-containing protein | 12.7 | 0.00E+00 | Clostridiales bacterium |
| 365 | AF-A0A371S019-F1-model_v4 LYZ2 domain-containing protein | 16 | 0.00E+00 | Bacillus sp. HNG |
| 366 | AF-A0A328TZD6-F1-model_v4 Uncharacterized protein | 10.6 | 0.00E+00 | Paenibacillus montanisoli |
| 367 | AF-K2CL96-F1-model_v4 Uncharacterized protein | 13.6 | 0.00E+00 | uncultured bacterium |
| 368 | AF-A0A254N0K3-F1-model_v4 Uncharacterized protein | 14.2 | 0.00E+00 | Pelomonas puraquae |
| 369 | AF-A0A1H8YUL9-F1-model_v4 Uncharacterized protein | 15.5 | 0.00E+00 | Pseudomonas lutea |
| 370 | AF-A0A645CZ87-F1-model_v4 Uncharacterized protein | 10.9 | 0.00E+00 | bioreactor metagenome |
| 371 | AF-A0A1K1QMZ2-F1-model_v4 Cell wall-associated hydrolase, NlpC family | 11.3 | 0.00E+00 | Paenibacillus sp. UNCCL117 |
| 372 | AF-A0A2M8Q2Y1-F1-model_v4 SH3b domain-containing protein | 10 | 0.00E+00 | Phototrophicales bacterium |
| 373 | AF-A0A845GN66-F1-model_v4 SH3 domain-containing protein | 15.2 | 0.00E+00 | Duganella vulcania |
| 374 | AF-A0A0C5JC36-F1-model_v4 Uncharacterized protein | 9.5 | 0.00E+00 | Rugosibacter aromaticivorans |
| 375 | AF-A0A661BKE2-F1-model_v4 SH3b domain-containing protein | 10.7 | 0.00E+00 | Gammaproteobacteria bacterium |
| 376 | AF-A0A2U2BWA7-F1-model_v4 Uncharacterized protein | 12.6 | 0.00E+00 | Marinicauda salina |
| 377 | AF-A0A2R6JR74-F1-model_v4 N-acetylmuramoyl-L-alanine amidase | 12.6 | 0.00E+00 | Halobacteriales archaeon QS_8_69_26 |
| 378 | AF-A0A2G6FTY5-F1-model_v4 SH3b domain-containing protein | 10.8 | 0.00E+00 | Deltaproteobacteria bacterium |
| 379 | AF-A0A352JPJ0-F1-model_v4 Uncharacterized protein | 10 | 0.00E+00 | candidate division Zixibacteria bacterium |
| 380 | AF-A0A1H6GQG3-F1-model_v4 SH3 domain-containing protein | 11.5 | 0.00E+00 | Magnetospirillum fulvum |
| 381 | AF-A0A2V7KDF7-F1-model_v4 Beta_helix domain-containing protein | 11.8 | 0.00E+00 | Gemmatimonadetes bacterium |
| 382 | AF-A0A812KVI3-F1-model_v4 Hypothetical protein | 11.5 | 0.00E+00 | Symbiodinium natans |

| | | | | |
|---|---|---|---|---|
| 383 | AF-A0A1H4N811-F1-model_v4 SH3 domain-containing protein | 13.8 | 0.00E+00 | Rhodobacter sp. 24-YEA-8 |
| 384 | AF-E1ICP5-F1-model_v4 SH3 type 3 domain-containing protein | 11.3 | 0.00E+00 | Oscillochloris trichoides DG-6 |
| 385 | AF-A0A2N2EGM8-F1-model_v4 Uncharacterized protein | 11.2 | 0.00E+00 | Firmicutes bacterium HGW-Firmicutes-1 |
| 386 | AF-A0A4R0YLQ5-F1-model_v4 Curli production assembly/transport component CsgG | 13.9 | 0.00E+00 | Dyella soli |
| 387 | AF-A0A231RR77-F1-model_v4 Hydrolase Nlp/P60 | 12.7 | 0.00E+00 | Cohnella sp. CIP 111063 |
| 388 | AF-A0A0U4E5J8-F1-model_v4 SH3b domain-containing protein | 11.3 | 0.00E+00 | Lentibacillus amyloliquefaciens |
| 389 | AF-F2NYD8-F1-model_v4 SH3b domain-containing protein | 12.2 | 0.00E+00 | Treponema succinifaciens DSM 2489 |
| 390 | AF-S9ZJQ0-F1-model_v4 Uncharacterized protein | 9.7 | 0.00E+00 | Thauera terpenica 58Eu |
| 391 | AF-A0A6I1QG31-F1-model_v4 SH3 domain-containing protein | 13 | 0.00E+00 | Vibrio sp. B1Z05 |
| 392 | AF-A0A7W1DXH2-F1-model_v4 SH3 domain-containing protein | 17.4 | 0.00E+00 | Chloroflexia bacterium |
| 393 | AF-A0A844ZCN4-F1-model_v4 Uncharacterized protein | 14.7 | 0.00E+00 | Parapontixanthobacter aurantiacus |
| 394 | AF-A0A2A8MH91-F1-model_v4 Cell wall-binding protein | 16.1 | 0.00E+00 | Bacillus sp. AFS001701 |
| 395 | AF-A0A4R6RIL8-F1-model_v4 Uncharacterized protein | 11.3 | 0.00E+00 | Aquabacterium commune |
| 396 | AF-A0A813FBF0-F1-model_v4 Hypothetical protein | 14.7 | 0.00E+00 | Polarella glacialis |
| 397 | AF-A0A1H9GNF3-F1-model_v4 Uncharacterized protein | 9.2 | 0.00E+00 | Treponema bryantii |
| 398 | AF-A0A4V2Z7Q4-F1-model_v4 SH3 domain-containing protein | 15.2 | 0.00E+00 | Antarcticimicrobium sediminis |
| 399 | AF-A0A845CU74-F1-model_v4 SH3 domain-containing protein | 10.4 | 0.00E+00 | Caldilineaceae bacterium SB0662 bin 25 |
| 400 | AF-A0A653J1E5-F1-model_v4 Uncharacterized protein | 13.1 | 0.00E+00 | Novosphingobium sp. 9U |
| 401 | AF-A0A7V3UHX2-F1-model_v4 SH3b domain-containing protein | 15.8 | 0.00E+00 | bacterium |
| 402 | AF-A0A3G5HRB2-F1-model_v4 Uncharacterized protein | 11.2 | 0.00E+00 | Propionibacterium acidifaciens |
| 403 | AF-A0A812PD46-F1-model_v4 Hypothetical protein | 12.3 | 0.00E+00 | Symbiodinium sp. CCMP2592 |
| 404 | AF-R0BZZ3-F1-model_v4 SH3b domain-containing protein | 15.7 | 0.00E+00 | Enterocloster bolteae 90A9 |
| 405 | AF-A0A1R1I295-F1-model_v4 Uncharacterized protein | 9.1 | 0.00E+00 | Azonexus hydrophilus |
| 406 | AF-A0A7G1GZS7-F1-model_v4 SH3b domain-containing protein | 7 | 0.00E+00 | Dissulfurispira thermophila |
| 407 | AF-A0A4U8RY23-F1-model_v4 SH3b domain-containing protein | 13.1 | 0.00E+00 | Helicobacter typhlonius |
| 408 | AF-A0A178M552-F1-model_v4 Ligand-binding protein SH3 | 11.3 | 0.00E+00 | Chloroflexus islandicus |
| 409 | AF-A0A348N5I7-F1-model_v4 SH3b domain-containing protein | 12 | 0.00E+00 | Lachnospiraceae bacterium |
| 410 | AF-A0A1S9BVG9-F1-model_v4 SH3b domain-containing protein | 9.5 | 0.00E+00 | Oribacterium sp. C9 |
| 411 | AF-A0A099GBV6-F1-model_v4 Uncharacterized protein | 10.7 | 0.00E+00 | Paracoccus sanguinis |
| 412 | AF-A0A2E4A2J9-F1-model_v4 Uncharacterized protein | 9.6 | 0.00E+00 | Candidatus Marinimicrobia bacterium |
| 413 | AF-A0A7Z8YPF8-F1-model_v4 Chaperone protein DnaJ | 14.1 | 0.00E+00 | Bergeyella zoohelcum |
| 414 | AF-A0A2V9LAW5-F1-model_v4 Uncharacterized protein | 14.8 | 0.00E+00 | Acidobacteria bacterium |
| 415 | AF-A0A7W0S603-F1-model_v4 SH3 domain-containing protein | 12.9 | 0.00E+00 | Chloroflexia bacterium |
| 416 | AF-A0A2V7L188-F1-model_v4 Uncharacterized protein | 8.9 | 0.00E+00 | Gemmatimonadetes bacterium |
| 417 | AF-A0A3M1YD68-F1-model_v4 Uncharacterized protein | 12.2 | 0.00E+00 | Bacteroidetes bacterium |

| | | | | |
|---|---|---|---|---|
| 418 | AF-A0A7S2HAX7-F1-model_v4 Hypothetical protein | 10 | 0.00E+00 | Alexandrium andersonii |
| 419 | AF-A0A4Q3DNQ8-F1-model_v4 SH3 domain-containing protein | 8.6 | 0.00E+00 | Hyphomicrobiales bacterium |
| 420 | AF-A0A6M2BWN1-F1-model_v4 Uncharacterized protein | 11 | 0.00E+00 | Solimonas terrae |
| 421 | AF-A0A2V9FGK1-F1-model_v4 SH3b domain-containing protein | 14.5 | 0.00E+00 | Acidobacteria bacterium |
| 422 | AF-N6YG11-F1-model_v4 Uncharacterized protein | 7.5 | 0.00E+00 | Thauera linaloolentis 47Lol = DSM 12138 |
| 423 | AF-A0A2E7BU14-F1-model_v4 Uncharacterized protein | 13 | 0.00E+00 | Myxococcales bacterium |
| 424 | AF-A0A6L9IUF1-F1-model_v4 SH3 domain-containing protein | 10.5 | 0.00E+00 | Chloroflexi bacterium |
| 425 | AF-A0A2R7L7D9-F1-model_v4 SH3b domain-containing protein | 16.1 | 0.00E+00 | Caulobacter sp. HMWF025 |
| 426 | AF-A0A6M0IDQ9-F1-model_v4 DnaJ domain-containing protein | 8.9 | 0.00E+00 | Spirosoma agri |
| 427 | AF-A0A3D1E9P2-F1-model_v4 Ion channel protein | 11.3 | 0.00E+00 | Flavobacteriaceae bacterium |
| 428 | AF-W7ZMW5-F1-model_v4 SH3b domain-containing protein | 9.4 | 0.00E+00 | Bacillus sp. JCM 19047 |
| 429 | AF-A0A329KSB1-F1-model_v4 Uncharacterized protein | 7.7 | 0.00E+00 | Paenibacillus sp. YN15 |
| 430 | AF-A0A1I5UL46-F1-model_v4 SH3 domain-containing protein | 16.3 | 0.00E+00 | Lachnospiraceae bacterium XBB1006 |
| 431 | AF-A0A3S1AI16-F1-model_v4 PPM-type phosphatase domain-containing protein | 15.1 | 0.00E+00 | Calothrix desertica PCC 7102 |
| 432 | AF-A0A7S1B0F5-F1-model_v4 Hypothetical protein | 13.9 | 0.00E+00 | Noctiluca scintillans |
| 433 | AF-A0A0F9KHA1-F1-model_v4 SH3b domain-containing protein | 8.6 | 0.00E+00 | marine sediment metagenome |
| 434 | AF-A0A1Q9LKR9-F1-model_v4 Uncharacterized protein | 8.9 | 0.00E+00 | Actinokineospora bangkokensis |
| 435 | AF-A0A3D5T073-F1-model_v4 Uncharacterized protein | 8.6 | 0.00E+00 | Rhodocyclaceae bacterium |
| 436 | AF-A0A3M1TG86-F1-model_v4 SH3 domain-containing protein | 9.8 | 0.00E+00 | Cyanobacteria bacterium J055 |
| 437 | AF-A0A7Y8STV0-F1-model_v4 SH3 domain-containing protein | 11.6 | 0.00E+00 | Hydrogenophilaceae bacterium |
| 438 | AF-A0A1Y3C7M3-F1-model_v4 Uncharacterized protein | 10.3 | 0.00E+00 | Acinetobacter sp. ANC 3903 |
| 439 | AF-A0A0R1WXL1-F1-model_v4 N-acetylmuramoyl-L-alanine amidase | 11.4 | 0.00E+00 | Ligilactobacillus hayakitensis DSM 18933 = JCM 14209 |
| 440 | AF-A0A7S2IYD3-F1-model_v4 Hypothetical protein | 15.2 | 0.00E+00 | Alexandrium andersonii |
| 441 | AF-A0A2M8RTE1-F1-model_v4 Uncharacterized protein | 12 | 0.00E+00 | Caviibacterium pharyngocola |
| 442 | AF-A0A3D2JJL4-F1-model_v4 GH18 domain-containing protein | 9 | 0.00E+00 | Lachnospiraceae bacterium |
| 443 | AF-A0A836TBG1-F1-model_v4 NUDIX domain-containing protein | 13.7 | 0.00E+00 | Chromatiaceae bacterium |
| 444 | AF-A0A5R9GPF8-F1-model_v4 SH3b domain-containing protein | 11.7 | 0.00E+00 | Mariprofundus erugo |
| 445 | AF-A0A2R6G7A9-F1-model_v4 Uncharacterized protein | 8.9 | 0.00E+00 | Halobacteriales archaeon QS_1_68_20 |
| 446 | AF-A0A840SGW6-F1-model_v4 Uncharacterized protein YgiM (DUF1202 family) | 15.2 | 0.00E+00 | Treponema rectale |
| 447 | AF-A0A160TPJ9-F1-model_v4 Uncharacterized protein | 14.1 | 0.00E+00 | hydrothermal vent metagenome |
| 448 | AF-A0A511AS69-F1-model_v4 Uncharacterized protein | 9.3 | 0.00E+00 | Alkalibacterium kapii |
| 449 | AF-A0A564WD70-F1-model_v4 Uncharacterized protein | 7.6 | 0.00E+00 | Candidatus Defluviicoccus seviourii |
| 450 | AF-A0A1M5SGA6-F1-model_v4 SH3b domain-containing protein | 14.6 | 0.00E+00 | Tepidibacter thalassicus DSM 15285 |
| 451 | AF-A0A7S1PID9-F1-model_v4 Hypothetical protein | 12.8 | 0.00E+00 | Alexandrium catenella |
| 452 | AF-M6G566-F1-model_v4 SH3b domain-containing protein | 26.7 | 0.00E+00 | Leptospira interrogans str. 2006001854 |

| | | | | |
|---|---|---|---|---|
| 453 | AF-A0A1Q7TH32-F1-model_v4 SH3b domain-containing protein | 12.1 | 0.00E+00 | Gemmatimonadetes bacterium 13_1_20CM_4_66_11 |
| 454 | AF-A0A0P9EBU6-F1-model_v4 SH3-like domain-containing protein | 8.1 | 0.00E+00 | Thiohalorhabdus denitrificans |
| 455 | AF-V5FKU3-F1-model_v4 Uncharacterized protein | 12.7 | 0.00E+00 | Vibrio halioticoli NBRC 102217 |
| 456 | AF-A0A7S4VH77-F1-model_v4 Hypothetical protein | 9.6 | 0.00E+00 | Alexandrium monilatum |
| 457 | AF-A0A4V6HYJ5-F1-model_v4 SH3b domain-containing protein | 14.2 | 0.00E+00 | Helicobacter sp. MIT 05-5293 |
| 458 | AF-S3K1G0-F1-model_v4 Uncharacterized protein | 13.5 | 0.00E+00 | Treponema maltophilum ATCC 51939 |
| 459 | AF-A0A520E178-F1-model_v4 Ligand-binding protein SH3 | 10.5 | 0.00E+00 | Variovorax sp. |
| 460 | AF-A0A3R6TU91-F1-model_v4 Uncharacterized protein | 9.5 | 0.00E+00 | Clostridium sp. AF12-19 |
| 461 | AF-A0A844WUA6-F1-model_v4 Uncharacterized protein | 10.5 | 0.00E+00 | Gilliamella sp. Pas-s27 |
| 462 | AF-A0A812ULJ6-F1-model_v4 STOML2 protein | 14.2 | 0.00E+00 | Symbiodinium sp. CCMP2592 |
| 463 | AF-A0A812YRP3-F1-model_v4 Hypothetical protein | 11.5 | 0.00E+00 | Symbiodinium sp. CCMP2456 |
| 464 | AF-A0A7S1AGW5-F1-model_v4 Hypothetical protein | 10.3 | 0.00E+00 | Noctiluca scintillans |
| 465 | AF-A0A4Q3DTW1-F1-model_v4 SH3 domain-containing protein | 14.4 | 0.00E+00 | Hyphomicrobiales bacterium |
| 466 | AF-A0A7T9ECQ4-F1-model_v4 Uncharacterized protein | 15.1 | 0.00E+00 | Fibrobacteres bacterium |
| 467 | AF-A0A346QWK2-F1-model_v4 Nlp p60 protein | 13.6 | 0.00E+00 | uncultured bacterium |
| 468 | AF-A0A6B0H4E0-F1-model_v4 SH3 domain-containing protein | 10.4 | 0.00E+00 | Saccharibacillus sp. WB 17 |
| 469 | AF-A0A2G6FGN6-F1-model_v4 SH3b domain-containing protein | 11 | 0.00E+00 | Desulfobacterales bacterium |
| 470 | AF-A0A4R0PKP3-F1-model_v4 Type VI secretion system protein TssA | 15.6 | 0.00E+00 | Pseudomonas sp. IC_126 |
| 471 | AF-M1YM91-F1-model_v4 Uncharacterized protein | 11.6 | 0.00E+00 | Nitrospina gracilis 3/211 |
| 472 | AF-A0A2A7UUW6-F1-model_v4 Peptide-binding protein | 12.2 | 0.00E+00 | Comamonas terrigena |
| 473 | AF-A0A4R2VRW4-F1-model_v4 Uncharacterized protein | 14 | 0.00E+00 | Niastella sp. CF465 |
| 474 | AF-A0A534S358-F1-model_v4 SH3 domain-containing protein | 12.7 | 0.00E+00 | Deltaproteobacteria bacterium |
| 475 | AF-A0A7Y2SSF3-F1-model_v4 SH3 domain-containing protein | 14.1 | 0.00E+00 | Chloroflexales bacterium ZM16-3 |
| 476 | AF-R6CA66-F1-model_v4 Uncharacterized protein | 10.3 | 0.00E+00 | Clostridium sp. CAG:510 |
| 477 | AF-A0A838UQH2-F1-model_v4 Uncharacterized protein | 11.8 | 0.00E+00 | Ktedonobacterales bacterium |
| 478 | AF-E8N102-F1-model_v4 Uncharacterized protein | 14.9 | 0.00E+00 | Anaerolinea thermophila UNI-1 |
| 479 | AF-A0A7S1MU02-F1-model_v4 Hypothetical protein | 9.2 | 0.00E+00 | Alexandrium catenella |
| 480 | AF-A0A3D2GZT2-F1-model_v4 Uncharacterized protein | 10.5 | 0.00E+00 | Clostridiales bacterium |
| 481 | AF-A0A812L040-F1-model_v4 STOML2 protein | 10.8 | 0.00E+00 | Symbiodinium natans |
| 482 | AF-A0A849MHA0-F1-model_v4 SH3 domain-containing protein | 11.4 | 0.00E+00 | Proteobacteria bacterium |
| 483 | AF-A0A5N7J2I3-F1-model_v4 SH3 domain-containing protein | 11.9 | 0.00E+00 | Clostridium estertheticum |
| 484 | AF-A0A849YZP5-F1-model_v4 SH3 domain-containing protein | 9.9 | 0.00E+00 | Polyangiaceae bacterium |
| 485 | AF-A0A1Z9SEX5-F1-model_v4 NlpC/P60 domain-containing protein | 10.7 | 0.00E+00 | Rhodospirillaceae bacterium TMED256 |
| 486 | AF-E7GEP5-F1-model_v4 Uncharacterized protein | 7.5 | 0.00E+00 | Coprobacillus cateniformis |
| 487 | AF-A0A7C2PFK9-F1-model_v4 SH3 domain-containing protein | 10.1 | 0.00E+00 | Spirochaetes bacterium |
| 488 | AF-A0A7S1RQP7-F1-model_v4 Hypothetical protein | 10 | 0.00E+00 | Alexandrium catenella |

| | | | | |
|---|---|---|---|---|
| 489 | AF-A3IYE7-F1-model_v4 SH3b domain-containing protein | 7.1 | 0.00E+00 | Crocosphaera chwakensis CCY0110 |
| 490 | AF-A0A3N1HNW9-F1-model_v4 Uncharacterized protein | 14 | 0.00E+00 | Streptomyces sp. PanSC9 |
| 491 | AF-A0A0D5A037-F1-model_v4 SH3 type 3 domain-containing protein | 10.9 | 0.00E+00 | Ochrobactrum sp. LM19 |
| 492 | AF-A0A7C9P2P3-F1-model_v4 Uncharacterized protein | 11.5 | 0.00E+00 | Sulfuriferula multivorans |
| 493 | AF-A0A3M1C5S4-F1-model_v4 Uncharacterized protein | 9.5 | 0.00E+00 | Bdellovibrio sp. |
| 494 | AF-G4CVM6-F1-model_v4 SH3b domain-containing protein | 12.7 | 0.00E+00 | Cutibacterium avidum ATCC 25577 |
| 495 | AF-A0A2T5K2L6-F1-model_v4 SH3-like domain-containing protein | 6 | 0.00E+00 | Nitrosospira sp. Nsp2 |
| 496 | AF-A0A0Q7I513-F1-model_v4 Ligand-binding protein SH3 | 10.9 | 0.00E+00 | Variovorax sp. Root434 |
| 497 | AF-A0A813LWA7-F1-model_v4 Hypothetical protein | 7 | 0.00E+00 | Polarella glacialis |
| 498 | AF-D3EDZ5-F1-model_v4 NLP/P60 protein | 13.2 | 0.00E+00 | Paenibacillus sp. Y412MC10 |
| 499 | AF-A0A536TWX5-F1-model_v4 SH3b domain-containing protein | 10.9 | 0.00E+00 | Betaproteobacteria bacterium |
| 500 | AF-A0A4Q0U6Y5-F1-model_v4 NlpC/P60 domain-containing protein | 9.8 | 0.00E+00 | Candidatus Amulumruptor caecigallinarius |
| 501 | AF-A0A7W1TN07-F1-model_v4 C40 family peptidase | 9.2 | 0.00E+00 | Herpetosiphonaceae bacterium |
| 502 | AF-A0A1Q6SRG6-F1-model_v4 Uncharacterized protein | 14.7 | 0.00E+00 | Roseburia sp. CAG:10041_57 |
| 503 | AF-A0A6A5BXB1-F1-model_v4 SH3b domain-containing protein | 7.6 | 0.00E+00 | Naegleria fowleri |
| 504 | AF-A0A7S4T4T8-F1-model_v4 Hypothetical protein | 12.3 | 0.00E+00 | Alexandrium monilatum |
| 505 | AF-A0A1Q8QDJ4-F1-model_v4 Uncharacterized protein | 16.5 | 0.00E+00 | Desulfovibrio sp. DV |
| 506 | AF-A0A3E1DID4-F1-model_v4 Uncharacterized protein | 14.6 | 0.00E+00 | Candidatus Nitrotoga sp. SPKER |
| 507 | AF-A0A812W9S7-F1-model_v4 STOML2 protein | 11 | 0.00E+00 | Symbiodinium sp. CCMP2456 |
| 508 | AF-I4B0Y6-F1-model_v4 Uncharacterized protein | 7.3 | 0.00E+00 | Turneriella parva DSM 21527 |
| 509 | AF-A0A661WR21-F1-model_v4 SH3b domain-containing protein | 12.4 | 0.00E+00 | Chloroflexi bacterium |
| 510 | AF-J8GGJ5-F1-model_v4 SH3b domain-containing protein | 9.8 | 0.00E+00 | Bacillus cereus MSX-D12 |
| 511 | AF-A0A4V6I468-F1-model_v4 SH3b domain-containing protein | 11.6 | 0.00E+00 | Helicobacter japonicus |
| 512 | AF-A0A7S1QHN2-F1-model_v4 Hypothetical protein | 9.2 | 0.00E+00 | Alexandrium catenella |
| 513 | AF-A0A2E9AN17-F1-model_v4 SH3b domain-containing protein | 9.6 | 0.00E+00 | Salinisphaera sp. |
| 514 | AF-C0EHW9-F1-model_v4 Glycosyl hydrolase family 25 | 9.4 | 0.00E+00 | [Clostridium] methylpentosum DSM 5476 |
| 515 | AF-A0A1G9FK35-F1-model_v4 Cell wall-associated hydrolase, NlpC family | 13 | 0.00E+00 | Paenibacillus typhae |
| 516 | AF-B6IWB1-F1-model_v4 SH3b domain-containing protein | 12 | 0.00E+00 | Rhodospirillum centenum SW |
| 517 | AF-A0A268RVC0-F1-model_v4 Uncharacterized protein | 9.9 | 0.00E+00 | Alkalihalobacillus clausii |
| 518 | AF-A0A3N9UWN0-F1-model_v4 SH3b domain-containing protein | 12.5 | 0.00E+00 | Methanothrix sp. |
| 519 | AF-E1ZIE4-F1-model_v4 SH3b domain-containing protein | 14.5 | 0.00E+00 | Chlorella variabilis |
| 520 | AF-A0A6B4KH23-F1-model_v4 SH3 domain-containing protein | 11.1 | 0.00E+00 | Clostridium botulinum |
| 521 | AF-A0A1W6MZG0-F1-model_v4 SH3b domain-containing protein | 8 | 0.00E+00 | Methylocystis bryophila |
| 522 | AF-A0A2M8U5C8-F1-model_v4 Uncharacterized protein | 10.6 | 0.00E+00 | Ferrovibrio sp. |
| 523 | AF-A0A4Q5RH70-F1-model_v4 SH3 domain-containing protein | 10.7 | 0.00E+00 | Cytophagaceae bacterium |
| 524 | AF-A0A7X0E3Q0-F1-model_v4 Uncharacterized protein YraI | 15.8 | 0.00E+00 | Rhodanobacter sp. MP1X3 |

| | | | | |
|---|---|---|---|---|
| 525 | AF-A0A7X8F336-F1-model_v4 Uncharacterized protein | 14.7 | 0.00E+00 | Treponema sp. |
| 526 | AF-A0A1X7LW17-F1-model_v4 Cell wall-associated hydrolase, NlpC family | 11.1 | 0.00E+00 | Paenibacillus aquistagni |
| 527 | AF-A0A7S0B6U2-F1-model_v4 Hypothetical protein | 11.4 | 0.00E+00 | Pyrodinium bahamense |
| 528 | AF-R7MCW5-F1-model_v4 Dockerin type I repeat protein | 9.3 | 0.00E+00 | Clostridium sp. CAG:628 |
| 529 | AF-A0A812H8V8-F1-model_v4 Ttc4 protein | 13.3 | 0.00E+00 | Symbiodinium sp. CCMP2456 |
| 530 | AF-A0A364JZ10-F1-model_v4 Uncharacterized protein YraI | 12.5 | 0.00E+00 | Falsochrobactrum ovis |
| 531 | AF-A0A0M9E8P7-F1-model_v4 SH3, type 3 domain protein | 13.4 | 0.00E+00 | Candidatus Magnetomorum sp. HK-1 |
| 532 | AF-A0A3P3RA12-F1-model_v4 Uncharacterized protein | 12.4 | 0.00E+00 | Halomarina oriensis |
| 533 | AF-A0A377JLF0-F1-model_v4 Bacterial SH3 domain | 11.6 | 0.00E+00 | Helicobacter canis |
| 534 | AF-A0A1S2UXG5-F1-model_v4 SH3b domain-containing protein | 10.5 | 0.00E+00 | Pseudomonas costantinii |
| 535 | AF-A0A084SGA4-F1-model_v4 Uncharacterized protein | 10.8 | 0.00E+00 | Archangium violaceum Cb vi76 |
| 536 | AF-R7QW32-F1-model_v4 Bacterial SH3 domain | 10.9 | 0.00E+00 | Roseburia sp. CAG:182 |
| 537 | AF-A0A2N6BYY7-F1-model_v4 SH3b domain-containing protein | 10.8 | 0.00E+00 | Desulfobulbaceae bacterium |
| 538 | AF-A0A2H5ZPP9-F1-model_v4 SH3b domain-containing protein | 13 | 0.00E+00 | bacterium HR30 |
| 539 | AF-A0A812SE78-F1-model_v4 Pkd2 protein | 9.6 | 0.00E+00 | Symbiodinium pilosum |
| 540 | AF-A0A1A9NLG7-F1-model_v4 Uncharacterized protein | 14.6 | 0.00E+00 | Methylobacillus sp. MM3 |
| 541 | AF-A0A855KWT0-F1-model_v4 Enterotoxin | 9.9 | 0.00E+00 | Bacillus sp. AKBS9 |
| 542 | AF-A0A836WVM9-F1-model_v4 SH3 domain-containing protein | 13.2 | 0.00E+00 | Thiotrichaceae bacterium |
| 543 | AF-A0A2P1VC11-F1-model_v4 Ligand-binding protein SH3 | 9.7 | 0.00E+00 | Variovorax sp. PMC12 |
| 544 | AF-A0A6B1DWR4-F1-model_v4 SH3 domain-containing protein | 10.7 | 0.00E+00 | Caldilineaceae bacterium SB0662_bin_9 |
| 545 | AF-A0A3N4N2H2-F1-model_v4 GIY-YIG nuclease family protein | 15 | 0.00E+00 | Neisseria sp. 10009 |
| 546 | AF-A0A2V9V4W3-F1-model_v4 SH3b domain-containing protein | 13 | 0.00E+00 | Acidobacteria bacterium |
| 547 | AF-A0A2G2ATW9-F1-model_v4 EF-hand domain-containing protein | 15.9 | 0.00E+00 | Sulfurimonas sp. |
| 548 | AF-A0A5C4ZL24-F1-model_v4 Enterotoxin | 10.1 | 0.00E+00 | Bacillus pacificus |
| 549 | AF-E6XBY3-F1-model_v4 CHAP domain containing protein | 21.1 | 0.00E+00 | Cellulophaga algicola DSM 14237 |
| 550 | AF-A0A7X1NSV4-F1-model_v4 Excalibur domain-containing protein | 7.8 | 0.00E+00 | Deinococcus terrestris |
| 551 | AF-A0A7C4L467-F1-model_v4 Uncharacterized protein | 11.3 | 0.00E+00 | Spirochaetes bacterium |
| 552 | AF-A0A1H5DJ53-F1-model_v4 Uncharacterized protein | 10.3 | 0.00E+00 | Amycolatopsis tolypomycina |
| 553 | AF-A0A537CQ40-F1-model_v4 SH3 domain-containing protein | 4.7 | 0.00E+00 | Betaproteobacteria bacterium |
| 554 | AF-A0A2W6W144-F1-model_v4 SH3 domain-containing protein | 7.4 | 0.00E+00 | Thauera sp. |
| 555 | AF-A0A523N2R2-F1-model_v4 SH3 domain-containing protein | 14.5 | 0.00E+00 | Nitrospina sp. |
| 556 | AF-A0A437LA83-F1-model_v4 SH3 domain-containing protein | 15.3 | 0.00E+00 | Rhodobacteraceae bacterium CCMM004 |
| 557 | AF-A0A015NN06-F1-model_v4 N-acetylmuramoyl-L-alanine amidase | 11 | 0.00E+00 | Paenibacillus darwinianus |
| 558 | AF-W6K6L5-F1-model_v4 Uncharacterized protein | 9.4 | 0.00E+00 | Magnetospira sp. QH-2 |
| 559 | AF-A0A813ES10-F1-model_v4 Hypothetical protein | 10.3 | 0.00E+00 | Polarella glacialis |
| 560 | AF-A0A359EIJ5-F1-model_v4 SH3b domain-containing protein | 11.6 | 0.00E+00 | Chloroflexi bacterium |

| | | | | |
|---|---|---|---|---|
| 561 | AF-A0A800M8R4-F1-model_v4 SH3b domain-containing protein | 15.2 | 0.00E+00 | Candidatus Lambdaproteobacteria bacterium |
| 562 | AF-A0A2H2WVA6-F1-model_v4 Enterotoxin | 11.5 | 0.00E+00 | Bacillus cereus |
| 563 | AF-A0A7V7XK07-F1-model_v4 Uncharacterized protein | 11.5 | 0.00E+00 | Anaerolineae bacterium |
| 564 | AF-A0A3D4UGQ6-F1-model_v4 Ligand-binding protein SH3 | 9.3 | 0.00E+00 | Moraxellaceae bacterium |
| 565 | AF-A0A7S1WSF0-F1-model_v4 Hypothetical protein | 10.6 | 0.00E+00 | Alexandrium catenella |
| 566 | AF-A0A264DU99-F1-model_v4 Hydrolase | 14.9 | 0.00E+00 | Paenibacillus odorifer |
| 567 | AF-A0A7C3JKJ2-F1-model_v4 SH3 domain-containing protein | 15.5 | 0.00E+00 | Deltaproteobacteria bacterium |
| 568 | AF-A0A2K1EEN9-F1-model_v4 Uncharacterized protein | 10.5 | 0.00E+00 | Paenibacillus sp. F4 |
| 569 | AF-C5CX27-F1-model_v4 SH3 type 3 domain protein | 12 | 0.00E+00 | Variovorax paradoxus S110 |
| 570 | AF-A0A5C1M8F8-F1-model_v4 Uncharacterized protein | 14.6 | 0.00E+00 | Geobacter sp. FeAm09 |
| 571 | AF-A0A6L4AA09-F1-model_v4 SUMF1/EgtB/PvdO family nonheme iron enzyme | 11.5 | 0.00E+00 | Anaerolineae bacterium |
| 572 | AF-A0A259KWG1-F1-model_v4 SH3b domain-containing protein | 13.2 | 0.00E+00 | Rhizobiales bacterium 39-66-18 |
| 573 | AF-A0A2N4XL50-F1-model_v4 SH3b domain-containing protein | 8.6 | 0.00E+00 | Uliginosibacterium sp. TH139 |
| 574 | AF-A0A2N7QPG9-F1-model_v4 SH3b domain-containing protein | 13.7 | 0.00E+00 | Dyella sp. AD56 |
| 575 | AF-A0A6M8HVL5-F1-model_v4 SH3 domain-containing protein | 14.9 | 0.00E+00 | Lichenicola cladoniae |
| 576 | AF-A0A4Y8QAB3-F1-model_v4 Uncharacterized protein | 14.1 | 0.00E+00 | Paenibacillus athensensis |
| 577 | AF-A0A7X6X4P0-F1-model_v4 C40 family peptidase | 9.8 | 0.00E+00 | Clostridiaceae bacterium |
| 578 | AF-A0A329KM62-F1-model_v4 Hydrolase Nlp/P60 | 12.1 | 0.00E+00 | Paenibacillus sp. YN15 |
| 579 | AF-A0A0G1N8I9-F1-model_v4 Uncharacterized protein | 12.8 | 0.00E+00 | Parcubacteria group bacterium GW2011_GWF2_44_8b |
| 580 | AF-B9IR39-F1-model_v4 Enterotoxin / cell-wall binding protein | 9.9 | 0.00E+00 | Bacillus cereus Q1 |
| 581 | AF-A0A3N5RP76-F1-model_v4 Uncharacterized protein | 13.7 | 0.00E+00 | Chloroflexi bacterium |
| 582 | AF-A0A840YBY4-F1-model_v4 Uncharacterized protein | 17.1 | 0.00E+00 | Sphingomonas xinjiangensis |
| 583 | AF-A0A1M3KQP1-F1-model_v4 SH3b domain-containing protein | 10.2 | 0.00E+00 | Devosia sp. 66-22 |
| 584 | AF-W8YYW4-F1-model_v4 SH3b domain-containing protein | 13.8 | 0.00E+00 | Paenibacillus sp. P22 |
| 585 | AF-A0A813DCY5-F1-model_v4 Hypothetical protein | 7.6 | 0.00E+00 | Polarella glacialis |
| 586 | AF-A0A382HWC6-F1-model_v4 SH3b domain-containing protein | 12 | 0.00E+00 | marine metagenome |
| 587 | AF-A0A6B1E156-F1-model_v4 SH3 domain-containing protein | 14.7 | 0.00E+00 | Chloroflexi bacterium |
| 588 | AF-A0A3N5LB54-F1-model_v4 Uncharacterized protein | 16.7 | 0.00E+00 | Chloroflexi bacterium |
| 589 | AF-A0A1A6KNT2-F1-model_v4 SH3b domain-containing protein | 12.2 | 0.00E+00 | Vibrio sp. UCD-FRSSP16_10 |
| 590 | AF-A0A2N5K0Y5-F1-model_v4 SH3b domain-containing protein | 13 | 0.00E+00 | Chloroflexi bacterium |
| 591 | AF-A0A3A9CBR4-F1-model_v4 SH3 domain-containing protein | 11.2 | 0.00E+00 | bacterium C-53 |
| 592 | AF-Q2RX13-F1-model_v4 Heat shock protein DnaJ-like | 11.6 | 0.00E+00 | Rhodospirillum rubrum ATCC 11170 |
| 593 | AF-A0A7S4QN38-F1-model_v4 Hypothetical protein | 11.1 | 0.00E+00 | Alexandrium monilatum |
| 594 | AF-A0A813LKM7-F1-model_v4 Hypothetical protein | 8.5 | 0.00E+00 | Polarella glacialis |
| 595 | AF-A0A7S1PJG8-F1-model_v4 Hypothetical protein | 6.1 | 0.00E+00 | Alexandrium catenella |

| | | | | |
|---|---|---|---|---|
| 596 | AF-H8H2C2-F1-model_v4 Putative lipoprotein | 10.8 | 0.00E+00 | Deinococcus gobiensis I-0 |
| 597 | AF-A0A4Q6BMW2-F1-model_v4 SH3 domain-containing protein | 17.3 | 0.00E+00 | Proteobacteria bacterium |
| 598 | AF-A0A417DKM3-F1-model_v4 SH3 domain-containing protein | 10.3 | 0.00E+00 | Ruminococcus sp. AM36-17 |
| 599 | AF-A0A7S1RG63-F1-model_v4 Hypothetical protein | 12 | 0.00E+00 | Alexandrium catenella |
| 600 | AF-A0A383BPH8-F1-model_v4 Uncharacterized protein | 15.9 | 0.00E+00 | marine metagenome |
| 601 | AF-A0A2H0SG33-F1-model_v4 Uncharacterized protein | 10.8 | 0.00E+00 | Parcubacteria group bacterium CG10_big_fil_rev_8_21_14_0_10_38_31 |
| 602 | AF-A0A4Q0J894-F1-model_v4 Uncharacterized protein | 10.1 | 0.00E+00 | Muribaculaceae bacterium Isolate-013 (NCI) |
| 603 | AF-A0A496W6X0-F1-model_v4 SH3b domain-containing protein | 13.6 | 0.00E+00 | Gammaproteobacteria bacterium |
| 604 | AF-A0A7X8VZI5-F1-model_v4 Dockerin domain-containing protein | 12.3 | 0.00E+00 | Clostridiaceae bacterium |
| 605 | AF-A0A1C4WKT2-F1-model_v4 Uncharacterized protein | 12.3 | 0.00E+00 | Micromonospora echinospora |
| 606 | AF-H3SM09-F1-model_v4 Uncharacterized protein | 15.6 | 0.00E+00 | Paenibacillus dendritiformis C454 |
| 607 | AF-A0A853VIW2-F1-model_v4 Ligand-binding protein SH3 | 14.6 | 0.00E+00 | Mesorhizobium sp. LCM 4577 |
| 608 | AF-S9Q9M8-F1-model_v4 SH3b domain-containing protein | 11.3 | 0.00E+00 | Salipiger mucosus DSM 16094 |
| 609 | AF-A0A7W8EMB5-F1-model_v4 Uncharacterized protein YraI | 12.8 | 0.00E+00 | Pseudochrobactrum saccharolyticum |
| 610 | AF-A0A2Y9ADP2-F1-model_v4 SH3 domain-containing protein | 13.4 | 0.00E+00 | Jannaschia seohaensis |
| 611 | AF-A0A0F7JYD6-F1-model_v4 Uncharacterized protein | 11.1 | 0.00E+00 | Sedimenticola thiotaurini |
| 612 | AF-A0A2B8HVK1-F1-model_v4 Enterotoxin | 9.4 | 0.00E+00 | Bacillus anthracis |
| 613 | AF-A0A559KDK8-F1-model_v4 NlpC/P60 family protein | 8.7 | 0.00E+00 | Paenibacillus sp. JC52 |
| 614 | AF-A0A2N1T019-F1-model_v4 SH3b domain-containing protein | 12.7 | 0.00E+00 | Spirochaetae bacterium HGW-Spirochaetae-10 |
| 615 | AF-A0A136KSS8-F1-model_v4 Uncharacterized protein | 15.5 | 0.00E+00 | Chloroflexi bacterium OLB15 |
| 616 | AF-A0A329LEG1-F1-model_v4 MurNAc-LAA domain-containing protein | 9.2 | 0.00E+00 | Paenibacillus sp. YN15 |
| 617 | AF-A0A3M1IK13-F1-model_v4 SH3b domain-containing protein | 13.7 | 0.00E+00 | Verrucomicrobia bacterium |
| 618 | AF-A0A7V9N1P8-F1-model_v4 N-acetylmuramoyl-L-alanine amidase | 9.6 | 0.00E+00 | Chloroflexia bacterium |
| 619 | AF-A0A7S1F706-F1-model_v4 Hypothetical protein | 8.7 | 0.00E+00 | Noctiluca scintillans |
| 620 | AF-A0A0L0DM35-F1-model_v4 Uncharacterized protein | 12.1 | 0.00E+00 | Thecamonas trahens ATCC 50062 |
| 621 | AF-A0A849I8T4-F1-model_v4 Uncharacterized protein | 6.5 | 0.00E+00 | Burkholderiales bacterium |
| 622 | AF-A0A4R1BEI4-F1-model_v4 SH3 domain-containing protein | 15.5 | 0.00E+00 | Parasulfuritortus cantonensis |
| 623 | AF-A0A7S4E577-F1-model_v4 Hypothetical protein | 13 | 0.00E+00 | Pelagomonas calceolata |
| 624 | AF-A0A2V9E388-F1-model_v4 SH3b domain-containing protein | 13.3 | 0.00E+00 | Acidobacteria bacterium |
| 625 | AF-A0A1H4H493-F1-model_v4 Uncharacterized conserved protein YraI | 13.3 | 0.00E+00 | Paraburkholderia sartisoli |
| 626 | AF-A0A3M0XLC1-F1-model_v4 Peptidase P60 | 10.2 | 0.00E+00 | Alphaproteobacteria bacterium |
| 627 | AF-A0A250JL17-F1-model_v4 Uncharacterized protein | 12.9 | 0.00E+00 | Cystobacter fuscus |
| 628 | AF-A0A7Y1V8Y6-F1-model_v4 C40 family peptidase | 8.6 | 0.00E+00 | Saprospiraceae bacterium |
| 629 | AF-A0A416JLC0-F1-model_v4 Uncharacterized protein | 9.1 | 0.00E+00 | Clostridium sp. AM49-4BH |
| 630 | AF-A0A1I0MS96-F1-model_v4 Metal-dependent hydrolase, beta-lactamase superfamily II | 9.4 | 0.00E+00 | Roseivirga pacifica |

| | | | | |
|---|---|---|---|---|
| 631 | AF-A0A6L3EU07-F1-model_v4 DUF3380 domain-containing protein | 7.8 | 0.00E+00 | Chloroflexi bacterium |
| 632 | AF-A0A5C0SAR0-F1-model_v4 SH3 domain-containing protein | 12.7 | 0.00E+00 | Crassaminicella thermophila |
| 633 | AF-A0A554KUL1-F1-model_v4 Uncharacterized protein | 14.4 | 0.00E+00 | Parcubacteria group bacterium Gr01-1014 13 |
| 634 | AF-A0A7S0ZSN2-F1-model_v4 Hypothetical protein | 8.2 | 0.00E+00 | Noctiluca scintillans |
| 635 | AF-A0A484HKN5-F1-model_v4 Uncharacterized protein | 15 | 0.00E+00 | uncultured Desulfobacteraceae bacterium |
| 636 | AF-A0A3C1B404-F1-model_v4 Uncharacterized protein | 9 | 0.00E+00 | Clostridiales bacterium |
| 637 | AF-A0A533QP93-F1-model_v4 SH3b domain-containing protein | 10.8 | 0.00E+00 | Candidatus Accumulibacter sp. |
| 638 | AF-A0A562FA05-F1-model_v4 Uncharacterized protein YraI | 9.7 | 0.00E+00 | Aminobacter sp. J15 |
| 639 | AF-A0A3M2AGB2-F1-model_v4 SH3 domain-containing protein | 13.7 | 0.00E+00 | Chloroflexi bacterium |
| 640 | AF-A0A845C7P4-F1-model_v4 SH3 domain-containing protein | 9.6 | 0.00E+00 | Caldilineaceae bacterium SB0666 bin 21 |
| 641 | AF-A0A350IEI8-F1-model_v4 Uncharacterized protein | 9.5 | 0.00E+00 | Saprospirales bacterium |
| 642 | AF-A0A7C1MQN2-F1-model_v4 SH3 domain-containing protein | 11.1 | 0.00E+00 | Candidatus Moranbacteria bacterium |
| 643 | AF-A0A1Q6KX27-F1-model_v4 Uncharacterized protein | 10.8 | 0.00E+00 | Clostridiales bacterium 41 12 two minus |
| 644 | AF-A9IRC1-F1-model_v4 SH3b domain-containing protein | 8 | 0.00E+00 | Bartonella tribocorum CIP 105476 |
| 645 | AF-A0A6H2KV82-F1-model_v4 SH3 domain-containing protein | 11.3 | 0.00E+00 | Paracoccus sanguinis |
| 646 | AF-A0A536T6M2-F1-model_v4 Uncharacterized protein | 6.9 | 0.00E+00 | Betaproteobacteria bacterium |
| 647 | AF-A0A5B2ZDE0-F1-model_v4 SH3 domain-containing protein | 12.7 | 0.00E+00 | Arenimonas fontis |
| 648 | AF-A0A7S1RQH0-F1-model_v4 Hypothetical protein | 10.9 | 0.00E+00 | Alexandrium catenella |
| 649 | AF-A0A3Q9IC01-F1-model_v4 Uncharacterized protein | 14.9 | 0.00E+00 | Paenibacillus lutimineralis |
| 650 | AF-A5V1I3-F1-model_v4 SH3, type 3 domain protein | 12.2 | 0.00E+00 | Roseiflexus sp. RS-1 |
| 651 | AF-A0A4U2LDH3-F1-model_v4 Ribonuclease | 8.6 | 0.00E+00 | Puteibacter caeruleilacunae |
| 652 | AF-A0A1I0DHU5-F1-model_v4 Uncharacterized protein | 17.3 | 0.00E+00 | Stigmatella erecta |
| 653 | AF-A0A0D5NMP1-F1-model_v4 Uncharacterized protein | 11.4 | 0.00E+00 | Paenibacillus beijingensis |
| 654 | AF-A0A2T7FST3-F1-model_v4 SH3b domain-containing protein | 11.2 | 0.00E+00 | Thalassorhabdomicrobium marinisediminis |
| 655 | AF-R9C7Y3-F1-model_v4 Uncharacterized protein | 10.4 | 0.00E+00 | Clostridium sartagoforme AAU1 |
| 656 | AF-A0A3N5DXK2-F1-model_v4 DUF3761 domain-containing protein | 13 | 0.00E+00 | Bacteroidales bacterium |
| 657 | AF-A0A4Y9QB02-F1-model_v4 SH3 domain-containing protein | 11.6 | 0.00E+00 | Oxalobacteraceae bacterium OM1 |
| 658 | AF-A0A2R6K994-F1-model_v4 Peptidase_M23 domain-containing protein | 12.1 | 0.00E+00 | Halobacteriales archaeon QS_9_68_17 |
| 659 | AF-A0A679IXS5-F1-model_v4 SH3b domain-containing protein | 9.7 | 0.00E+00 | Variovorax paradoxus |
| 660 | AF-A0A661I2V2-F1-model_v4 Uncharacterized protein | 14.3 | 0.00E+00 | Epsilonproteobacteria bacterium |
| 661 | AF-E7QML6-F1-model_v4 N-acetylmuramyl-L-alanine amidase, negative regulator of AmpC, AmpD | 6.5 | 0.00E+00 | Haladaptatus paucihalophilus DX253 |
| 662 | AF-A0A4U0ZGZ3-F1-model_v4 SH3 domain-containing protein | 10 | 0.00E+00 | Desulforhopalus sp. IMCC35007 |
| 663 | AF-A0A7C4IM60-F1-model_v4 SH3b domain-containing protein | 9.3 | 0.00E+00 | Nitrospirae bacterium |
| 664 | AF-A0A4R8IJW7-F1-model_v4 SH3 domain-containing protein | 13.5 | 0.00E+00 | Epilithonimonas xixisoli |
| 665 | AF-A0A6H9Y7M6-F1-model_v4 SH3 domain-containing protein | 8.6 | 0.00E+00 | Betaproteobacteria bacterium SCN1 |

| | | | | |
|---|---|---|---|---|
| 666 | AF-A0A4D4JIH2-F1-model_v4 Uncharacterized protein | 12.3 | 0.00E+00 | Acidovorax sp. NB1 |
| 667 | AF-A0A6C2CIY5-F1-model_v4 SH3b domain-containing protein | 11.4 | 0.00E+00 | Zoogloea oleivorans |
| 668 | AF-A0A813JCP9-F1-model_v4 Hypothetical protein | 9.3 | 0.00E+00 | Polarella glacialis |
| 669 | AF-A0A6J4T6U8-F1-model_v4 SH3_16 domain-containing protein | 9.6 | 0.00E+00 | uncultured Sphingomonas sp. |
| 670 | AF-A0A411Z5H3-F1-model_v4 SH3 domain-containing protein | 17.5 | 0.00E+00 | Tabrizicola alkalilacus |
| 671 | AF-A0A6N7ZZR4-F1-model_v4 SH3b domain-containing protein | 12.3 | 0.00E+00 | Roseibium sp. RKSG952 |
| 672 | AF-A0A6N0ZAC9-F1-model_v4 SH3 domain-containing protein | 10.8 | 0.00E+00 | Candidatus Accumulibacter sp. |
| 673 | AF-A0A3N5K963-F1-model_v4 SH3 domain-containing protein | 6.6 | 0.00E+00 | Zetaproteobacteria bacterium |
| 674 | AF-A0A3N5MRS9-F1-model_v4 SH3b domain-containing protein | 13.3 | 0.00E+00 | Chloroflexi bacterium |
| 675 | AF-A0A1S2DUG1-F1-model_v4 SH3b domain-containing protein | 14.6 | 0.00E+00 | Agrobacterium vitis |
| 676 | AF-H7EM44-F1-model_v4 SH3 type 3 domain protein | 9.6 | 0.00E+00 | Treponema saccharophilum DSM 2985 |
| 677 | AF-A0A6P2E1T2-F1-model_v4 SH3b domain-containing protein | 7.6 | 0.00E+00 | Variovorax sp. PBL-E5 |
| 678 | AF-A0A523K7X8-F1-model_v4 Uncharacterized protein | 11.1 | 0.00E+00 | Gammaproteobacteria bacterium |
| 679 | AF-A0A7V9VI36-F1-model_v4 SH3 domain-containing protein | 11.2 | 0.00E+00 | Chloroflexia bacterium |
| 680 | AF-A0A4U2MIB5-F1-model_v4 Enterotoxin | 11.8 | 0.00E+00 | Bacillus wiedmannii |
| 681 | AF-A0A7V9VUY8-F1-model_v4 SH3 domain-containing protein | 9.4 | 0.00E+00 | Chloroflexia bacterium |
| 682 | AF-A0A6J4TXK0-F1-model_v4 Non-specific serine/threonine protein kinase | 9.8 | 0.00E+00 | uncultured Thermomicrobiales bacterium |
| 683 | AF-A0A813FCN7-F1-model_v4 Hypothetical protein | 9.7 | 0.00E+00 | Polarella glacialis |
| 684 | AF-A0A4Q2XC19-F1-model_v4 Uncharacterized protein | 9.2 | 0.00E+00 | Verrucomicrobiaceae bacterium |
| 685 | AF-A0A515EJF3-F1-model_v4 DUF4384 domain-containing protein | 10.3 | 0.00E+00 | Rhodoferax sediminis |
| 686 | AF-A0A497B2L4-F1-model_v4 SH3b domain-containing protein | 14.4 | 0.00E+00 | Chloroflexi bacterium |
| 687 | AF-A0A353MWG8-F1-model_v4 SH3b domain-containing protein | 13 | 0.00E+00 | Clostridium sp. |
| 688 | AF-A0A839NCJ2-F1-model_v4 Long-subunit fatty acid transport protein | 11.4 | 0.00E+00 | Flexivirga oryzae |
| 689 | AF-A0A3A8T6L7-F1-model_v4 Uncharacterized protein | 13.5 | 0.00E+00 | Corallococcus sp. AB038B |
| 690 | AF-A0A2E9F472-F1-model_v4 Uncharacterized protein | 8.4 | 0.00E+00 | Candidatus Poribacteria bacterium |
| 691 | AF-A0A7C4J9E1-F1-model_v4 SH3 domain-containing protein | 14.7 | 0.00E+00 | Chloroflexi bacterium |
| 692 | AF-A0A179FDG5-F1-model_v4 Uncharacterized protein | 10 | 0.00E+00 | Pochonia chlamydosporia 170 |
| 693 | AF-A0A1N7AHH7-F1-model_v4 SH3 domain-containing protein | 12 | 0.00E+00 | Rhizobium sp. RU35A |
| 694 | AF-A0A536UQZ2-F1-model_v4 Uncharacterized protein | 6.9 | 0.00E+00 | Betaproteobacteria bacterium |
| 695 | AF-A0A5K8A8F5-F1-model_v4 Uncharacterized protein | 12.9 | 0.00E+00 | Desulfosarcina ovata subsp. ovata |
| 696 | AF-A0A327JXT4-F1-model_v4 Uncharacterized protein | 7.8 | 0.00E+00 | Rhodobium orientis |
| 697 | AF-A0A2V7NNI0-F1-model_v4 SH3b domain-containing protein | 10.6 | 0.00E+00 | Gemmatimonadetes bacterium |
| 698 | AF-A0A2C1REA8-F1-model_v4 Enterotoxin | 11 | 0.00E+00 | Bacillus cereus |
| 699 | AF-A0A7V9HLL3-F1-model_v4 SH3 domain-containing protein | 12.3 | 0.00E+00 | Chloroflexia bacterium |
| 700 | AF-A0A813H100-F1-model_v4 Hypothetical protein | 9.3 | 0.00E+00 | Polarella glacialis |
| 701 | AF-A0A436F988-F1-model_v4 SH3 domain-containing protein | 15.5 | 0.00E+00 | Mesorhizobium sp. M7A.F.Ca.CA.004.05.1.1 |

| | | | | |
|---|---|---|---|---|
| 702 | AF-A0A6I0EC38-F1-model_v4 SH3 domain-containing protein | 10.3 | 0.00E+00 | Hyphomicrobium sp. |
| 703 | AF-A0A535XQQ3-F1-model_v4 SH3_16 domain-containing protein | 15.4 | 0.00E+00 | Chloroflexi bacterium |
| 704 | AF-A0A2E3WU05-F1-model_v4 Uncharacterized protein | 11.1 | 0.00E+00 | Bdellovibrionaceae bacterium |
| 705 | AF-A0A5B0WCU2-F1-model_v4 SH3 domain-containing protein | 10.8 | 0.00E+00 | Rhizobium tropici |
| 706 | AF-A0A2S5R1P7-F1-model_v4 Ligand-binding protein SH3 | 9 | 0.00E+00 | Methylotenera sp. |
| 707 | AF-A0A522M241-F1-model_v4 SH3 domain-containing protein | 5.1 | 0.00E+00 | Gammaproteobacteria bacterium |
| 708 | AF-A0A7V7PR49-F1-model_v4 Uncharacterized protein | 13.4 | 0.00E+00 | Aureimonas leprariae |
| 709 | AF-A0A1I4E894-F1-model_v4 SH3 domain-containing protein | 10 | 0.00E+00 | Falsiroseomonas stagni DSM 19981 |
| 710 | AF-A0A809KCC0-F1-model_v4 Uncharacterized protein | 9.5 | 0.00E+00 | Lactobacillus acidophilus |
| 711 | AF-A0A7S3HTJ6-F1-model_v4 Hypothetical protein | 9.2 | 0.00E+00 | Spumella elongata |
| 712 | AF-A0A519LPC8-F1-model_v4 SH3 domain-containing protein | 9.8 | 0.00E+00 | Acidovorax sp. |
| 713 | AF-A0A3P1WDR9-F1-model_v4 SH3 domain-containing protein | 10.4 | 0.00E+00 | Tessaracoccus sp. OH4464_COT-324 |
| 714 | AF-A0A2V9QBU7-F1-model_v4 Uncharacterized protein | 13.7 | 0.00E+00 | Acidobacteria bacterium |
| 715 | AF-A0A2N5ZIF5-F1-model_v4 Uncharacterized protein | 12.1 | 0.00E+00 | Candidatus Muirbacterium halophilum |
| 716 | AF-A0A2M8CQS6-F1-model_v4 Uncharacterized protein | 12.4 | 0.00E+00 | Anaerolineae bacterium CG 4 9 14 3 um filter 57 17 |
| 717 | AF-A0A0M0K0A2-F1-model_v4 Uncharacterized protein | 9 | 0.00E+00 | Chrysochromulina tobinii |
| 718 | AF-A0A7S3HM67-F1-model_v4 Hypothetical protein | 12.5 | 0.00E+00 | Spumella elongata |
| 719 | AF-A0A3P3XTA2-F1-model_v4 Uncharacterized protein | 10 | 0.00E+00 | uncultured spirochete |
| 720 | AF-A0A660CI61-F1-model_v4 Uncharacterized protein | 13.6 | 0.00E+00 | Prauserella rugosa |
| 721 | AF-A0A2T4IEC8-F1-model_v4 Uncharacterized protein | 9 | 0.00E+00 | Pseudothauera lacus |
| 722 | AF-A0A099T731-F1-model_v4 SH3b domain-containing protein | 14.5 | 0.00E+00 | Thalassobacter sp. 16PALIMAR09 |
| 723 | AF-A0A522D5P1-F1-model_v4 Uncharacterized protein | 9.4 | 0.00E+00 | Alphaproteobacteria bacterium |
| 724 | AF-A0A0E4GBX8-F1-model_v4 SH3-like domain, bacterial-type | 8 | 0.00E+00 | Syntrophomonas zehnderi OL-4 |
| 725 | AF-A0A3L7J8T6-F1-model_v4 SH3b domain-containing protein | 8.8 | 0.00E+00 | Notoacmeibacter ruber |
| 726 | AF-A0A1F4ARW6-F1-model_v4 Uncharacterized protein | 13.1 | 0.00E+00 | Betaproteobacteria bacterium RIFCSPLOWO2_02_FULL_63_19 |
| 727 | AF-A0A1T4LS12-F1-model_v4 N-acetylmuramoyl-L-alanine amidase | 12.1 | 0.00E+00 | Anaerorhabdus furcosa |
| 728 | AF-A0A1J1C560-F1-model_v4 SH3 domain-containing protein | 11.5 | 0.00E+00 | Caldithrix abyssi DSM 13497 |
| 729 | AF-A0A2U0XDH5-F1-model_v4 Uncharacterized protein | 12.7 | 0.00E+00 | Streptomyces sp. 3212.2 |
| 730 | AF-A0A7S3HRP9-F1-model_v4 Hypothetical protein | 11.6 | 0.00E+00 | Spumella elongata |
| 731 | AF-A0A7S2NAA2-F1-model_v4 Hypothetical protein | 11.7 | 0.00E+00 | Alexandrium andersonii |
| 732 | AF-A0A4Q4ICN4-F1-model_v4 Mannosyl-glycoprotein endo-beta-N-acetylglucosamidase | 15.8 | 0.00E+00 | Sporolactobacillus sp. THM19-2 |
| 733 | AF-A0A813LRB3-F1-model_v4 Hypothetical protein | 12.8 | 0.00E+00 | Polarella glacialis |
| 734 | AF-A0A356M3F6-F1-model_v4 SH3b domain-containing protein | 11.2 | 0.00E+00 | Clostridiales bacterium |
| 735 | AF-A0A7W0TXX3-F1-model_v4 SH3 domain-containing protein | 11.9 | 0.00E+00 | Chloroflexia bacterium |
| 736 | AF-A0A7S1KWS7-F1-model_v4 Hypothetical protein | 14.1 | 0.00E+00 | Alexandrium catenella |

| | | | | |
|---|---|---|---|---|
| 737 | AF-A0A7U0A0J4-F1-model_v4 Uncharacterized protein | 12 | 0.00E+00 | Enterococcus casseliflavus |
| 738 | AF-A0A1N7PGR5-F1-model_v4 Uncharacterized protein | 12.7 | 0.00E+00 | Achromobacter sp. MFA1 R4 |
| 739 | AF-G5J970-F1-model_v4 Uncharacterized protein | 12.7 | 0.00E+00 | Crocosphaera watsonii WH 0003 |
| 740 | AF-A0A347UKQ7-F1-model_v4 SH3 domain-containing protein | 12.1 | 0.00E+00 | Profundibacter amoris |
| 741 | AF-A0A431I3Z3-F1-model_v4 SH3 domain-containing protein | 7.6 | 0.00E+00 | Neisseriaceae bacterium |
| 742 | AF-A0A7G1HRC2-F1-model_v4 SH3b domain-containing protein | 14.3 | 0.00E+00 | Helicobacter pylori |
| 743 | AF-A0A7S4REF8-F1-model_v4 Hypothetical protein | 10.9 | 0.00E+00 | Alexandrium monilatum |
| 744 | AF-A0A0G1K0L9-F1-model_v4 Endonuclease/exonuclease/phosphatase | 15.8 | 0.00E+00 | Candidatus Giovannonibacteria bacterium GW2011 GWA2 44 13b |
| 745 | AF-A0A7W5DCS2-F1-model_v4 Uncharacterized protein YraI | 6.8 | 0.00E+00 | Variovorax sp. Sphag1AA |
| 746 | AF-A0A7S3HG59-F1-model_v4 Hypothetical protein | 9.5 | 0.00E+00 | Spumella elongata |
| 747 | AF-A0A534VHE8-F1-model_v4 SH3 domain-containing protein | 8.5 | 0.00E+00 | Deltaproteobacteria bacterium |
| 748 | AF-A0A1F5VNB9-F1-model_v4 Uncharacterized protein | 7.7 | 0.00E+00 | Candidatus Giovannonibacteria bacterium RIFCSPHIGHO2 02 42 15 |
| 749 | AF-A0A3M1UU40-F1-model_v4 Uncharacterized protein | 11.2 | 0.00E+00 | Gammaproteobacteria bacterium |
| 750 | AF-A0A259TWU3-F1-model_v4 Uncharacterized protein | 14.8 | 0.00E+00 | Rubricoccus marinus |
| 751 | AF-A0A7S2F3Q6-F1-model_v4 Hypothetical protein | 13 | 0.00E+00 | Alexandrium andersonii |
| 752 | AF-A0A562CDS4-F1-model_v4 Uncharacterized protein YraI | 11.7 | 0.00E+00 | Mesorhizobium sp. J18 |
| 753 | AF-A0A838S012-F1-model_v4 Uncharacterized protein | 8.9 | 0.00E+00 | Pyrinomonadaceae bacterium |
| 754 | AF-A0A7C3FY88-F1-model_v4 Uncharacterized protein | 13.7 | 0.00E+00 | Thermoflexia bacterium |
| 755 | AF-A0A2N5DQH5-F1-model_v4 Hydrolase Nlp/P60 | 11.8 | 0.00E+00 | Caulobacter zeae |
| 756 | AF-A0A2M8ZGP3-F1-model_v4 Uncharacterized protein YraI | 8.8 | 0.00E+00 | Afipia broomeae |
| 757 | AF-A0A512CLK7-F1-model_v4 NlpC/P60 domain-containing protein | 9.1 | 0.00E+00 | Alicyclobacillus acidoterrestris |
| 758 | AF-A0A4Q3XRY8-F1-model_v4 SH3 domain-containing protein | 14.8 | 0.00E+00 | Alphaproteobacteria bacterium |
| 759 | AF-A0A3M1FKY3-F1-model_v4 SH3 domain-containing protein | 10.1 | 0.00E+00 | Candidatus Dadabacteria bacterium |
| 760 | AF-A0A7X1HFI4-F1-model_v4 SH3 domain-containing protein | 12.5 | 0.00E+00 | Rhizobium sp. AQ_MP |
| 761 | AF-D1C1X4-F1-model_v4 SH3b domain-containing protein | 10.7 | 0.00E+00 | Sphaerobacter thermophilus DSM 20745 |
| 762 | AF-A0A239I588-F1-model_v4 SH3 domain-containing protein | 10.2 | 0.00E+00 | Tropicimonas sediminicola |
| 763 | AF-A0A7Y5H8Q8-F1-model_v4 Uncharacterized protein | 12.5 | 0.00E+00 | Candidatus Brocadiae bacterium |
| 764 | AF-A0A7S3T7M0-F1-model_v4 Hypothetical protein | 15 | 0.00E+00 | Strombidinopsis acuminata |
| 765 | AF-A0A2A7EEF2-F1-model_v4 SH3b domain-containing protein | 6.9 | 0.00E+00 | Bacillus cereus |
| 766 | AF-A0A2S8WPV8-F1-model_v4 D-alanyl-D-alanine carboxypeptidase | 13.6 | 0.00E+00 | Arthrobacter sp. MYb227 |
| 767 | AF-A0A812JZG6-F1-model_v4 Hypothetical protein | 10.2 | 0.00E+00 | Symbiodinium pilosum |
| 768 | AF-A0A2A2ASD0-F1-model_v4 SH3b domain-containing protein | 14.2 | 0.00E+00 | Vandammella animalimorsus |
| 769 | AF-A0A6L5X6J4-F1-model_v4 SH3 domain-containing protein | 9.2 | 0.00E+00 | Porcincola intestinalis |
| 770 | AF-A0A2M7GDB3-F1-model_v4 SH3b domain-containing protein | 13 | 0.00E+00 | Anaerolineae bacterium CG17_big_fil_post_rev_8_21_14_2 50 57 27 |

| | | | | |
|---|---|---|---|---|
| 771 | AF-A0A233VJD7-F1-model_v4 Enterotoxin | 12.9 | 0.00E+00 | Finegoldia magna |
| 772 | AF-T0SHD7-F1-model_v4 SH3 domain protein | 7.8 | 0.00E+00 | Bacteriovorax sp. BSW11_IV |
| 773 | AF-A0A1Y5RJ51-F1-model_v4 Bacterial SH3 domain protein | 13 | 0.00E+00 | Aquimixticola soesokkakensis |
| 774 | AF-A0A1F3X8Y0-F1-model_v4 Uncharacterized protein | 11.3 | 0.00E+00 | Bdellovibrionales bacterium RIFOXYD1_FULL_44_7 |
| 775 | AF-A0A3N5G613-F1-model_v4 Uncharacterized protein | 18 | 0.00E+00 | Chloroflexi bacterium |
| 776 | AF-A0A661WIZ4-F1-model_v4 SH3b domain-containing protein | 11.9 | 0.00E+00 | Chloroflexi bacterium |
| 777 | AF-A0A3D5SQC8-F1-model_v4 Uncharacterized protein | 15.6 | 0.00E+00 | Blastocatellia bacterium |
| 778 | AF-A0A6C0FZ37-F1-model_v4 SH3 domain-containing protein | 12.8 | 0.00E+00 | Paenibacillus lycopersici |
| 779 | AF-A0A5C4JF14-F1-model_v4 Uncharacterized protein | 9.2 | 0.00E+00 | Actinomadura soli |
| 780 | AF-A0A2M8NIK7-F1-model_v4 Uncharacterized protein | 9 | 0.00E+00 | Phototrophicales bacterium |
| 781 | AF-A0A7S2M7E2-F1-model_v4 Hypothetical protein | 10.2 | 0.00E+00 | Brandtodinium nutricula |
| 782 | AF-A0A1M3ACF2-F1-model_v4 SH3b domain-containing protein | 10.6 | 0.00E+00 | Alphaproteobacteria bacterium 62-8 |
| 783 | AF-A0A2W5A1U0-F1-model_v4 Uncharacterized protein | 13.5 | 0.00E+00 | Pseudoxanthomonas suwonensis |
| 784 | AF-A0A0M6Z2B8-F1-model_v4 SH3b domain-containing protein | 11.2 | 0.00E+00 | Roseibium album |
| 785 | AF-A0A1G6VJL4-F1-model_v4 Uncharacterized protein | 9.3 | 0.00E+00 | Paenibacillus sp. CF095 |
| 786 | AF-A0A7W1DVI8-F1-model_v4 Uncharacterized protein | 14.1 | 0.00E+00 | Chloroflexia bacterium |
| 787 | AF-B8HV27-F1-model_v4 SH3b domain-containing protein | 8.2 | 0.00E+00 | Cyanothece sp. PCC 7425 |
| 788 | AF-A0A7S1L1S9-F1-model_v4 Hypothetical protein | 10.6 | 0.00E+00 | Alexandrium catenella |
| 789 | AF-A0A7X6Z7L1-F1-model_v4 SH3 domain-containing protein | 13.4 | 0.00E+00 | Clostridiaceae bacterium |
| 790 | AF-L7U4G5-F1-model_v4 TPR repeat containing protein | 7.6 | 0.00E+00 | Myxococcus stipitatus DSM 14675 |
| 791 | AF-A0A504KWS3-F1-model_v4 SH3 domain-containing protein | 11.1 | 0.00E+00 | Mesorhizobium sp. B1-1-5 |
| 792 | AF-A0A7V7BCT5-F1-model_v4 Uncharacterized protein | 11.7 | 0.00E+00 | Alphaproteobacteria bacterium |
| 793 | AF-A0A561T7D8-F1-model_v4 Uncharacterized protein | 12.7 | 0.00E+00 | Kitasatospora viridis |
| 794 | AF-A0A3A4WDR0-F1-model_v4 SH3 domain-containing protein | 10.4 | 0.00E+00 | Actinomycetia bacterium |
| 795 | AF-A0A258BWJ5-F1-model_v4 SH3b domain-containing protein | 15.4 | 0.00E+00 | Sphingobium sp. 32-64-5 |
| 796 | AF-D2VG09-F1-model_v4 Predicted protein | 10.1 | 0.00E+00 | Naegleria gruberi |
| 797 | AF-A0A417DG83-F1-model_v4 SH3 domain-containing protein | 12.4 | 0.00E+00 | Ruminococcus sp. AM40-10AC |
| 798 | AF-A0A1Z4IHD7-F1-model_v4 SH3b domain-containing protein | 10.3 | 0.00E+00 | Nostoc sp. NIES-2111 |
| 799 | AF-A0A4Q7E1M3-F1-model_v4 Uncharacterized protein | 15.5 | 0.00E+00 | Leptolyngbya sp. LK |
| 800 | AF-A0A812QWG6-F1-model_v4 Hypothetical protein | 10.8 | 0.00E+00 | Symbiodinium sp. KB8 |
| 801 | AF-A0A2M6UZ49-F1-model_v4 DUF4384 domain-containing protein | 11.1 | 0.00E+00 | Limnohabitans sp. B9-3 |
| 802 | AF-A0A1X7C0N7-F1-model_v4 SH3b domain-containing protein | 11.9 | 0.00E+00 | Desulfovibrio gilichinskyi |
| 803 | AF-A0A538FKV1-F1-model_v4 SH3 domain-containing protein | 13.6 | 0.00E+00 | Actinomycetia bacterium |
| 804 | AF-A0A1F6U6C4-F1-model_v4 SH3b domain-containing protein | 9.9 | 0.00E+00 | Candidatus Muproteobacteria bacterium RIFCSPLOWO2_01_FULL_60_18 |
| 805 | AF-A0A0L1JN42-F1-model_v4 SH3b domain-containing protein | 15.2 | 0.00E+00 | Pseudaestuariivita atlantica |

| | | | | |
|---|---|---|---|---|
| 806 | AF-J0R1C4-F1-model_v4 SH3b domain-containing protein | 12.7 | 0.00E+00 | Bartonella tamiae Th239 |
| 807 | AF-E1X0U8-F1-model_v4 Putative membrane protein | 12.2 | 0.00E+00 | Halobacteriovorax marinus SJ |
| 808 | AF-A0A1H0N7G7-F1-model_v4 SH3b domain-containing protein | 11.7 | 0.00E+00 | Nakamurella panacisegetis |
| 809 | AF-A0A3M1CGV7-F1-model_v4 NlpC/P60 domain-containing protein | 8.9 | 0.00E+00 | candidate division Zixibacteria bacterium |
| 810 | AF-A0A7S1MD63-F1-model_v4 Hypothetical protein | 13 | 0.00E+00 | Alexandrium catenella |
| 811 | AF-A0A372GBD0-F1-model_v4 HTH cro/C1-type domain-containing protein | 14.3 | 0.00E+00 | Actinomadura spongiicola |
| 812 | AF-A0A538PNL0-F1-model_v4 Uncharacterized protein | 14.4 | 0.00E+00 | Deltaproteobacteria bacterium |
| 813 | AF-A0A7S3TMB9-F1-model_v4 Hypothetical protein | 8.8 | 0.00E+00 | Strombidinopsis acuminata |
| 814 | AF-A0A1R0WKX8-F1-model_v4 Hydrolase | 10.7 | 0.00E+00 | Paenibacillus odorifer |
| 815 | AF-A0A547PMM9-F1-model_v4 SH3b domain-containing protein | 13.7 | 0.00E+00 | Paenimaribius caenipelagi |
| 816 | AF-A0A1H3ND04-F1-model_v4 SH3 domain-containing protein | 14.4 | 0.00E+00 | Hymenobacter psychrophilus |
| 817 | AF-A0A438W8P0-F1-model_v4 SH3b domain-containing protein | 14.7 | 0.00E+00 | Helicobacter pylori |
| 818 | AF-A0A2E2EEL8-F1-model_v4 SH3b domain-containing protein | 10.6 | 0.00E+00 | Halobacteriovoraceae bacterium |
| 819 | AF-A0A813M439-F1-model_v4 Hypothetical protein | 11.2 | 0.00E+00 | Polarella glacialis |
| 820 | AF-A0A7V2JG34-F1-model_v4 NlpC/P60 domain-containing protein | 11.1 | 0.00E+00 | Thermotogaceae bacterium |
| 821 | AF-A0A2E5SIH8-F1-model_v4 Uncharacterized protein | 7.6 | 0.00E+00 | Candidatus Marinimicrobia bacterium |
| 822 | AF-A0A495F6J1-F1-model_v4 Curli biogenesis system outer membrane secretion channel CsgG | 8.3 | 0.00E+00 | Sphingomonas elodea |
| 823 | AF-A0A7Z9YZM7-F1-model_v4 SH3b domain-containing protein | 16.6 | 0.00E+00 | Desulfobacterales bacterium |
| 824 | AF-A0A7V5R994-F1-model_v4 DUF2075 domain-containing protein | 12.7 | 0.00E+00 | Deltaproteobacteria bacterium |
| 825 | AF-A0A2S6U0E4-F1-model_v4 Uncharacterized protein | 12.8 | 0.00E+00 | Alphaproteobacteria bacterium MarineAlpha3 Bin7 |
| 826 | AF-R5Z250-F1-model_v4 Glycoside hydrolase family 18 | 9.4 | 0.00E+00 | Clostridium sp. CAG:492 |
| 827 | AF-A0A812PFQ6-F1-model_v4 ME1 protein | 16.5 | 0.00E+00 | Symbiodinium pilosum |
| 828 | AF-A0A847X3X7-F1-model_v4 SH3 domain-containing protein | 10.5 | 0.00E+00 | Gallicola sp. |
| 829 | AF-A0A455SF20-F1-model_v4 Uncharacterized protein | 9 | 0.00E+00 | Thermosporothrix sp. COM3 |
| 830 | AF-A0A7Y2FT42-F1-model_v4 SH3 domain-containing protein | 13.2 | 0.00E+00 | Hyphomicrobiales bacterium |
| 831 | AF-A0A318EB52-F1-model_v4 Uncharacterized protein | 10 | 0.00E+00 | Sinimarinibacterium flocculans |
| 832 | AF-A0A4U3BQN6-F1-model_v4 Enterotoxin | 9.7 | 0.00E+00 | Bacillus cereus |
| 833 | AF-A0A0G1RW11-F1-model_v4 SH3b domain-containing protein | 8 | 0.00E+00 | Candidatus Beckwithbacteria bacterium GW2011_GWB1_47_15 |
| 834 | AF-A0A2R8BHG2-F1-model_v4 SH3b domain-containing protein | 15.1 | 0.00E+00 | Ascidiaceihabitans donghaensis |
| 835 | AF-A0A6N8F3U9-F1-model_v4 SH3 domain-containing protein | 12.6 | 0.00E+00 | Paenibacillus macerans |
| 836 | AF-A0A1F5EWX9-F1-model_v4 SH3b domain-containing protein | 12.7 | 0.00E+00 | Candidatus Coatesbacteria bacterium RBG 13 66 14 |
| 837 | AF-A0A848G7Z8-F1-model_v4 Uncharacterized protein | 11.4 | 0.00E+00 | Zoogloea dura |
| 838 | AF-A0A7X8PIQ8-F1-model_v4 SH3 domain-containing protein | 9.7 | 0.00E+00 | Rhizobium sp. P28RR-XV |
| 839 | AF-A0A813FKE5-F1-model_v4 Hypothetical protein | 11.3 | 0.00E+00 | Polarella glacialis |
| 840 | AF-A0A533RDF4-F1-model_v4 SH3 domain-containing protein | 10.1 | 0.00E+00 | Deltaproteobacteria bacterium |

| | | | | |
|---|---|---|---|---|
| 841 | AF-A0A318STW3-F1-model_v4 SH3 domain-containing protein | 13.9 | 0.00E+00 | Pseudoroseicyclus aestuarii |
| 842 | AF-A0A1I2VJV0-F1-model_v4 SH3 domain-containing protein | 9.7 | 0.00E+00 | Desulfotruncus arcticus DSM 17038 |
| 843 | AF-A0A6B1E4S1-F1-model_v4 SH3 domain-containing protein | 9.2 | 0.00E+00 | Chloroflexi bacterium |
| 844 | AF-A0A329ZUS4-F1-model_v4 SH3b domain-containing protein | 9.8 | 0.00E+00 | Helicobacter sp. 15-1451 |
| 845 | AF-A0A0B0I0E4-F1-model_v4 Gamma-DL-glutamyl hydrolase | 9.4 | 0.00E+00 | Paenibacillus sp. P1XP2 |
| 846 | AF-A0A534HET5-F1-model_v4 SH3 domain-containing protein | 13.3 | 0.00E+00 | Gammaproteobacteria bacterium |
| 847 | AF-A0A3M1VR76-F1-model_v4 SH3b domain-containing protein | 11.6 | 0.00E+00 | Candidatus Dadabacteria bacterium |
| 848 | AF-A0A813GSS6-F1-model_v4 Hypothetical protein | 10.2 | 0.00E+00 | Polarella glacialis |
| 849 | AF-A0A3A9FGW4-F1-model_v4 NlpC/P60 domain-containing protein | 6.6 | 0.00E+00 | bacterium 1XD42-54 |
| 850 | AF-A0A2H6GSX2-F1-model_v4 SH3b domain-containing protein | 10.7 | 0.00E+00 | bacterium BMS3Abin15 |
| 851 | AF-A0A7W3VS06-F1-model_v4 SH3 domain-containing protein | 7.7 | 0.00E+00 | Amycolatopsis dendrobii |
| 852 | AF-A0A813I004-F1-model_v4 Hypothetical protein | 7.7 | 0.00E+00 | Polarella glacialis |
| 853 | AF-A0A6M8WYM7-F1-model_v4 SH3 domain-containing protein | 7 | 0.00E+00 | Rhizobium indicum |
| 854 | AF-A0A7X8E515-F1-model_v4 SH3 domain-containing protein | 13.6 | 0.00E+00 | Clostridiaceae bacterium |
| 855 | AF-A0A5E7YIF4-F1-model_v4 SH3b domain-containing protein | 11.9 | 0.00E+00 | Rhizobium sp. EC-SD404 |
| 856 | AF-A0A1G3L6X1-F1-model_v4 SH3b domain-containing protein | 14.4 | 0.00E+00 | Spirochaetes bacterium GWB1_36_13 |
| 857 | AF-A0A7S4Q1M1-F1-model_v4 Hypothetical protein | 11.7 | 0.00E+00 | Alexandrium monilatum |
| 858 | AF-A0A7S3LYQ9-F1-model_v4 Hypothetical protein | 13.4 | 0.00E+00 | Spumella elongata |
| 859 | AF-A0A1E3ZYK0-F1-model_v4 SH3b domain-containing protein | 7.7 | 0.00E+00 | Chryseobacterium sp. SCN 40-13 |
| 860 | AF-A0A8A7W9T5-F1-model_v4 SH3 domain-containing protein | 13.6 | 0.00E+00 | Cognatishimia activa |
| 861 | AF-A0A2W1JQ60-F1-model_v4 SH3b domain-containing protein | 14.2 | 0.00E+00 | Acaryochloris thomasi RCC1774 |
| 862 | AF-A0A813K1M9-F1-model_v4 Hypothetical protein | 8.4 | 0.00E+00 | Polarella glacialis |
| 863 | AF-B4WNY1-F1-model_v4 Bacterial SH3 domain family | 9.2 | 0.00E+00 | Synechococcus sp. PCC 7335 |
| 864 | AF-A0A2R6M791-F1-model_v4 Peptidase_M23 domain-containing protein | 8.5 | 0.00E+00 | Halobacteriales archaeon SW_6_65_15 |
| 865 | AF-A0A1Q6KU94-F1-model_v4 SH3b domain-containing protein | 11.8 | 0.00E+00 | Clostridiales bacterium 41_12_two_minus |
| 866 | AF-A0A2E4QSX2-F1-model_v4 SH3b domain-containing protein | 12.5 | 0.00E+00 | Hirschia sp. |
| 867 | AF-A0A1G7C0N9-F1-model_v4 DnaJ domain-containing protein | 14.1 | 0.00E+00 | Rhodospira trueperi |
| 868 | AF-A0A2J6WPV5-F1-model_v4 AAA domain-containing protein | 11.3 | 0.00E+00 | Calditerrivibrio nitroreducens |
| 869 | AF-A0A7Z9Y772-F1-model_v4 SH3b domain-containing protein | 14.2 | 0.00E+00 | Anaerolineae bacterium |
| 870 | AF-A0A853HZK9-F1-model_v4 Uncharacterized protein | 11.5 | 0.00E+00 | Azospirillum oleiclasticum |
| 871 | AF-A0A1G9BEW4-F1-model_v4 SH3 domain-containing protein | 9 | 0.00E+00 | Microbulbifer yueqingensis |
| 872 | AF-A0A812IXR3-F1-model_v4 Man1a1 protein | 12.7 | 0.00E+00 | Symbiodinium necroappetens |
| 873 | AF-A0A350VJR5-F1-model_v4 SH3 domain-containing protein | 10.3 | 0.00E+00 | Eubacterium sp. |
| 874 | AF-A0A1K1MDG6-F1-model_v4 SH3 domain-containing protein | 11.8 | 0.00E+00 | Ruminococcus sp. YE71 |
| 875 | AF-A0A5B8BTD9-F1-model_v4 SH3b domain-containing protein | 9.8 | 0.00E+00 | Oceanicola sp. D3 |

| | | | | |
|---|---|---|---|---|
| 876 | AF-A0A7V9HJI4-F1-model_v4 Transglycosylase SLT domain-containing protein | 10.4 | 0.00E+00 | Chloroflexia bacterium |
| 877 | AF-A0A1J0GIA6-F1-model_v4 SH3b domain-containing protein | 11.1 | 0.00E+00 | Clostridium estertheticum subsp. estertheticum |
| 878 | AF-A0A371XFQ2-F1-model_v4 Uncharacterized protein | 13.7 | 0.00E+00 | Mesorhizobium denitrificans |
| 879 | AF-A0A1U7NJQ3-F1-model_v4 Uncharacterized protein | 20 | 0.00E+00 | Dubosiella newyorkensis |
| 880 | AF-A0A5A7NTR3-F1-model_v4 SH3b domain-containing protein | 11.4 | 0.00E+00 | Zafaria cholistanensis |
| 881 | AF-A0A497CAM3-F1-model_v4 SH3b domain-containing protein | 14.7 | 0.00E+00 | Chloroflexi bacterium |
| 882 | AF-A0A176U9K3-F1-model_v4 N-acetylmuramoyl-L-alanine amidase | 10.1 | 0.00E+00 | Clostridiales bacterium KLE1615 |
| 883 | AF-A0A2N2JEJ5-F1-model_v4 Uncharacterized protein | 11.2 | 0.00E+00 | Deltaproteobacteria bacterium HGW-Deltaproteobacteria-14 |
| 884 | AF-A0A812T136-F1-model_v4 LanA protein | 6.8 | 0.00E+00 | Symbiodinium microadriaticum |
| 885 | AF-A0A7C4CT62-F1-model_v4 Uncharacterized protein | 12.2 | 0.00E+00 | Candidatus Bathyarchaeota archaeon |
| 886 | AF-A0A812P738-F1-model_v4 Epi-1 protein | 10.1 | 0.00E+00 | Symbiodinium sp. CCMP2456 |
| 887 | AF-A0A7W0TX77-F1-model_v4 SH3 domain-containing protein | 11.8 | 0.00E+00 | Chloroflexia bacterium |
| 888 | AF-A0A7S2AK42-F1-model_v4 Hypothetical protein | 11.3 | 0.00E+00 | Alexandrium andersonii |
| 889 | AF-A0A813I5N5-F1-model_v4 Hypothetical protein | 11.5 | 0.00E+00 | Polarella glacialis |
| 890 | AF-A0A7V8X5J9-F1-model_v4 SH3 domain-containing protein | 9.6 | 0.00E+00 | Chloroflexia bacterium |
| 891 | AF-A0A7W0Z100-F1-model_v4 SH3 domain-containing protein | 11.6 | 0.00E+00 | Chloroflexia bacterium |
| 892 | AF-A0A7C3DI05-F1-model_v4 SH3b domain-containing protein | 11.2 | 0.00E+00 | Chloroflexi bacterium |
| 893 | AF-A0A3T1D163-F1-model_v4 Uncharacterized protein | 16.7 | 0.00E+00 | Cohnella abietis |
| 894 | AF-A0A2D9YSD5-F1-model_v4 SH3b domain-containing protein | 15.4 | 0.00E+00 | Maritimibacter sp. |
| 895 | AF-A0A7S3JRA9-F1-model_v4 Hypothetical protein | 11.6 | 0.00E+00 | Aureoumbra lagunensis |
| 896 | AF-A0A6I4PWD5-F1-model_v4 Uncharacterized protein | 13.7 | 0.00E+00 | Actinomadura sp. J1-007 |
| 897 | AF-A0A1G8VMD2-F1-model_v4 Uncharacterized protein | 9.8 | 0.00E+00 | Actinopolyspora mzabensis |
| 898 | AF-A0A2G3DZR7-F1-model_v4 SH3b domain-containing protein | 11.1 | 0.00E+00 | Agathobacter ruminis |
| 899 | AF-A0A2B0DF68-F1-model_v4 SH3b domain-containing protein | 14.6 | 0.00E+00 | Bacillus thuringiensis |
| 900 | AF-A0A535BE20-F1-model_v4 Uncharacterized protein | 9.4 | 0.00E+00 | Deltaproteobacteria bacterium |
| 901 | AF-A6CP25-F1-model_v4 Uncharacterized protein | 10.5 | 0.00E+00 | Bacillus sp. SG-1 |
| 902 | AF-A0A4Q3U3N8-F1-model_v4 Phosphoribosylformylglycinamidine synthase | 9.6 | 0.00E+00 | bacterium |
| 903 | AF-A0A433DG22-F1-model_v4 NlpC/P60 domain-containing protein | 12.4 | 0.00E+00 | Jimgerdemannia flammicorona |
| 904 | AF-A0A7D3WSU0-F1-model_v4 DUF4468 domain-containing protein | 6.2 | 0.00E+00 | Hymenobacter sp. BRD67 |
| 905 | AF-A0A380WMW7-F1-model_v4 Uncharacterized protein conserved in bacteria | 14.2 | 0.00E+00 | Aminobacter aminovorans |
| 906 | AF-A0A812ZNX3-F1-model_v4 Hypothetical protein | 12 | 0.00E+00 | Symbiodinium microadriaticum |
| 907 | AF-A0A1I7LY33-F1-model_v4 SH3 domain-containing protein | 11.3 | 0.00E+00 | Methylobacterium sp. 174MFSha1.1 |
| 908 | AF-A0A7S1WF40-F1-model_v4 Hypothetical protein | 16.2 | 0.00E+00 | Alexandrium catenella |
| 909 | AF-A0A7S1REI9-F1-model_v4 Hypothetical protein | 14.4 | 0.00E+00 | Alexandrium catenella |

| | | | | |
|---|---|---|---|---|
| 910 | AF-A0A4R8GK07-F1-model_v4 S-layer family protein | 6.4 | 0.00E+00 | Cytobacillus oceanisediminis |
| 911 | AF-A0A7S2M273-F1-model_v4 Hypothetical protein | 9.3 | 0.00E+00 | Brandtodinium nutricula |
| 912 | AF-A0A7S3TFF0-F1-model_v4 Hypothetical protein | 15 | 0.00E+00 | Strombidinopsis acuminata |
| 913 | AF-A0A7S2WU80-F1-model_v4 Hypothetical protein | 13.7 | 0.00E+00 | Rhizochromulina marina |
| 914 | AF-A0A4Q1D3G9-F1-model_v4 SH3 domain-containing protein | 8.3 | 0.00E+00 | Filimonas effusa |
| 915 | AF-A0A6P0JS04-F1-model_v4 Uncharacterized protein | 12.5 | 0.00E+00 | Kamptonema sp. SIO4C4 |
| 916 | AF-A0A2V7UG09-F1-model_v4 SH3b domain-containing protein | 11.2 | 0.00E+00 | Acidobacteria bacterium |
| 917 | AF-A0A839ACU9-F1-model_v4 Uncharacterized protein | 16.3 | 0.00E+00 | Stappia albiluteola |
| 918 | AF-A0A426TSC7-F1-model_v4 Uncharacterized protein | 12.5 | 0.00E+00 | Candidatus Viridilinea halotolerans |
| 919 | AF-A0A351UQ41-F1-model_v4 Uncharacterized protein | 7.6 | 0.00E+00 | Elusimicrobia bacterium |
| 920 | AF-A0A2T3FVL9-F1-model_v4 LysM domain-containing protein | 14.1 | 0.00E+00 | Clostridium fessum |
| 921 | AF-A0A1I3NZP3-F1-model_v4 Uncharacterized protein | 12.1 | 0.00E+00 | Amycolatopsis sacchari |
| 922 | AF-A0A813IAP2-F1-model_v4 Hypothetical protein | 7 | 0.00E+00 | Polarella glacialis |
| 923 | AF-V7FC63-F1-model_v4 SH3b domain-containing protein | 12.7 | 0.00E+00 | Mesorhizobium sp. LSHC420B00 |
| 924 | AF-A0A7V5C439-F1-model_v4 Uncharacterized protein | 7.5 | 0.00E+00 | Campylobacterales bacterium |
| 925 | AF-A0A538AE49-F1-model_v4 SH3 domain-containing protein | 11.2 | 0.00E+00 | Actinomycetia bacterium |
| 926 | AF-E2CLX3-F1-model_v4 Putative DNA translocase FtsK | 14.8 | 0.00E+00 | Roseibium sp. TrichSKD4 |
| 927 | AF-R6Q7J0-F1-model_v4 SH3 type 3 domain protein | 8.8 | 0.00E+00 | Clostridium sp. CAG:508 |
| 928 | AF-A0A534UM86-F1-model_v4 Uncharacterized protein | 13 | 0.00E+00 | Deltaproteobacteria bacterium |
| 929 | AF-A0A6S5Y4C5-F1-model_v4 Uncharacterized protein | 3.1 | 0.00E+00 | Aeromonas veronii |
| 930 | AF-A0A7S3M1R5-F1-model_v4 Hypothetical protein | 8 | 0.00E+00 | Spumella elongata |
| 931 | AF-A0A7S2DC39-F1-model_v4 Hypothetical protein | 9 | 0.00E+00 | Alexandrium andersonii |
| 932 | AF-A0A535YGD0-F1-model_v4 Uncharacterized protein | 10.7 | 0.00E+00 | Chloroflexi bacterium |
| 933 | AF-A0A812UMZ9-F1-model_v4 Hypothetical protein | 8.4 | 0.00E+00 | Symbiodinium natans |
| 934 | AF-A0A2T6AS62-F1-model_v4 SH3 domain-containing protein | 11 | 0.00E+00 | Allosediminivita pacifica |
| 935 | AF-A0A2E0UYC8-F1-model_v4 Uncharacterized protein | 16.8 | 0.00E+00 | Anaerolineaceae bacterium |
| 936 | AF-A0A7W6W0I9-F1-model_v4 Uncharacterized protein YraI | 6.5 | 0.00E+00 | Rhodoblastus acidophilus |
| 937 | AF-A0A535XV03-F1-model_v4 Uncharacterized protein | 17 | 0.00E+00 | Chloroflexi bacterium |
| 938 | AF-A0A6P1HKM3-F1-model_v4 Polysaccharide deacetylase family protein | 10.5 | 0.00E+00 | Pontibacillus sp. HMF3514 |
| 939 | AF-A0A1E3BDQ2-F1-model_v4 Uncharacterized protein | 9.3 | 0.00E+00 | Aspergillus cristatus |
| 940 | AF-A0A292YCN7-F1-model_v4 SH3b domain-containing protein | 9.5 | 0.00E+00 | Effusibacillus lacus |
| 941 | AF-A0A534V8K9-F1-model_v4 Uncharacterized protein | 10.5 | 0.00E+00 | Deltaproteobacteria bacterium |
| 942 | AF-A0A1G2UR04-F1-model_v4 Uncharacterized protein | 12.3 | 0.00E+00 | Candidatus Zambryskibacteria bacterium RIFCSPLOWO2_12_FULL_39_16 |
| 943 | AF-A0A1J3FU41-F1-model_v4 F-box/kelch-repeat protein | 6 | 0.00E+00 | Noccaea caerulescens |

| | | | | |
|---|---|---|---|---|
| 944 | AF-A0A268NTD1-F1-model_v4 Uncharacterized protein | 14.2 | 0.00E+00 | Alkalihalobacillus clausii |
| 945 | AF-A0A7R6XFV2-F1-model_v4 Uncharacterized protein | 11.5 | 0.00E+00 | Mesorhizobium sp. 113-3-9 |
| 946 | AF-A0A291M1S9-F1-model_v4 SH3b domain-containing protein | 17.2 | 0.00E+00 | Pacificitalea manganoxidans |
| 947 | AF-A0A7W8XXS1-F1-model_v4 Uncharacterized protein YraI | 7.2 | 0.00E+00 | Rhizobium paranaense |
| 948 | AF-A0A4R1R6P1-F1-model_v4 N-acetylmuramoyl-L-alanine amidase | 12.8 | 0.00E+00 | Kineothrix alysoides |
| 949 | AF-A0A7X8HMX3-F1-model_v4 SH3 domain-containing protein | 11.9 | 0.00E+00 | Clostridiaceae bacterium |
| 950 | AF-A0A7S3U8H8-F1-model_v4 Hypothetical protein | 10.1 | 0.00E+00 | Strombidinopsis acuminata |
| 951 | AF-A0A3P1W3T9-F1-model_v4 SH3 domain-containing protein | 13.1 | 0.00E+00 | Tessaracoccus sp. OH4464_COT-324 |
| 952 | AF-A0A285NFB9-F1-model_v4 SH3 domain-containing protein | 12.5 | 0.00E+00 | Persephonella hydrogeniphila |
| 953 | AF-V6SR94-F1-model_v4 SH3b domain-containing protein | 12.2 | 0.00E+00 | Flavobacterium limnosediminis JC2902 |
| 954 | AF-A0A2T0RGI7-F1-model_v4 SH3 domain-containing protein | 13.7 | 0.00E+00 | Aliiruegeria haliotis |
| 955 | AF-A0P2R7-F1-model_v4 SH3b domain-containing protein | 11.6 | 0.00E+00 | Roseibium aggregatum IAM 12614 |
| 956 | AF-A0A7T1WSX1-F1-model_v4 Uncharacterized protein | 7.8 | 0.00E+00 | Streptomyces bathyalis |
| 957 | AF-A0A347UD67-F1-model_v4 SH3 domain-containing protein | 8.3 | 0.00E+00 | Profundibacter amoris |
| 958 | AF-A0A4P9K5Y9-F1-model_v4 Uncharacterized protein | 10.3 | 0.00E+00 | Thiomicrorhabdus sediminis |
| 959 | AF-A0A534SYV8-F1-model_v4 Uncharacterized protein | 12.6 | 0.00E+00 | Deltaproteobacteria bacterium |
| 960 | AF-A0A168W920-F1-model_v4 PPM-type phosphatase domain-containing protein | 9.7 | 0.00E+00 | Phormidium willei BDU 130791 |
| 961 | AF-A0A166FA04-F1-model_v4 Uncharacterized protein | 13.3 | 0.00E+00 | Methanobrevibacter filiformis |
| 962 | AF-A0A1E2RWU0-F1-model_v4 Bacterial SH3 domain protein | 7.9 | 0.00E+00 | Methyloligella halotolerans |
| 963 | AF-A0A1G2KQQ4-F1-model_v4 Uncharacterized protein | 7.7 | 0.00E+00 | Candidatus Sungbacteria bacterium RIFCSPHIGHO2_02_FULL_49_12 |
| 964 | AF-A0A350C1D0-F1-model_v4 SH3b domain-containing protein | 11.2 | 0.00E+00 | Lachnospiraceae bacterium |
| 965 | AF-A0A813KWR0-F1-model_v4 Hypothetical protein | 15.4 | 0.00E+00 | Polarella glacialis |
| 966 | AF-A0A812JMY5-F1-model_v4 LanA protein | 12.3 | 0.00E+00 | Symbiodinium necroappetens |
| 967 | AF-A0A1G5YQI8-F1-model_v4 SH3 domain-containing protein | 9.2 | 0.00E+00 | Sinorhizobium sp. NFACC03 |
| 968 | AF-A0A1B4XC20-F1-model_v4 Ligand-binding protein SH3 | 11 | 0.00E+00 | Sulfuricaulis limicola |
| 969 | AF-A0A812VDF6-F1-model_v4 Hypothetical protein | 11.1 | 0.00E+00 | Symbiodinium pilosum |
| 970 | AF-U4L097-F1-model_v4 Uncharacterized protein | 10.9 | 0.00E+00 | Pyronema omphalodes CBS 100304 |
| 971 | AF-A0A3M1QTB6-F1-model_v4 Uncharacterized protein | 7.5 | 0.00E+00 | Chloroflexi bacterium |
| 972 | AF-A0A2M7T5A6-F1-model_v4 SH3b domain-containing protein | 13.3 | 0.00E+00 | Candidatus Aquicultor secundus |
| 973 | AF-A0A257SJA9-F1-model_v4 NlpC/P60 domain-containing protein | 7 | 0.00E+00 | Gemmatimonadetes bacterium 21-71-4 |
| 974 | AF-A0A2N2PL36-F1-model_v4 Uncharacterized protein | 10 | 0.00E+00 | Chloroflexi bacterium HGW-Chloroflexi-10 |
| 975 | AF-A0A1T4V928-F1-model_v4 Cell wall-associated hydrolase, NlpC family | 10 | 0.00E+00 | Intestinibacter bartlettii DSM 16795 |
| 976 | AF-A0A535SX64-F1-model_v4 Transposase | 18.1 | 0.00E+00 | Chloroflexi bacterium |
| 977 | AF-A0A0F4VV57-F1-model_v4 Uncharacterized protein | 6.6 | 0.00E+00 | Clostridium sp. IBUN13A |

| | | | | |
|---|---|---|---|---|
| 978 | AF-A0A367UKI1-F1-model_v4 SH3b domain-containing protein | 13.3 | 0.00E+00 | Thalassospira xianhensis MCCC 1A02616 |
| 979 | AF-A0A813HET8-F1-model_v4 Hypothetical protein | 11.8 | 0.00E+00 | Polarella glacialis |
| 980 | AF-A0A151Z9E8-F1-model_v4 Lipase_3 domain-containing protein | 17.1 | 0.00E+00 | Tieghemostelium lacteum |
| 981 | AF-A0A433XL90-F1-model_v4 SH3 domain-containing protein | 11.9 | 0.00E+00 | Arsenicitalea aurantiaca |
| 982 | AF-A0A1H0Q646-F1-model_v4 Uncharacterized protein | 6.4 | 0.00E+00 | Actinokineospora alba |
| 983 | AF-A0A6T9AXX8-F1-model_v4 Hypothetical protein | 12 | 0.00E+00 | Alexandrium catenella |
| 984 | AF-A0A2V8MWX4-F1-model_v4 SH3b domain-containing protein | 9.7 | 0.00E+00 | Acidobacteria bacterium |
| 985 | AF-A0A527HQR6-F1-model_v4 DUF4236 domain-containing protein | 14.6 | 0.00E+00 | Mesorhizobium sp. |
| 986 | AF-A0A2X2JR55-F1-model_v4 Sensor protein fixL | 6.8 | 0.00E+00 | Sphingobacterium multivorum |
| 987 | AF-A0A2E4QEN0-F1-model_v4 Histidine kinase | 11.3 | 0.00E+00 | Legionellales bacterium |
| 988 | AF-A0A502FXG9-F1-model_v4 SH3b domain-containing protein | 11.4 | 0.00E+00 | Roseomonas nepalensis |
| 989 | AF-A0A7X4RTM2-F1-model_v4 SH3 domain-containing protein | 10.4 | 0.00E+00 | Vibrio sp. CAIM 722 |
| 990 | AF-A0A7S1LH19-F1-model_v4 Hypothetical protein | 6.1 | 0.00E+00 | Alexandrium catenella |
| 991 | AF-A0A812XKW3-F1-model_v4 LanA protein | 10 | 0.00E+00 | Symbiodinium sp. CCMP2592 |
| 992 | AF-A0A5R2N880-F1-model_v4 SH3 domain-containing protein | 14.8 | 0.00E+00 | Mesorhizobium sp. M2D.F.Ca.ET.145.01.1.1 |
| 993 | AF-A0A0R2AZS3-F1-model_v4 Glycosyl hydrolase family 25 | 11.2 | 0.00E+00 | Apilactobacillus ozensis DSM 23829 = JCM 17196 |
| 994 | AF-A0A7C6AKS8-F1-model_v4 Uncharacterized protein | 10.9 | 0.00E+00 | Candidatus Aminicenantes bacterium |
| 995 | AF-A0A1Q2YZZ5-F1-model_v4 Bacterial SH3 domain protein | 11.1 | 0.00E+00 | Roseomonas sp. TAS13 |
| 996 | AF-A0A1L8CK43-F1-model_v4 Bacterial SH3 domain protein | 10.5 | 0.00E+00 | Mariprofundus micogutta |
| 997 | AF-A0A351H816-F1-model_v4 Dockerin domain-containing protein | 15 | 0.00E+00 | Oscillospiraceae bacterium |
| 998 | AF-A0A3D1YY67-F1-model_v4 SH3b domain-containing protein | 13.6 | 0.00E+00 | Actinomycetia bacterium |
| 999 | AF-A0A3C1BHV6-F1-model_v4 HTH araC/xylS-type domain-containing protein | 9.8 | 0.00E+00 | Mucilaginibacter sp. |
| 1000 | AF-A0A6A6EQ92-F1-model_v4 Uncharacterized protein | 11.7 | 0.00E+00 | Zopfia rhizophila CBS 207.26 |
