## Supplemental Table S5 for "Amidase and Lysozyme Dual Functions in TseP Reveal a New Family of Chimeric Effectors in the Type VI Secretion System"

| Number | Target | Sequence identity | E-value | Scientific Name |
|---|---|---|---|---|
| 1 | AF-K1JFN7-F1-model_v4 EF-hand domain-containing protein | 100 | 3.09E-36 | Aeromonas dhakensis |
| 2 | AF-A0A7R8GJM6-F1-model_v4 EF-hand domain-containing protein | 70.9 | 1.11E-30 | Aeromonas jandaei |
| 3 | AF-A0A0A0CQD6-F1-model_v4 EF-hand domain-containing protein | 50 | 2.42E-23 | Photorhabdus luminescens |
| 4 | AF-A0A1B7L3A5-F1-model_v4 EF-hand domain-containing protein | 53.5 | 3.08E-23 | Mangrovibacter phragmitis |
| 5 | AF-A0A1G8DTN4-F1-model_v4 Predicted chitinase | 53.7 | 1.55E-22 | Vibrio xiamenensis |
| 6 | AF-A0A6B8NFJ9-F1-model_v4 Uncharacterized protein | 54.7 | 1.15E-22 | Vibrio sp. THAF191c |
| 7 | AF-A0A2K8QIQ6-F1-model_v4 Uncharacterized protein | 53.3 | 1.09E-21 | Dickeya fangzhongdai |
| 8 | AF-A0A318IY89-F1-model_v4 Putative chitinase | 49.3 | 4.99E-22 | Burkholderia pyrrocinia |
| 9 | AF-A0A1H6RCP4-F1-model_v4 Predicted chitinase | 51 | 8.53E-22 | Paraburkholderia diazotrophica |
| 10 | AF-A0A1H0VDX3-F1-model_v4 Predicted chitinase | 46 | 1.69E-21 | Ralstonia sp. 25mfcol4.1 |
| 11 | AF-A0A316IHM5-F1-model_v4 Putative chitinase | 51.4 | 2.27E-21 | Fulvimonas soli |
| 12 | AF-A0A6N4TYA1-F1-model_v4 Uncharacterized protein | 51.2 | 2.50E-21 | Burkholderia sp. THE68 |
| 13 | AF-A0A5D3CC78-F1-model_v4 Putative chitinase | 47.5 | 3.88E-21 | Cucumis melo var. makuwa |
| 14 | AF-A0A2X2DKC6-F1-model_v4 Glycoside hydrolase | 48.3 | 6.63E-21 | Pseudomonas luteola |
| 15 | AF-A0A2L0X3F1-F1-model_v4 Chitinase | 47.9 | 2.14E-20 | Cupriavidus metallidurans |
| 16 | AF-N2J2S6-F1-model_v4 Uncharacterized protein | 47.8 | 3.16E-20 | Pseudomonas sp. HPB0071 |
| 17 | AF-A0A5C1ZMT8-F1-model_v4 Calcium-binding protein | 49.5 | 5.69E-20 | Xanthomonas translucens pv. undulosa |
| 18 | AF-A0A0M3CVU4-F1-model_v4 EF-hand domain-containing protein | 46.8 | 2.02E-19 | Pseudomonas putida |
| 19 | AF-E5B1N5-F1-model_v4 Uncharacterized protein HI1415 | 53 | 1.22E-17 | Erwinia amylovora ATCC BAA-2158 |
| 20 | AF-A0A5J6QZL7-F1-model_v4 Uncharacterized protein | 46.7 | 3.98E-18 | Pseudomonas denitrificans (nom. rej.) |
| 21 | AF-A0A344KSE9-F1-model_v4 Lytic enzyme | 53.6 | 1.56E-17 | Vibrio campbellii |
| 22 | AF-A0A4R7V6P1-F1-model_v4 Putative chitinase | 42.3 | 5.88E-18 | Pseudomonas helmanticensis |
| 23 | AF-A0A7Y9WFN0-F1-model_v4 Putative chitinase | 41.8 | 2.21E-18 | Paraburkholderia bryophila |
| 24 | AF-A0A7Y8EKY7-F1-model_v4 Uncharacterized protein | 44.3 | 1.22E-17 | Pseudomonas yamanorum |
| 25 | AF-A0A7Y8FG31-F1-model_v4 EF-hand domain-containing protein | 44.2 | 1.99E-17 | Pseudomonas yamanorum |
| 26 | AF-A0A7X3HE31-F1-model_v4 Lytic enzyme | 44.7 | 3.58E-17 | Pseudomonas otitidis |
| 27 | AF-A0A4S3FRQ1-F1-model_v4 Lytic enzyme | 45.2 | 2.65E-16 | Massilia sp. Mn16-1_5 |
| 28 | AF-A0A318N7W5-F1-model_v4 Lytic enzyme | 39.1 | 1.79E-16 | Commensalibacter sp. ESL0284 |
| 29 | AF-A0A2D9Y7E2-F1-model_v4 PG_binding_1 domain-containing protein | 36.8 | 1.20E-14 | Candidatus Campbellbacteria bacterium |
| 30 | AF-A0A2G6GA07-F1-model_v4 PG_binding_1 domain-containing protein | 31.9 | 1.92E-12 | Candidatus Campbellbacteria bacterium |

| | | | | |
|---|---|---|---|---|
| 31 | AF-A0A4S4BC25-F1-model_v4 Glycoside hydrolase family 19 protein | 30.2 | 6.15E-11 | Deinococcus sp. Arct2-2 |
| 32 | AF-A0A7G6A3Q1-F1-model_v4 Uncharacterized protein | 35.4 | 4.19E-12 | Massilia sp. Se16.2.3 |
| 33 | AF-A0A354IRC3-F1-model_v4 Lytic enzyme | 46 | 6.45E-11 | Massilia sp. |
| 34 | AF-A0A1W1VAW0-F1-model_v4 Putative chitinase | 25.9 | 1.63E-10 | Deinococcus hopiensis KR-140 |
| 35 | AF-A0A368L8Z6-F1-model_v4 Glycoside hydrolase family 19 protein | 30 | 8.65E-11 | Parvibium lacunae |
| 36 | AF-A0A3S0JUC1-F1-model_v4 Glycoside hydrolase family 19 protein | 27.4 | 1.00E-10 | Deinococcus radiophilus |
| 37 | AF-F0EUW3-F1-model_v4 Uncharacterized protein | 31.7 | 2.31E-11 | Haemophilus parainfluenzae ATCC 33392 |
| 38 | AF-A0A0Q9EU01-F1-model_v4 Uncharacterized protein | 30.5 | 3.93E-10 | Lysobacter sp. Root96 |
| 39 | AF-A0A353FFS7-F1-model_v4 Glycoside hydrolase | 31.7 | 2.66E-10 | Cryomorphaceae bacterium |
| 40 | AF-A0A4R0XPM8-F1-model_v4 Glycoside hydrolase family 19 | 27.6 | 6.73E-10 | Paraburkholderia steynii |
| 41 | AF-A0A4T1DSL0-F1-model_v4 DNA primase | 32.6 | 4.78E-10 | Stenotrophomonas maltophilia |
| 42 | AF-A0A0C2U619-F1-model_v4 Lytic enzyme | 28.3 | 5.53E-10 | Magnetospirillum magnetotacticum MS-1 |
| 43 | AF-A0A323U6P3-F1-model_v4 Glycoside hydrolase family 19 protein | 30.3 | 6.10E-10 | Meiothermus sp. Pnk-1 |
| 44 | AF-A0A015Y6P9-F1-model_v4 Chitinase class I family protein | 33.1 | 3.56E-10 | Bacteroides fragilis str. J-143-4 |
| 45 | AF-A0A2V3V2M0-F1-model_v4 Putative chitinase | 28.1 | 8.58E-10 | Blastomonas natatoria |
| 46 | AF-A0A7X6NAS0-F1-model_v4 Type VI secretion system tip protein VgrG | 33 | 1.34E-10 | Vibrio aestuarianus |
| 47 | AF-A0A7Y5G1X0-F1-model_v4 Glycoside hydrolase family 19 protein | 27.9 | 2.79E-10 | candidate division KSB1 bacterium |
| 48 | AF-A0A3M7LQP5-F1-model_v4 Type VI secretion system tip protein VgrG | 31.9 | 5.06E-11 | Vibrio anguillarum |
| 49 | AF-A0A378RNQ1-F1-model_v4 Predicted chitinase | 31.2 | 6.40E-10 | Myroides odoratus |
| 50 | AF-A0A7K1SKT8-F1-model_v4 Glycoside hydrolase family 19 protein | 30.6 | 2.28E-09 | Spirosoma arboris |
| 51 | AF-A0A7Z7YGE7-F1-model_v4 Type VI secretion system tip protein VgrG | 32.8 | 1.28E-10 | Vibrio cholerae |
| 52 | AF-A0A523J2I7-F1-model_v4 Glycoside hydrolase family 19 protein | 28.8 | 1.27E-09 | Deltaproteobacteria bacterium |
| 53 | AF-A0A3A8QKE1-F1-model_v4 Glycoside hydrolase family 19 protein | 26 | 9.94E-10 | Corallococcus llansteffanensis |
| 54 | AF-A0A853I320-F1-model_v4 Glycoside hydrolase family 19 protein | 26 | 9.94E-10 | Endozoicomonas sp. SM1973 |
| 55 | AF-A0A317MB17-F1-model_v4 Putative chitinase | 28.6 | 8.58E-10 | Chitinophaga sp. S165 |
| 56 | AF-A0A375AFJ9-F1-model_v4 Phage endolysin | 29.3 | 2.07E-09 | Dickeya aquatica |
| 57 | AF-A0A117DPR9-F1-model_v4 Glycoside hydrolase family protein | 27.1 | 4.74E-09 | Deinococcus grandis |
| 58 | AF-A0A1N7K3K2-F1-model_v4 Putative chitinase | 29.5 | 1.04E-09 | Neptunomonas antarctica |

| | | | | |
|---|---|---|---|---|
| 59 | AF-A0A4U1BMN7-F1-model_v4 LysM peptidoglycan-binding domain-containing protein | 32.5 | 2.39E-09 | Ferrimonas aestuarii |
| 60 | AF-A0A5P8P3N5-F1-model_v4 Uncharacterized protein | 25 | 5.27E-10 | Sulfurimonas lithotrophica |
| 61 | AF-A0A250DS70-F1-model_v4 Glycoside hydrolase family 19 | 26.9 | 3.90E-09 | Variovorax boronicumulans |
| 62 | AF-N6Y999-F1-model_v4 Lytic enzyme | 25.6 | 2.91E-09 | Thauera sp. 27 |
| 63 | AF-A0A251ZSE0-F1-model_v4 Uncharacterized protein | 31.3 | 9.01E-10 | Commensalibacter intestini |
| 64 | AF-A0A2K8QRT5-F1-model_v4 Glycoside hydrolase family 19 | 26.4 | 5.53E-10 | Dickeya fangzhongdai |
| 65 | AF-W0EWB4-F1-model_v4 Lytic enzyme | 29 | 3.21E-09 | Niabella soli DSM 19437 |
| 66 | AF-A0A2E4QVS9-F1-model_v4 Glyco_hydro_19_cat domain-containing protein | 31.6 | 1.88E-09 | Hirschia sp. |
| 67 | AF-A0A849UDK0-F1-model_v4 Uncharacterized protein | 27.4 | 7.06E-10 | Methylococcaceae bacterium |
| 68 | AF-A0A3P3QTS3-F1-model_v4 LysM peptidoglycan-binding domain-containing protein | 25.9 | 9.39E-09 | Pararheinheimera mesophila |
| 69 | AF-A0A1H6AWQ2-F1-model_v4 Putative chitinase | 32.2 | 1.54E-09 | Algoriphagus boritolerans DSM 17298 = JCM 18970 |
| 70 | AF-A0A7W3AY70-F1-model_v4 Glycoside hydrolase family 19 protein | 30.7 | 2.91E-09 | Enterobacter hormaechei |
| 71 | AF-A0A839LDQ4-F1-model_v4 Glycoside hydrolase family 19 protein | 27.3 | 3.21E-09 | Mitsuaria sp. WAJ17 |
| 72 | AF-A0A2N1F5Z8-F1-model_v4 Glycoside hydrolase family 19 | 33.7 | 2.64E-09 | Tenacibaculum sp. Bg11-29 |
| 73 | AF-A0A350ZX06-F1-model_v4 Glyco_hydro_19_cat domain-containing protein | 29.2 | 1.88E-09 | Thauera sp. |
| 74 | AF-A0A0J1BVM8-F1-model_v4 Glyco_hydro_19_cat domain-containing protein | 31.6 | 1.97E-09 | Flavobacterium sp. ABG |
| 75 | AF-A0A143PMA6-F1-model_v4 Putative family GH19 chitinase | 33 | 3.37E-09 | Luteitalea pratensis |
| 76 | AF-A0A0X8GMH3-F1-model_v4 Uncharacterized protein | 27.4 | 5.49E-09 | Janthinobacterium sp. B9-8 |
| 77 | AF-A0A7U5E647-F1-model_v4 Abhydrolase_3 domain-containing protein | 30.8 | 3.23E-10 | Vibrio anguillarum |
| 78 | AF-A0A6A7N4M1-F1-model_v4 Glycoside hydrolase family 19 protein | 30.4 | 1.88E-09 | Rugamonas aquatica |
| 79 | AF-A0A554L8L8-F1-model_v4 Lytic enzyme | 25.6 | 4.52E-09 | Parcubacteria group bacterium Gr01-1014_3 |
| 80 | AF-U3CE46-F1-model_v4 LysM domain-containing protein | 32 | 6.36E-09 | Vibrio azureus NBRC 104587 |
| 81 | AF-A0A6A4UF91-F1-model_v4 Putative chitinase | 32.3 | 1.54E-09 | Ignavibacteria bacterium |
| 82 | AF-A0A855F1C3-F1-model_v4 Glycoside hydrolase family 19 | 28.5 | 4.30E-09 | Raoultella ornithinolytica |
| 83 | AF-A0A7U5EDT1-F1-model_v4 Type VI secretion protein VgrG | 29.6 | 8.18E-10 | Vibrio anguillarum |
| 84 | AF-A0A1K0JE46-F1-model_v4 Uncharacterized protein | 27.9 | 3.37E-09 | Cupriavidus necator |
| 85 | AF-A0A7X9I305-F1-model_v4 Glycoside hydrolase family 19 protein | 29.4 | 2.39E-09 | Serratia sp. |
| 86 | AF-A0A0U3LH66-F1-model_v4 Glycoside hydrolase family 19 | 26.5 | 1.97E-09 | Roseateles depolymerans |
| 87 | AF-A0A6L9LDJ3-F1-model_v4 Glycoside hydrolase family 19 protein | 28.9 | 2.17E-09 | Spirosoma terrae |

| | | | | |
|---|---|---|---|---|
| 88 | AF-A0A1S2N7B3-F1-model_v4 Chitinase class I family protein | 30.7 | 1.39E-08 | Massilia timonae |
| 89 | AF-C6E6S2-F1-model_v4 Lytic enzyme | 29.6 | 9.39E-09 | Geobacter sp. M21 |
| 90 | AF-A0A3D1J0H8-F1-model_v4 Glycoside hydrolase family 19 | 28.4 | 2.49E-08 | Oxalobacteraceae bacterium |
| 91 | AF-A0A508AAY6-F1-model_v4 Uncharacterized protein | 27.5 | 1.54E-09 | Lysobacter aestuarii |
| 92 | AF-A0A5M8R3E3-F1-model_v4 Glycoside hydrolase family 19 protein | 31.3 | 4.30E-09 | Dyadobacter flavalbus |
| 93 | AF-A0A564FWH6-F1-model_v4 Uncharacterized protein | 23.5 | 1.15E-09 | Methylobacterium dankookense |
| 94 | AF-A0A318KYE7-F1-model_v4 Putative chitinase | 32.3 | 1.46E-08 | Rivicola pingtungensis |
| 95 | AF-A0A844WUE5-F1-model_v4 Uncharacterized protein | 23.6 | 9.01E-10 | Gilliamella sp. Pas-s27 |
| 96 | AF-A0A0K1ECW6-F1-model_v4 Uncharacterized protein | 24.6 | 1.53E-08 | Chondromyces crocatus |
| 97 | AF-A0A0E3ZQW3-F1-model_v4 Putative phage lysozyme | 30.8 | 8.95E-09 | Pasteurella multocida subsp. multocida OH4807 |
| 98 | AF-A0A158DVF9-F1-model_v4 Glycoside hydrolase family protein | 25.4 | 1.04E-08 | Caballeronia pedi |
| 99 | AF-A0A1V3PSJ5-F1-model_v4 Glyco_hydro_19_cat domain-containing protein | 28.6 | 3.03E-08 | Rhodanobacter sp. B05 |
| 100 | AF-A0A8A6BH94-F1-model_v4 Glycoside hydrolase family 19 protein | 25.4 | 1.95E-08 | Pseudoalteromonas xiamenensis |
| 101 | AF-A0A2K4IBV5-F1-model_v4 Glycoside hydrolase family 19 | 28.8 | 1.39E-08 | Pseudomonas sp. MPR-ANC1 |
| 102 | AF-A0A443TVH8-F1-model_v4 EF-hand domain-containing protein | 28.2 | 5.76E-09 | Arcobacter venerupis |
| 103 | AF-A0A4Z1R3I3-F1-model_v4 Uncharacterized protein | 27.4 | 2.77E-09 | Luteimonas yindakuii |
| 104 | AF-A0A1I6HJ86-F1-model_v4 Putative chitinase | 28.2 | 3.21E-09 | Marinobacter daqiaonensis |
| 105 | AF-A0A7C9JNU0-F1-model_v4 Glycoside hydrolase family 19 protein | 27 | 3.18E-08 | Malikia spinosa |
| 106 | AF-A0A853HMK3-F1-model_v4 Glycoside hydrolase family 19 protein | 26.2 | 9.86E-09 | Azospirillum oleiclasticum |
| 107 | AF-A0A1T4SJ72-F1-model_v4 Putative chitinase | 30 | 6.05E-09 | Vibrio cincinnatiensis DSM 19608 |
| 108 | AF-A0A840MWB7-F1-model_v4 Putative chitinase | 25.6 | 4.98E-09 | Chitinivorax tropicus |
| 109 | AF-A0A4R8LQD6-F1-model_v4 Putative chitinase | 23.7 | 9.86E-09 | Paraburkholderia rhizosphaerae |
| 110 | AF-A0A4R2SFX2-F1-model_v4 Putative chitinase | 31.4 | 3.54E-09 | Cricetibacter osteomyelitidis |
| 111 | AF-N6XMG7-F1-model_v4 Uncharacterized protein | 27.4 | 2.17E-09 | Thauera sp. 27 |
| 112 | AF-A0A1H7YTM5-F1-model_v4 Predicted chitinase | 31.4 | 6.05E-09 | Chryseobacterium taichungense |
| 113 | AF-A0A0U5BUT1-F1-model_v4 Glycoside hydrolase family 19 protein | 23.3 | 3.03E-08 | Xanthomonas citri pv. citri |
| 114 | AF-A0A1U7P2X5-F1-model_v4 Phage endolysin | 29.3 | 1.86E-08 | Deinococcus marmoris |
| 115 | AF-A0A6J5F2F2-F1-model_v4 Uncharacterized protein | 28.6 | 3.21E-09 | Paraburkholderia humisilvae |
| 116 | AF-A0A017SZV3-F1-model_v4 Uncharacterized protein | 27.8 | 2.49E-08 | Chondromyces apiculatus DSM 436 |
| 117 | AF-A0A354V6S8-F1-model_v4 Glycoside hydrolase | 27 | 3.34E-08 | Acetobacteraceae bacterium |

| | | | | |
|---|---|---|---|---|
| 118 | AF-A0A0C2F229-F1-model_v4 Lytic enzyme | 31.4 | 4.48E-08 | Pseudomonas batumici |
| 119 | AF-A0A847NT58-F1-model_v4 Glycoside hydrolase family 19 protein | 28 | 1.39E-08 | Mollicutes bacterium |
| 120 | AF-A0A418VFK1-F1-model_v4 Uncharacterized protein | 24.6 | 2.25E-07 | Deinococcus cavernae |
| 121 | AF-A0A1I6G8C6-F1-model_v4 Putative chitinase | 29.9 | 2.91E-09 | Litoreibacter janthinus |
| 122 | AF-A0A4R2SJH7-F1-model_v4 Putative chitinase | 23.7 | 2.28E-09 | Cricetibacter osteomyelitidis |
| 123 | AF-A0A511NAD8-F1-model_v4 Uncharacterized protein | 21.4 | 3.18E-08 | Deinococcus cellulosilyticus NBRC 106333 = KACC 11606 |
| 124 | AF-A0A0S2F7C1-F1-model_v4 Chitinase class I family protein | 25.2 | 7.30E-08 | Lysobacter antibioticus |
| 125 | AF-A0A4R7BG38-F1-model_v4 Putative chitinase | 27.1 | 9.39E-09 | Paludibacterium purpuratum |
| 126 | AF-A0A4Q0UBC4-F1-model_v4 Uncharacterized protein | 26.3 | 1.69E-08 | Aliarcobacter skirrowii CCUG 10374 |
| 127 | AF-A0A853JGQ1-F1-model_v4 Uncharacterized protein | 25.4 | 4.27E-08 | Luteimonas salinisoli |
| 128 | AF-A0A226GZ78-F1-model_v4 Uncharacterized protein | 25.5 | 5.49E-09 | Flavobacterium hercynium |
| 129 | AF-A0A6G8NSN0-F1-model_v4 Uncharacterized protein | 23.5 | 1.32E-08 | Caballeronia sp. SBC1 |
| 130 | AF-A0A6V6Z7A3-F1-model_v4 Uncharacterized protein | 26.2 | 2.64E-09 | Flavobacterium salmonis |
| 131 | AF-A0A368P782-F1-model_v4 Glycoside hydrolase family 19 protein | 31.7 | 2.49E-08 | Oceanihabitans sediminis |
| 132 | AF-A0A266M011-F1-model_v4 Lytic enzyme | 26.9 | 1.32E-08 | Pseudomonas fragi |
| 133 | AF-A0A1M4VRU7-F1-model_v4 Putative chitinase | 30.5 | 1.20E-08 | Cnuella takakiae |
| 134 | AF-A0A0T2YIH3-F1-model_v4 Glyco_hydro_19_cat domain-containing protein | 26.6 | 2.75E-08 | Hydrogenophaga sp. Root209 |
| 135 | AF-A0A6L8KMT6-F1-model_v4 Glycoside hydrolase family 19 protein | 24.6 | 4.27E-08 | Duganella sp. CY15W |
| 136 | AF-A0A7W8WI83-F1-model_v4 Putative chitinase | 26.5 | 1.39E-08 | Paraburkholderia sp. WSM4180 |
| 137 | AF-A0A076J9D7-F1-model_v4 Uncharacterized protein | 27 | 5.76E-09 | Pseudomonas putida |
| 138 | AF-A0A217EGR4-F1-model_v4 Predicted chitinase | 25.4 | 1.04E-08 | Acinetobacter apis |
| 139 | AF-A0A1G0HY28-F1-model_v4 Glyco_hydro_19_cat domain-containing protein | 28.7 | 5.72E-08 | Gammaproteobacteria bacterium RIFCSPHIGHO2_12_FULL_63_22 |
| 140 | AF-A0A257S9C0-F1-model_v4 Glyco_hydro_19_cat domain-containing protein | 23.9 | 2.14E-07 | Acidiphilium sp. 21-60-14 |
| 141 | AF-A0A1L1PH70-F1-model_v4 Putative lysozyme | 22.8 | 5.19E-08 | Hydrogenophaga intermedia |
| 142 | AF-Q9RZ37-F1-model_v4 Glycohydrolase, putative | 22.8 | 2.75E-08 | Deinococcus radiodurans R1 |
| 143 | AF-A0A1T4Y0S9-F1-model_v4 Predicted chitinase | 25 | 2.26E-08 | Prosthecobacter debontii |
| 144 | AF-A0A1V3JP93-F1-model_v4 Uncharacterized protein | 27 | 5.49E-09 | Rodentibacter genomosp. 2 |
| 145 | AF-A0A0Q6S9Y5-F1-model_v4 Uncharacterized protein | 22 | 7.73E-09 | Massilia sp. Root335 |
| 146 | AF-A0A2E5ZN30-F1-model_v4 Uncharacterized protein | 27.4 | 1.08E-07 | Flavobacteriaceae bacterium |
| 147 | AF-A0A4Q8S0W8-F1-model_v4 EF-hand domain-containing protein | 24 | 1.14E-08 | Marinilabiliaceae bacterium JC017 |
| 148 | AF-A0A1I1X869-F1-model_v4 Predicted chitinase | 25.2 | 4.71E-08 | Nannocystis exedens |

| | | | | |
|---|---|---|---|---|
| 149 | AF-A0A3T0SBJ4-F1-model_v4 Uncharacterized protein | 26.8 | 4.74E-09 | Pseudomonadaceae bacterium SI-3 |
| 150 | AF-A0A1R1J8H9-F1-model_v4 Uncharacterized protein | 28.5 | 5.76E-09 | Burkholderia ubonensis |
| 151 | AF-A0A0B6FN09-F1-model_v4 Putative eF hand domain protein | 26.3 | 3.87E-08 | Yersinia frederiksenii Y225 |
| 152 | AF-K0JI41-F1-model_v4 Glycoside hydrolase family 19 | 28.2 | 3.51E-08 | Brachyspira pilosicoli WesB |
| 153 | AF-A0A855RPE9-F1-model_v4 Calcium-binding protein | 28.9 | 1.14E-08 | Photobacterium damselae subsp. damselae |
| 154 | AF-A0A2S0IC35-F1-model_v4 Uncharacterized protein | 27.5 | 2.26E-08 | Achromobacter spanius |
| 155 | AF-A0A2S6ZDT3-F1-model_v4 DNA primase | 26.3 | 3.51E-08 | Xanthomonas theicola |
| 156 | AF-G2HY68-F1-model_v4 Putative chitinase | 29.9 | 2.26E-08 | Arcobacter sp. L |
| 157 | AF-A0A5C0VK21-F1-model_v4 Glycoside hydrolase family 19 protein | 30 | 3.03E-08 | Pedobacter aquae |
| 158 | AF-A0A2N8QDG0-F1-model_v4 Uncharacterized protein | 26.3 | 1.32E-08 | Paraburkholderia fungorum |
| 159 | AF-A0A1G9XSX5-F1-model_v4 Predicted chitinase | 28.9 | 6.67E-09 | Stutzerimonas balearica DSM 6083 |
| 160 | AF-A0A6I6M6Y1-F1-model_v4 Glycoside hydrolase family 19 protein | 24.4 | 3.87E-08 | Pseudoduganella flava |
| 161 | AF-A0A0Q7R3Y0-F1-model_v4 Uncharacterized protein | 24.4 | 8.05E-08 | Brevundimonas sp. Root1423 |
| 162 | AF-A0A7X0F5F4-F1-model_v4 Putative chitinase | 27.3 | 8.05E-08 | Aminobacter aganoensis |
| 163 | AF-A0A7Y0ADD2-F1-model_v4 Uncharacterized protein | 24.5 | 8.05E-08 | Hymenobacter polaris |
| 164 | AF-A0A2N7W1U0-F1-model_v4 Uncharacterized protein | 27.7 | 7.73E-09 | Trinickia soli |
| 165 | AF-A0A1M4XY69-F1-model_v4 Predicted chitinase | 28.5 | 1.04E-08 | Chryseobacterium takakiae |
| 166 | AF-A0A2E4EDS5-F1-model_v4 Glycoside hydrolase | 32.3 | 9.79E-08 | Crocinitomicaceae bacterium |
| 167 | AF-F5RC29-F1-model_v4 Glyco_hydro_19_cat domain-containing protein | 24 | 7.67E-08 | Methyloversatilis universalis FAM5 |
| 168 | AF-A0A356JY00-F1-model_v4 Uncharacterized protein | 25.8 | 4.27E-08 | Treponema sp. |
| 169 | AF-F2NYH9-F1-model_v4 Uncharacterized protein | 24.2 | 6.62E-08 | Treponema succinifaciens DSM 2489 |
| 170 | AF-A0A504J3H1-F1-model_v4 Glycoside hydrolase family 19 protein | 26.4 | 8.45E-08 | Aquimarina algicola |
| 171 | AF-A0A1Q9QX61-F1-model_v4 Uncharacterized protein | 25.8 | 4.98E-09 | Pseudomonas putida |
| 172 | AF-A0A1I5UZK0-F1-model_v4 Putative chitinase | 32.4 | 8.05E-08 | Sphingomonas rubra |
| 173 | AF-A0A2N6FQ02-F1-model_v4 EF-hand domain-containing protein | 27.7 | 2.26E-08 | Arcobacter sp. |
| 174 | AF-A0A0N0JU86-F1-model_v4 Glyco_hydro_19_cat domain-containing protein | 30.9 | 1.52E-07 | beta proteobacterium AAP99 |
| 175 | AF-A0A3M8SXD4-F1-model_v4 PG_binding_1 domain-containing protein | 24.5 | 9.32E-08 | Lysobacter psychrotolerans |
| 176 | AF-A0A328T0D8-F1-model_v4 Uncharacterized protein | 25 | 9.39E-09 | Oleiagrimonas sp. MCCC 1A03011 |
| 177 | AF-A0A246BTH8-F1-model_v4 Uncharacterized protein | 22.8 | 1.03E-07 | Deinococcus indicus |
| 178 | AF-F6CWU7-F1-model_v4 Uncharacterized protein | 27.3 | 2.26E-08 | Marinomonas posidonica IVIA-Po-181 |

| | | | | |
|---|---|---|---|---|
| 179 | AF-A0A423P9F2-F1-model_v4 EF-hand domain-containing protein | 28.2 | 1.69E-08 | Pseudomonas fluorescens |
| 180 | AF-A0A7Z1KXB0-F1-model_v4 Glycoside hydrolase | 25 | 7.30E-08 | Janthinobacterium sp. BJB412 |
| 181 | AF-A0A2G0VN88-F1-model_v4 Chitinase | 27 | 6.36E-09 | Pseudomonas sp. ICMP 460 |
| 182 | AF-A0A2V6T5M4-F1-model_v4 Glycoside hydrolase family 19 | 27.1 | 1.03E-07 | Candidatus Rokubacteria bacterium |
| 183 | AF-A0A1X7I4L6-F1-model_v4 Putative chitinase | 25.6 | 5.45E-08 | Paraburkholderia susongensis |
| 184 | AF-A0A226X668-F1-model_v4 Lytic enzyme | 25.5 | 4.27E-08 | Caballeronia sordidicola |
| 185 | AF-A0A0K1ECJ4-F1-model_v4 Uncharacterized protein | 27.5 | 4.06E-08 | Chondromyces crocatus |
| 186 | AF-A0A2N4YG44-F1-model_v4 Phage_Mu_F domain-containing protein | 25.7 | 3.69E-08 | Tabrizicola sp. TH137 |
| 187 | AF-A0A0A1YIQ8-F1-model_v4 EF-hand domain-containing protein | 26.4 | 1.26E-08 | Pseudomonas taeanensis MS-3 |
| 188 | AF-A0A4V2F084-F1-model_v4 Putative chitinase | 22.8 | 4.03E-07 | Pseudobacter ginsenosidimutans |
| 189 | AF-A0A809H4M1-F1-model_v4 Uncharacterized protein | 25 | 5.96E-07 | [Mycobacterium] chelonae subsp. gwanakae |
| 190 | AF-A0A3A9VJR6-F1-model_v4 LysM peptidoglycan-binding domain-containing protein | 23.4 | 4.48E-08 | Aquimarina sp. AD10 |
| 191 | AF-A0A1M6E5F3-F1-model_v4 Putative chitinase | 23.9 | 6.62E-08 | Aureimonas altamirensis DSM 21988 |
| 192 | AF-A0A7Z2W023-F1-model_v4 Uncharacterized protein | 22.6 | 6.95E-08 | Massilia forsythiae |
| 193 | AF-A0A6I4NRL5-F1-model_v4 Uncharacterized protein | 21.9 | 1.53E-08 | Flavobacterium hydrocarbonoxydans |
| 194 | AF-A0A7Y2PYF2-F1-model_v4 Glycoside hydrolase family 19 | 26 | 1.95E-08 | Acinetobacter sp. ANC 5414 |
| 195 | AF-A0A4Z1AYU2-F1-model_v4 Uncharacterized protein | 21.6 | 1.53E-08 | Empedobacter tilapiae |
| 196 | AF-A0A1N6LPS1-F1-model_v4 Predicted chitinase | 26.4 | 2.05E-08 | Burkholderia sp. GAS332 |
| 197 | AF-A0A6J4EDA1-F1-model_v4 Uncharacterized protein | 26.4 | 1.32E-08 | Pseudomonas tohonis |
| 198 | AF-A0A2K9NI96-F1-model_v4 Endolysin | 26.7 | 6.01E-08 | Niveispirillum cyanobacteriorum |
| 199 | AF-A5FGP2-F1-model_v4 Glycoside hydrolase family 19 | 25.4 | 3.69E-08 | Flavobacterium johnsoniae UW101 |
| 200 | AF-A0A552U9X6-F1-model_v4 Glycoside hydrolase family 19 protein | 25.7 | 1.45E-07 | Glacieibacterium frigidum |
| 201 | AF-A0A3N0VLV6-F1-model_v4 Glycoside hydrolase family 19 protein | 24.7 | 1.76E-07 | Stagnimonas aquatica |
| 202 | AF-Q8ZQH4-F1-model_v4 Putative Fels-1 prophage chitinase | 28.3 | 7.30E-08 | Salmonella enterica subsp. enterica serovar Typhimurium str. LT2 |
| 203 | AF-Q87UT8-F1-model_v4 EF hand domain protein | 24.6 | 1.95E-08 | Pseudomonas syringae pv. tomato str. DC3000 |
| 204 | AF-A0A4P7A8R4-F1-model_v4 Glycoside hydrolase family 19 protein | 27.7 | 2.73E-07 | Herbaspirillum huttiense |
| 205 | AF-A0A7Z2Z4D7-F1-model_v4 EF-hand domain-containing protein | 22.2 | 5.76E-09 | Aeromonas hydrophila |
| 206 | AF-E0SF81-F1-model_v4 EF hand domain protein | 23.6 | 6.62E-08 | Dickeya dadantii 3937 |
| 207 | AF-A0A848FF36-F1-model_v4 Glycoside hydrolase family 19 protein | 20.1 | 3.01E-07 | Azohydromonas caseinilytica |

| | | | | |
|---|---|---|---|---|
| 208 | AF-A9MN02-F1-model_v4 EF-hand domain-containing protein | 25.7 | 9.32E-08 | Salmonella enterica subsp. arizonae serovar 62:z4,z23:- |
| 209 | AF-A0A6S6QM92-F1-model_v4 Uncharacterized protein | 22.2 | 2.36E-07 | Terrihabitans soli |
| 210 | AF-A0A4Q0UH99-F1-model_v4 Uncharacterized protein | 23.9 | 3.18E-08 | Arcobacter defluvii |
| 211 | AF-A0A0Q0AGB6-F1-model_v4 EF hand domain protein | 26.7 | 1.95E-08 | Pseudomonas syringae pv. solidagae |
| 212 | AF-A0A6I1U8Y1-F1-model_v4 Peptidoglycan DD-metalloendopeptidase family protein | 25.3 | 2.62E-08 | Pseudomonas sp. SZ57 |
| 213 | AF-A0A1M7L1E1-F1-model_v4 Putative chitinase | 30.6 | 5.19E-08 | Flavobacterium chilense |
| 214 | AF-A0A4Q0Y0Y7-F1-model_v4 Uncharacterized protein | 25.2 | 5.72E-08 | Halarcobacter anaerophilus |
| 215 | AF-A0A4P8HVY2-F1-model_v4 Putative chitinase | 25 | 6.01E-08 | Pseudoduganella umbonata |
| 216 | AF-A0A423ES90-F1-model_v4 EF-hand domain-containing protein | 22.8 | 1.61E-08 | Pseudomonas poae |
| 217 | AF-A0A1T1BSE0-F1-model_v4 Uncharacterized protein | 18.3 | 2.89E-08 | Flavobacterium sp. LM4 |
| 218 | AF-A0A1Y0M6L5-F1-model_v4 Uncharacterized protein | 23.7 | 9.32E-08 | Polaribacter sp. SA4-10 |
| 219 | AF-A0A1H4E446-F1-model_v4 Predicted chitinase | 23.3 | 5.45E-08 | Chitinophaga terrae (ex Kim and Jung 2007) |
| 220 | AF-A0A2W5ECW6-F1-model_v4 Glycohydrolase | 22.7 | 1.76E-07 | Pseudopedobacter saltans |
| 221 | AF-A0A5T8WQ64-F1-model_v4 Uncharacterized protein | 25.7 | 1.13E-07 | Salmonella enterica |
| 222 | AF-A0A7R7Z9H1-F1-model_v4 Uncharacterized protein | 24.4 | 4.71E-08 | Pseudomonas alcaligenes |
| 223 | AF-A0A660LXP1-F1-model_v4 Glycoside hydrolase family 19 protein | 33.1 | 2.60E-07 | Candidatus Saccharimonas sp. |
| 224 | AF-A0A7U9F5S2-F1-model_v4 Lytic enzyme | 29.9 | 2.48E-07 | Pseudomonas alcaligenes OT 69 |
| 225 | AF-A0A5E6P719-F1-model_v4 Glyco_hydro_19_cat domain-containing protein | 26.6 | 5.41E-07 | Pseudomonas fluorescens |
| 226 | AF-A0A679GDI2-F1-model_v4 Uncharacterized protein | 24.5 | 3.34E-08 | Pseudomonas otitidis |
| 227 | AF-A0A1H0WXC3-F1-model_v4 Predicted chitinase | 27.3 | 2.62E-08 | Pseudomonas guguanensis |
| 228 | AF-A0A7Y5I3X9-F1-model_v4 Glycoside hydrolase family 19 protein | 24.8 | 1.52E-07 | Pseudanabaena biceps |
| 229 | AF-A0A7W3U526-F1-model_v4 Glycoside hydrolase family 19 protein | 25.9 | 8.05E-08 | Lysobacter penaei |
| 230 | AF-A0A3G9FZA3-F1-model_v4 Phage protein | 23.8 | 9.79E-08 | Asticcacaulis excentricus |
| 231 | AF-A0A5C7BYA2-F1-model_v4 Uncharacterized protein | 22.9 | 4.94E-08 | Serratia marcescens |
| 232 | AF-A0A853JH95-F1-model_v4 Calcium-binding protein | 25.6 | 5.19E-08 | Luteimonas salinisoli |
| 233 | AF-A0A7W7NXQ4-F1-model_v4 Putative chitinase | 24.1 | 2.36E-07 | Novosphingobium chloroacetimidivorans |
| 234 | AF-A0A1Y1SYD5-F1-model_v4 Lytic enzyme | 28.7 | 9.32E-08 | Zunongwangia atlantica 22II14-10F7 |
| 235 | AF-J2LZI2-F1-model_v4 Putative chitinase | 28.5 | 1.31E-07 | Pantoea sp. GM01 |
| 236 | AF-A0A2L0EYF8-F1-model_v4 Uncharacterized protein | 22.3 | 2.14E-07 | Sorangium cellulosum |
| 237 | AF-A0A1I4ZAJ8-F1-model_v4 Predicted chitinase | 20.3 | 2.75E-08 | Pseudomonas sp. ok602 |
| 238 | AF-A0A1Z4F176-F1-model_v4 Lysozyme | 25 | 1.18E-06 | [Mycobacterium] stephanolepidis |

| | | | | |
|---|---|---|---|---|
| 239 | AF-A0A7D7VNR8-F1-model_v4 Uncharacterized protein | 24.7 | 4.94E-08 | Flavobacteriaceae bacterium |
| 240 | AF-A0A1I5W5V9-F1-model_v4 Predicted chitinase | 22.6 | 6.31E-08 | Pseudarcicella hirudinis |
| 241 | AF-A0A495GV59-F1-model_v4 Putative chitinase | 26.2 | 8.88E-08 | Paraburkholderia sediminicola |
| 242 | AF-A0A158L147-F1-model_v4 Glycoside hydrolase family protein | 24.2 | 6.31E-08 | Caballeronia arvi |
| 243 | AF-A0A0P0DZ97-F1-model_v4 Uncharacterized protein | 25.7 | 5.15E-07 | Sphingopyxis macrogoltabida |
| 244 | AF-A0A2N8EL48-F1-model_v4 EF-hand domain-containing protein | 24.3 | 3.87E-08 | Pseudomonas sp. FW305-131 |
| 245 | AF-A0A2P1VPN0-F1-model_v4 Uncharacterized protein | 21.8 | 1.86E-08 | Plesiomonas shigelloides |
| 246 | AF-A0A2S3WF94-F1-model_v4 Glycoside hydrolase family 19 | 25.5 | 6.26E-07 | Pseudomonas putida |
| 247 | AF-A0A7C9QT91-F1-model_v4 Glycoside hydrolase family 19 protein | 23.7 | 1.38E-07 | Magnetospirillum aberrantis SpK |
| 248 | AF-A0A4U9HCI0-F1-model_v4 Predicted chitinase | 26.2 | 4.71E-08 | Serratia rubidaea |
| 249 | AF-A0A4P7R235-F1-model_v4 LysM peptidoglycan-binding domain-containing protein | 24.6 | 1.85E-07 | Sphingomonas sp. PAMC26645 |
| 250 | AF-A0A651E1M0-F1-model_v4 Glycoside hydrolase family 19 protein | 24.1 | 1.44E-06 | Leptolyngbya sp. DLM2.Bin15 |
| 251 | AF-A0A497WSH5-F1-model_v4 Putative chitinase | 23.2 | 3.69E-08 | Pseudomonas asplenii |
| 252 | AF-K8B3G2-F1-model_v4 EF-hand domain-containing protein | 25.4 | 7.30E-08 | Cronobacter dublinensis 1210 |
| 253 | AF-A0A1M7DKY1-F1-model_v4 Predicted chitinase | 22.7 | 1.13E-07 | Chitinophaga jiangningensis |
| 254 | AF-A0A1I5MXM3-F1-model_v4 Putative chitinase | 23 | 4.45E-07 | Pseudarcicella hirudinis |
| 255 | AF-A0A086XRN2-F1-model_v4 Uncharacterized protein | 26.1 | 1.03E-07 | Paenirhodobacter enshiensis |
| 256 | AF-A0A653B4B2-F1-model_v4 Predicted chitinase | 20.7 | 3.87E-08 | Pseudomonas oleovorans |
| 257 | AF-C9QG57-F1-model_v4 Uncharacterized protein | 24.5 | 3.69E-08 | Vibrio orientalis CIP 102891 = ATCC 33934 |
| 258 | AF-A0A2U8FQ98-F1-model_v4 Glycoside hydrolase family 19 | 22 | 3.66E-07 | Aquabacterium olei |
| 259 | AF-A0A7X3H9U9-F1-model_v4 Glycoside hydrolase family 19 protein | 25.7 | 1.31E-07 | Pseudomonas otitidis |
| 260 | AF-A0A7K0JGR1-F1-model_v4 Uncharacterized protein | 23.1 | 2.60E-07 | Phocaeicola vulgatus |
| 261 | AF-A0A4Q1KDR4-F1-model_v4 Uncharacterized protein | 26.6 | 3.01E-07 | Flavobacterium stagni |
| 262 | AF-J2YI06-F1-model_v4 Putative chitinase | 24.3 | 4.71E-08 | Pseudomonas sp. GM24 |
| 263 | AF-A0A3G6TNN0-F1-model_v4 Peptidase_M23 domain-containing protein | 24.1 | 2.62E-08 | Chryseobacterium balustinum |
| 264 | AF-A0A2X3I6P2-F1-model_v4 Putative glycoside hydrolase | 32.1 | 2.73E-07 | Klebsiella pneumoniae |
| 265 | AF-A0A7V9DYG9-F1-model_v4 Glycoside hydrolase family 19 protein | 23.7 | 2.14E-07 | Burkholderiaceae bacterium |
| 266 | AF-A0A848HK97-F1-model_v4 Glycoside hydrolase family 19 protein | 23.4 | 6.95E-08 | Massilia polaris |
| 267 | AF-A0A2W6VLY9-F1-model_v4 Glycoside hydrolase family 19 | 24.4 | 1.37E-06 | Sphingomonas sp. |

| | | | | |
|---|---|---|---|---|
| 268 | AF-B4W0P6-F1-model_v4 Glyco_hydro_19_cat domain-containing protein | 24 | 3.84E-07 | Coleofasciculus chthonoplastes PCC 7420 |
| 269 | AF-A0A7C9M8B0-F1-model_v4 Glycoside hydrolase family 19 protein | 24.2 | 3.16E-07 | Deinococcus arboris |
| 270 | AF-A0A1L6I4B6-F1-model_v4 Uncharacterized protein | 28.7 | 6.95E-08 | Paraburkholderia sp. SOS3 |
| 271 | AF-A0A1W1UX33-F1-model_v4 Putative chitinase | 26.8 | 5.15E-07 | Deinococcus hopiensis KR-140 |
| 272 | AF-A0A423KEQ9-F1-model_v4 Uncharacterized protein | 20.3 | 1.13E-07 | Pseudomonas frederiksbergensis |
| 273 | AF-A0A7W4D9P1-F1-model_v4 Uncharacterized protein | 24.1 | 1.38E-07 | Pseudomonas guryensis |
| 274 | AF-A0A4Y8BR01-F1-model_v4 Glycoside hydrolase family 19 protein | 27.9 | 1.60E-07 | Campylobacter sp. US33a |
| 275 | AF-A0A496KQ03-F1-model_v4 LysM peptidoglycan-binding domain-containing protein | 25.5 | 1.68E-07 | Neisseria sp. |
| 276 | AF-A0A3N5XGV9-F1-model_v4 Glycoside hydrolase family 19 protein | 33.6 | 2.73E-07 | Dehalococcoidia bacterium |
| 277 | AF-A0A4R0PKP3-F1-model_v4 Type VI secretion system protein TssA | 20.4 | 3.18E-08 | Pseudomonas sp. IC_126 |
| 278 | AF-A0A209B2B6-F1-model_v4 Uncharacterized protein | 22.7 | 2.60E-07 | Yersinia frederiksenii |
| 279 | AF-A0A838L7R3-F1-model_v4 Glycoside hydrolase family 19 protein | 23.9 | 2.60E-07 | Sphingomonas chungangi |
| 280 | AF-A0A1C0B574-F1-model_v4 Uncharacterized protein | 27.3 | 8.88E-08 | Aliarcobacter thereius |
| 281 | AF-I3Z4M3-F1-model_v4 Putative chitinase | 28.4 | 1.94E-07 | Belliella baltica DSM 15883 |
| 282 | AF-I2B6W7-F1-model_v4 EF-hand domain-containing protein | 24 | 1.31E-07 | Shimwellia blattae DSM 4481 = NBRC 105725 |
| 283 | AF-A0A4S3YGT8-F1-model_v4 Chitinase | 25.2 | 7.67E-08 | Pseudomonas atacamensis |
| 284 | AF-A0A848HMP3-F1-model_v4 Glycoside hydrolase family 19 protein | 24.5 | 3.84E-07 | Massilia polaris |
| 285 | AF-A0A2A4R3G4-F1-model_v4 PG_binding_1 domain-containing protein | 24.3 | 6.57E-07 | Hyphomicrobiales bacterium |
| 286 | AF-A0A4P7WY14-F1-model_v4 Uncharacterized protein | 21.8 | 4.27E-08 | Psychroserpens sp. NJDZ02 |
| 287 | AF-A0A1G6IK64-F1-model_v4 Predicted chitinase | 21.3 | 5.72E-08 | Pseudomonas chengduensis |
| 288 | AF-A0A6B3UPH8-F1-model_v4 Glycoside hydrolase family 19 protein | 25.2 | 1.02E-06 | Caulobacter sp. 17J65-9 |
| 289 | AF-A0A1L7ALF8-F1-model_v4 Uncharacterized protein | 23.3 | 1.45E-07 | Roseomonas gilardii |
| 290 | AF-A0A3D9CMV8-F1-model_v4 Peptidase_M23 domain-containing protein | 24.7 | 4.27E-08 | Chryseobacterium flavum |
| 291 | AF-A0A2G3JEJ2-F1-model_v4 Lytic enzyme | 28 | 2.48E-07 | Iodobacter sp. BJB302 |
| 292 | AF-A0A6J5FH05-F1-model_v4 Glyco_hydro_19_cat domain-containing protein | 22.5 | 1.76E-07 | Paraburkholderia caffeinitolerans |
| 293 | AF-A0A7Z0P8Y2-F1-model_v4 Uncharacterized protein | 24.2 | 6.31E-08 | Pseudomonas sp. SbOxS1 |
| 294 | AF-A0A258YN27-F1-model_v4 Uncharacterized protein | 27.6 | 8.45E-08 | Sphingobacteriia bacterium 24-36-13 |
| 295 | AF-A0A1J5H1D3-F1-model_v4 Glyco_hydro_19_cat domain-containing protein | 24.1 | 2.02E-06 | Oscillatoriales cyanobacterium CG2_30_44_21 |

| | | | | |
|---|---|---|---|---|
| 296 | AF-B9USE1-F1-model_v4 Endolysin | 29 | 3.16E-07 | Brachyspira intermedia |
| 297 | AF-A0A329H6B8-F1-model_v4 Glycoside hydrolase family 19 protein | 26.9 | 2.48E-07 | Enterobacter sp. RIT 418 |
| 298 | AF-F2NE75-F1-model_v4 Peptidoglycan-binding domain 1 protein | 18 | 7.61E-07 | Desulfobacca acetoxidans DSM 11109 |
| 299 | AF-U2BA02-F1-model_v4 Uncharacterized protein | 20.4 | 4.71E-08 | Pseudomonas sp. EGD-AK9 |
| 300 | AF-A0A514C5T0-F1-model_v4 Glycoside hydrolase family 19 protein | 30.5 | 2.60E-07 | Brevundimonas sp. M20 |
| 301 | AF-A0A0X3AN89-F1-model_v4 Predicted chitinase | 24.5 | 1.68E-07 | Apibacter mensalis |
| 302 | AF-A0A2S7WMJ9-F1-model_v4 Uncharacterized protein | 25.4 | 6.57E-07 | Polaribacter porphyrae |
| 303 | AF-A0A1Z4JS80-F1-model_v4 Putative glycohydrolase | 25.1 | 6.95E-08 | Leptolyngbya boryana NIES-2135 |
| 304 | AF-A0A1V4LN17-F1-model_v4 Uncharacterized protein | 23.1 | 1.94E-07 | Pseudomonas sp. VI4.1 |
| 305 | AF-A0A246F5F1-F1-model_v4 Uncharacterized protein | 23.7 | 1.45E-07 | Pseudomonas nitroreducens |
| 306 | AF-A0A840C160-F1-model_v4 Putative chitinase | 22.7 | 5.96E-07 | Chelatococcus caeni |
| 307 | AF-A0A2V4AY63-F1-model_v4 Putative chitinase | 22.3 | 4.67E-07 | Prauserella muralis |
| 308 | AF-A0A0T9UF44-F1-model_v4 Putative endolysin | 24.4 | 1.60E-07 | Yersinia enterocolitica |
| 309 | AF-A9E9U9-F1-model_v4 Uncharacterized protein | 24.8 | 3.48E-07 | Kordia algicida OT-1 |
| 310 | AF-A0A350IBY7-F1-model_v4 Glycoside hydrolase family 19 | 30 | 1.02E-06 | Saprospirales bacterium |
| 311 | AF-A0A6J5EWQ2-F1-model_v4 Uncharacterized protein | 24.8 | 9.32E-08 | Paraburkholderia solisilvae |
| 312 | AF-A0A1A7KIB3-F1-model_v4 Peptidase family M23 | 22.8 | 6.62E-08 | Chryseobacterium sp. MOF25P |
| 313 | AF-A0A5B6TGS4-F1-model_v4 Uncharacterized protein | 24.1 | 4.67E-07 | Rufibacter hautae |
| 314 | AF-A0A0U1KI05-F1-model_v4 Putative endolysin | 24.2 | 5.15E-07 | Yersinia mollaretii |
| 315 | AF-A0A1H1X442-F1-model_v4 Predicted chitinase | 22.5 | 1.85E-07 | Pseudomonas umsongensis |
| 316 | AF-A0A2E8NUW9-F1-model_v4 Uncharacterized protein | 26.4 | 3.48E-07 | Euryarchaeota archaeon |
| 317 | AF-A0A1M4UIF0-F1-model_v4 Putative chitinase | 27.6 | 8.39E-07 | Dysgonomonas macrotermitis |
| 318 | AF-A0A7X9FIF0-F1-model_v4 Glycoside hydrolase family 19 protein | 22.4 | 2.48E-07 | Desulfovibrio sp. |
| 319 | AF-A0A5C7LW93-F1-model_v4 Glycoside hydrolase family 19 protein | 28.8 | 6.90E-07 | Hyphomicrobiaceae bacterium |
| 320 | AF-F9ZTP3-F1-model_v4 Chinitase domain protein | 25.2 | 1.60E-07 | Acidithiobacillus caldus SM-1 |
| 321 | AF-A0A0S4P9Y1-F1-model_v4 Chaperone protein DnaK | 30.7 | 2.04E-07 | Janthinobacterium sp. CG23_2 |
| 322 | AF-A0A430GYB8-F1-model_v4 Glycoside hydrolase family 19 protein | 26.3 | 1.37E-06 | Variovorax sp. 679 |
| 323 | AF-A0A4V2Q2X0-F1-model_v4 Putative chitinase | 24 | 3.16E-07 | Sodalis ligni |
| 324 | AF-A0A1G6MAT0-F1-model_v4 Putative chitinase | 21.1 | 3.81E-06 | Williamwhitmania taraxaci |
| 325 | AF-A0A4Q9VI24-F1-model_v4 Glycoside hydrolase family 19 protein | 21.9 | 4.24E-07 | Siculibacillus lacustris |

| | | | | |
|---|---|---|---|---|
| 326 | AF-A0A7Y0BYZ7-F1-model_v4 Glycoside hydrolase family 19 protein | 30.3 | 4.03E-07 | Polaromonas sp. |
| 327 | AF-A0A1H4AZB7-F1-model_v4 Predicted chitinase | 23.4 | 8.39E-07 | Bizionia paragorgiae |
| 328 | AF-A0A3G2IKU8-F1-model_v4 EF-hand domain-containing protein | 24.4 | 2.73E-07 | Buttiauxella sp. 3AFRM03 |
| 329 | AF-A0A7K1FYA1-F1-model_v4 Glyco_hydro_19_cat domain-containing protein | 21.7 | 1.52E-07 | Flavobacterium sp. LC2016-13 |
| 330 | AF-A0A352AQ06-F1-model_v4 DUF4231 domain-containing protein | 18.8 | 4.20E-06 | Cyanobacteria bacterium UBA9273 |
| 331 | AF-A0A127NVX2-F1-model_v4 Peptidase C39-like family protein | 19.3 | 2.12E-06 | Corynebacterium simulans |
| 332 | AF-A0A3D4WHL9-F1-model_v4 Uncharacterized protein | 24.6 | 2.73E-07 | Psychrobacter sp. |
| 333 | AF-L8N6P4-F1-model_v4 Glycoside hydrolase family 19 | 25.3 | 2.46E-06 | Pseudanabaena biceps PCC 7429 |
| 334 | AF-A0A078AT44-F1-model_v4 Glycoside hydrolase family protein | 20.7 | 5.92E-06 | Stylonychia lemnae |
| 335 | AF-A0A1S1HCN4-F1-model_v4 Glyco_hydro_19_cat domain-containing protein | 24 | 2.46E-06 | Sphingomonas haloaromaticamans |
| 336 | AF-A0A0F4NCW9-F1-model_v4 Calcium-binding protein | 25.1 | 3.01E-07 | Vibrio neptunius |
| 337 | AF-A0A6S6PBD3-F1-model_v4 Peptidase_M15_4 domain-containing protein | 18.9 | 2.23E-06 | Mycolicibacterium litorale |
| 338 | AF-A0A6L7IK48-F1-model_v4 Uncharacterized protein | 25.3 | 5.41E-07 | Flavobacteriaceae bacterium W22 |
| 339 | AF-A0A4D7QS29-F1-model_v4 Glycoside hydrolase family 19 protein | 28.9 | 2.23E-06 | Phreatobacter sp. NMCR1094 |
| 340 | AF-A0A8B4XK26-F1-model_v4 Putative chitinase | 24.4 | 5.37E-06 | Xanthomonas vasicola |
| 341 | AF-A0A3S4PHR5-F1-model_v4 Predicted chitinase | 25.7 | 3.48E-07 | Chryseobacterium gleum |
| 342 | AF-A0A327KQB0-F1-model_v4 Uncharacterized protein | 22.1 | 1.13E-06 | Rhodoplanes elegans |
| 343 | AF-A0A1M5CW97-F1-model_v4 Predicted chitinase | 22.7 | 2.87E-07 | Chryseobacterium arachidis |
| 344 | AF-A0A1S9B053-F1-model_v4 Uncharacterized protein | 25.6 | 1.18E-06 | Hymenobacter sp. CRA2 |
| 345 | AF-A0A540WMB1-F1-model_v4 Glycoside hydrolase family 19 protein | 20.1 | 5.63E-06 | Myxococcus llanfairpwllgwyngyllgogerychwyrndrobwllllantysiliogogogochensis |
| 346 | AF-A0A1B9M6M0-F1-model_v4 Uncharacterized protein | 22.4 | 1.68E-07 | Gilliamella apicola |
| 347 | AF-A0A1G5MGA6-F1-model_v4 Putative chitinase | 21.8 | 1.75E-06 | Afifella marina DSM 2698 |
| 348 | AF-A0A7D4BG23-F1-model_v4 Uncharacterized protein | 25.8 | 3.66E-07 | Tenuifilum thalassicum |
| 349 | AF-Q0RZW9-F1-model_v4 Peptidase_C39_2 domain-containing protein | 19.6 | 8.39E-07 | Rhodococcus jostii RHA1 |
| 350 | AF-A0A1E5QL45-F1-model_v4 Uncharacterized protein | 22.2 | 9.25E-07 | Desertifilum sp. IPPAS B-1220 |
| 351 | AF-A0A1G0TDC2-F1-model_v4 Uncharacterized protein | 30.2 | 9.71E-07 | Ignavibacteria bacterium RBG_16_35_7 |
| 352 | AF-A0A101CFM5-F1-model_v4 Uncharacterized protein | 26.3 | 9.71E-07 | Chryseobacterium aquaticum subsp. greenlandense |
| 353 | AF-A0A7D7ZQ66-F1-model_v4 Glyco_hydro_19_cat domain-containing protein | 26.1 | 4.67E-07 | Flavobacteriaceae bacterium |
| 354 | AF-A0A1N6EGG0-F1-model_v4 Predicted chitinase | 24.7 | 3.84E-07 | Paraburkholderia phenazinium |

| | | | | |
|---|---|---|---|---|
| 355 | AF-A0A250DIG0-F1-model_v4 Glyco_hydro_19_cat domain-containing protein | 25.3 | 1.92E-06 | Variovorax boronicumulans |
| 356 | AF-A0A4V1T5D2-F1-model_v4 Uncharacterized protein | 23.3 | 4.00E-06 | Sphingobacteriales bacterium |
| 357 | AF-A0A3S0B7A5-F1-model_v4 Glycoside hydrolase family 19 protein | 25.4 | 1.02E-06 | Rickettsiales bacterium |
| 358 | AF-A0A7K1WXG5-F1-model_v4 Uncharacterized protein | 22.7 | 5.96E-07 | Flavobacterium sp. HBTb2-11-1 |
| 359 | AF-A0A1V2W3A7-F1-model_v4 Glyco_hydro_19_cat domain-containing protein | 24.8 | 1.58E-06 | Burkholderia cenocepacia |
| 360 | AF-A0A1V2V447-F1-model_v4 Uncharacterized protein | 25.5 | 4.67E-07 | Acinetobacter genomosp. 33YU |
| 361 | AF-A0A1H8S4S8-F1-model_v4 Putative chitinase | 21.5 | 8.81E-07 | Rhodopseudomonas pseudopalustris |
| 362 | AF-A0A357KLE6-F1-model_v4 Uncharacterized protein | 22.7 | 5.96E-07 | Gammaproteobacteria bacterium |
| 363 | AF-A0A1E4LS01-F1-model_v4 Uncharacterized protein | 25.3 | 6.90E-07 | Acetobacteraceae bacterium SCN 69-10 |
| 364 | AF-A0A848GX15-F1-model_v4 Uncharacterized protein | 21.1 | 6.26E-07 | Chitinophaga fulva |
| 365 | AF-A0A212QPM0-F1-model_v4 Putative chitinase | 25.6 | 3.46E-06 | Arboricoccus pini |
| 366 | AF-V8QMZ9-F1-model_v4 Chitinase | 26.5 | 4.03E-07 | Advenella kashmirensis W13003 |
| 367 | AF-A0A1H2PSB3-F1-model_v4 Putative chitinase | 23.7 | 7.99E-07 | Chitinasiproducens palmae |
| 368 | AF-A0A4Q3NNW1-F1-model_v4 Glycoside hydrolase | 22.9 | 4.63E-06 | Cytophagaceae bacterium |
| 369 | AF-J0PVX3-F1-model_v4 Uncharacterized protein | 21.4 | 1.37E-06 | Bartonella sp. DB5-6 |
| 370 | AF-A0A3D5YT26-F1-model_v4 Uncharacterized protein | 22.9 | 1.44E-06 | Neisseriales bacterium |
| 371 | AF-A0A658BL16-F1-model_v4 Peptidoglycan-binding protein | 17.7 | 2.58E-06 | Nitrosomonadales bacterium |
| 372 | AF-F9ZTN9-F1-model_v4 Lytic enzyme | 19 | 3.66E-07 | Acidithiobacillus caldus SM-1 |
| 373 | AF-A0A0J5KRC0-F1-model_v4 Uncharacterized protein | 23 | 3.32E-07 | Pluralibacter gergoviae |
| 374 | AF-A0A704QWL6-F1-model_v4 Uncharacterized protein | 24.8 | 5.41E-07 | Salmonella enterica |
| 375 | AF-Q1QNE6-F1-model_v4 Putative glycohydrolase | 23.7 | 6.26E-07 | Nitrobacter hamburgensis X14 |
| 376 | AF-A0A2N9YH13-F1-model_v4 Peptidoglycan-binding protein | 20.8 | 3.63E-06 | Beggiatoa leptomitoformis |
| 377 | AF-A0A6A8QA39-F1-model_v4 Uncharacterized protein | 27.4 | 4.67E-07 | Balneolaceae bacterium |
| 378 | AF-A0A1E7YM65-F1-model_v4 Uncharacterized protein | 18.6 | 2.87E-07 | Acidithiobacillus caldus |
| 379 | AF-A0A6P0TAS0-F1-model_v4 DUF4231 domain-containing protein | 21.7 | 2.99E-06 | Cyanothece sp. SIO2G6 |
| 380 | AF-A0A1M3ECE7-F1-model_v4 Uncharacterized protein | 21.5 | 4.67E-07 | Bacteroidetes bacterium 43-16 |
| 381 | AF-A0A260UFF6-F1-model_v4 Uncharacterized protein | 20.9 | 4.63E-06 | Rhodococcus fascians |
| 382 | AF-A0A7T9GKP8-F1-model_v4 Glyco_hydro_19_cat domain-containing protein | 24.5 | 7.61E-07 | Acidobacteria bacterium |
| 383 | AF-A0A3D0HY99-F1-model_v4 Chitinase | 20.1 | 4.41E-06 | Bacteroidales bacterium |
| 384 | AF-A0A846FV64-F1-model_v4 Glycoside hydrolase family 19 protein | 23.8 | 5.37E-06 | Cyanothece sp. SIO1E1 |
| 385 | AF-A0A139A6M7-F1-model_v4 Glycoside hydrolase family 19 protein | 27.2 | 1.83E-06 | Gonapodya prolifera JEL478 |

| | | | | |
|---|---|---|---|---|
| 386 | AF-A0A1G5MIB4-F1-model_v4 Putative chitinase | 23.5 | 1.75E-06 | Afifella marina DSM 2698 |
| 387 | AF-A0A202C482-F1-model_v4 Glyco_hydro_19_cat domain-containing protein | 23.1 | 2.46E-06 | Chryseobacterium mucoviscidosis |
| 388 | AF-A0A1H4NU50-F1-model_v4 Putative chitinase | 23.6 | 1.17E-05 | Pseudomonas mohnii |
| 389 | AF-A0A158KAQ9-F1-model_v4 Glycoside hydrolase family protein | 23.3 | 5.96E-07 | Caballeronia choica |
| 390 | AF-A0A2H1YCJ6-F1-model_v4 Uncharacterized protein | 21.7 | 1.85E-07 | Tenacibaculum finnmarkense |
| 391 | AF-A0A1I5Z0M3-F1-model_v4 Predicted chitinase | 20.9 | 1.76E-07 | Pseudarcicella hirudinis |
| 392 | AF-A0A2N5DQ23-F1-model_v4 Uncharacterized protein | 23.9 | 2.71E-06 | Caulobacter zeae |
| 393 | AF-A0A355DZC0-F1-model_v4 Glyco_hydro_19_cat domain-containing protein | 19.5 | 2.99E-06 | Cyanobacteria bacterium UBA11162 |
| 394 | AF-A0A162DV08-F1-model_v4 Glyco_hydro_19_cat domain-containing protein | 20.7 | 3.63E-06 | Mycobacterium ostraviense |
| 395 | AF-A0A3B7BYW7-F1-model_v4 Uncharacterized protein | 19.8 | 1.30E-06 | Aquimarina sp. AD1 |
| 396 | AF-E9T0N0-F1-model_v4 Chitinase class I | 18.7 | 1.83E-06 | Prescottella equi ATCC 33707 |
| 397 | AF-A0A1Y6CZE3-F1-model_v4 Predicted chitinase | 23.4 | 2.12E-06 | Methylomagnum ishizawai |
| 398 | AF-A0A7I7QKE0-F1-model_v4 Glyco_hydro_19_cat domain-containing protein | 20.7 | 6.85E-06 | Mycolicibacterium sediminis |
| 399 | AF-A0A1Y5MSP7-F1-model_v4 Uncharacterized protein | 25.1 | 1.25E-07 | Campylobacter concisus |
| 400 | AF-A0A1C4CWH0-F1-model_v4 Predicted chitinase | 22.4 | 2.73E-07 | Gilliamella intestini |
| 401 | AF-A0A2S8ZL20-F1-model_v4 Uncharacterized protein | 22.1 | 2.87E-07 | Chryseobacterium sp. MYb7 |
| 402 | AF-A0A484GAH0-F1-model_v4 Uncharacterized protein | 23.2 | 4.90E-07 | Candidatus Schmidhempelia bombi str. Bimp |
| 403 | AF-A0A1M5LZH2-F1-model_v4 Predicted chitinase | 17.1 | 9.25E-07 | Flavobacterium sp. CF108 |
| 404 | AF-A0A431IZL5-F1-model_v4 Uncharacterized protein | 25.2 | 2.58E-06 | Flavobacteriaceae bacterium |
| 405 | AF-A0A2T5NWX7-F1-model_v4 NlpC/P60 domain-containing protein | 26 | 6.57E-07 | Chromobacterium haemolyticum |
| 406 | AF-A0A0Q6V2N2-F1-model_v4 Uncharacterized protein | 26.8 | 1.83E-06 | Caulobacter sp. Root343 |
| 407 | AF-A0A1A3PFI9-F1-model_v4 Glyco_hydro_19_cat domain-containing protein | 21.6 | 5.11E-06 | Mycobacterium sp. 1245111.1 |
| 408 | AF-A0A7H2V014-F1-model_v4 Glycoside hydrolase family 19 | 23.9 | 9.25E-07 | Acinetobacter seifertii |
| 409 | AF-A0A6P1D8A1-F1-model_v4 Glycoside hydrolase family 19 protein | 23.2 | 1.18E-06 | Nocardia cyriacigeorgica |
| 410 | AF-A0A4V2QF48-F1-model_v4 Putative chitinase | 26.1 | 7.25E-07 | Rhizobium sp. BK251 |
| 411 | AF-E8WJ98-F1-model_v4 Glycoside hydrolase family 19 | 23.4 | 7.93E-06 | Geobacter sp. M18 |
| 412 | AF-A0A5N7UTH2-F1-model_v4 Uncharacterized protein | 24.7 | 4.45E-07 | Chryseobacterium sp. |
| 413 | AF-A0A1B9M0Y9-F1-model_v4 Uncharacterized protein | 16.5 | 3.81E-06 | Gilliamella apicola |
| 414 | AF-R4USM3-F1-model_v4 Putative secreted chitinase | 22.2 | 3.14E-06 | Mycobacteroides abscessus subsp. bolletii 50594 |
| 415 | AF-A0A4U1IV12-F1-model_v4 Serine protease | 23.6 | 3.29E-06 | Polyangium fumosum |

| | | | | |
|---|---|---|---|---|
| 416 | AF-A0A5D4FNK5-F1-model_v4 Basic endochitinase | 18.8 | 4.20E-06 | Corynebacterium urealyticum |
| 417 | AF-K5BJE9-F1-model_v4 Peptidase C39 like family protein | 19.3 | 1.02E-06 | Mycolicibacterium hassiacum DSM 44199 |
| 418 | AF-A0A7S9UHP5-F1-model_v4 LysM peptidoglycan-binding domain-containing protein | 22.4 | 1.29E-05 | Massilia antarctica |
| 419 | AF-C1D6N5-F1-model_v4 Lytic enzyme | 23.4 | 3.29E-06 | Laribacter hongkongensis HLHK9 |
| 420 | AF-A0A736I8B2-F1-model_v4 Uncharacterized protein | 20.5 | 7.93E-06 | Salmonella enterica subsp. houtenae serovar 44:z36[z38]:- |
| 421 | AF-A0A6P2CMV5-F1-model_v4 Glyco_hydro_19_cat domain-containing protein | 21.3 | 1.06E-05 | Rhodococcus rhodnii |
| 422 | AF-A0A483LVB5-F1-model_v4 Uncharacterized protein | 24.7 | 3.46E-06 | Klebsiella pneumoniae |
| 423 | AF-A0A350Y8X9-F1-model_v4 Glyco_hydro_19_cat domain-containing protein | 19.7 | 4.87E-06 | Cyanobacteria bacterium UBA11370 |
| 424 | AF-A0A2S9CMW2-F1-model_v4 Uncharacterized protein | 25.1 | 9.71E-07 | Chryseobacterium culicis |
| 425 | AF-A0A504JD61-F1-model_v4 DUF4280 domain-containing protein | 23.8 | 4.67E-07 | Aquimarina algicola |
| 426 | AF-A0A850BKR8-F1-model_v4 Peptidoglycan-binding protein | 21.9 | 1.24E-06 | Polyangiaceae bacterium |
| 427 | AF-A0A1I5W332-F1-model_v4 Predicted chitinase | 22.9 | 1.18E-06 | Pseudarcicella hirudinis |
| 428 | AF-A0A379K5U9-F1-model_v4 Lysozyme | 27.2 | 4.87E-06 | Pseudomonas oleovorans |
| 429 | AF-A0A3D0WRG2-F1-model_v4 Chitinase | 23.8 | 2.34E-06 | Porphyromonadaceae bacterium |
| 430 | AF-J0QVH3-F1-model_v4 Uncharacterized protein | 21.8 | 2.84E-06 | Bartonella rattimassiliensis 15908 |
| 431 | AF-I3CCF5-F1-model_v4 Putative chitinase | 17.5 | 6.85E-06 | Beggiatoa alba B18LD |
| 432 | AF-A0A2S1YME2-F1-model_v4 Uncharacterized protein | 18.8 | 9.25E-07 | Flavobacterium crocinum |
| 433 | AF-A0A0Q4Y0P4-F1-model_v4 Glyco_hydro_19_cat domain-containing protein | 20.7 | 1.73E-05 | Pseudorhodoferax sp. Leaf267 |
| 434 | AF-A0A263D7I3-F1-model_v4 Glyco_hydro_19_cat domain-containing protein | 19.3 | 4.41E-06 | Amycolatopsis antarctica |
| 435 | AF-A0A516WS70-F1-model_v4 Glyco_hydro_19_cat domain-containing protein | 21.8 | 2.46E-06 | Rhodococcus sp. WB9 |
| 436 | AF-A0A3C0AWD5-F1-model_v4 Uncharacterized protein | 22.1 | 5.41E-07 | Flavobacterium sp. |
| 437 | AF-A0A2E9QPT1-F1-model_v4 Uncharacterized protein | 22.6 | 6.85E-06 | Deltaproteobacteria bacterium |
| 438 | AF-A0A0N9VBK8-F1-model_v4 Chitinase | 23.5 | 8.81E-07 | Acinetobacter equi |
| 439 | AF-A0A2K4MQ98-F1-model_v4 NlpC/P60 domain-containing protein | 23.3 | 3.14E-06 | Chromobacterium sinusclupearum |
| 440 | AF-D0SNX4-F1-model_v4 Uncharacterized protein | 22.1 | 2.34E-06 | Acinetobacter junii SH205 |
| 441 | AF-A0A0G2ZII7-F1-model_v4 Membrane-bound lytic murein transglycosylase D | 23.2 | 4.63E-06 | Archangium gephyra |
| 442 | AF-W0BP28-F1-model_v4 Chitinase | 23.3 | 2.99E-06 | Enterobacter ludwigii |
| 443 | AF-A0A353Q604-F1-model_v4 Chitinase | 25 | 2.34E-06 | Porphyromonadaceae bacterium |
| 444 | AF-A0A2T4VQY9-F1-model_v4 Peptidoglycan-binding protein | 23.5 | 5.63E-06 | Vitiosangium sp. GDMCC 1.1324 |
| 445 | AF-A0A1B9P0H2-F1-model_v4 Uncharacterized protein | 18.5 | 5.63E-06 | Aliivibrio logei |

| | | | | |
|---|---|---|---|---|
| 446 | AF-A0A255W849-F1-model_v4 Lysozyme | 23.2 | 4.63E-06 | Pseudomonas mandelii |
| 447 | AF-A0A2S2CKP1-F1-model_v4 Glycoside hydrolase family 19 | 28.6 | 2.34E-06 | Azospirillum thermophilum |
| 448 | AF-A0A4Y9R8T9-F1-model_v4 Glycoside hydrolase family 19 protein | 24.3 | 3.14E-06 | Brevundimonas sp. S30B |
| 449 | AF-A0A7V3F8N3-F1-model_v4 LysM peptidoglycan-binding domain-containing protein | 23.3 | 7.55E-06 | bacterium |
| 450 | AF-A0A1Y0G589-F1-model_v4 Uncharacterized protein | 24.8 | 5.63E-06 | Sulfuriferula sp. AH1 |
| 451 | AF-A0A1Q3HYL8-F1-model_v4 Glyco_hydro_19_cat domain-containing protein | 23.6 | 4.41E-06 | Archangium sp. Cb G35 |
| 452 | AF-A0A498CGP7-F1-model_v4 Putative chitinase | 23.5 | 5.92E-06 | Stenotrophomonas rhizophila |
| 453 | AF-A0A1I0LA10-F1-model_v4 Putative chitinase | 25.2 | 3.29E-06 | Stigmatella erecta |
| 454 | AF-A0A2I0FSK9-F1-model_v4 Uncharacterized protein | 26 | 6.85E-06 | Enterobacterales bacterium CwR94 |
| 455 | AF-A0A528X283-F1-model_v4 Peptidase_M15_4 domain-containing protein | 17.5 | 1.57E-05 | Mesorhizobium sp. |
| 456 | AF-A0A1D9G8U6-F1-model_v4 Uncharacterized protein | 17.4 | 9.64E-06 | Moorena producens JHB |
| 457 | AF-A0A355UQ03-F1-model_v4 LysM domain-containing protein | 18.9 | 3.43E-05 | Cyanobacteria bacterium UBA8530 |
| 458 | AF-A0A2A2GNR6-F1-model_v4 Serine protease | 17.8 | 1.43E-05 | Paracoccus salipaludis |
| 459 | AF-A0A2E9QWU6-F1-model_v4 Uncharacterized protein | 19.5 | 4.41E-06 | Deltaproteobacteria bacterium |
| 460 | AF-A0A1A7KI92-F1-model_v4 Peptidase family M23 | 21 | 1.92E-06 | Chryseobacterium sp. MOF25P |
| 461 | AF-A0A0N0LIQ6-F1-model_v4 Uncharacterized protein | 21.2 | 1.37E-06 | Chryseobacterium sp. ERMR1:04 |
| 462 | AF-A0A7Y4QXK8-F1-model_v4 Chitinase | 20.3 | 9.18E-06 | Ignavibacteria bacterium |
| 463 | AF-A0A0S3U6J5-F1-model_v4 Glycoside hydrolase, family 19 | 22 | 4.00E-06 | Leptolyngbya sp. NIES-3755 |
| 464 | AF-A0A4Q5YIK7-F1-model_v4 Glyco_hydro_19_cat domain-containing protein | 24.6 | 1.66E-06 | Chitinophagaceae bacterium |
| 465 | AF-A0A6L3SU12-F1-model_v4 Glycoside hydrolase family 19 protein | 26 | 6.85E-06 | Methylobacterium soli |
| 466 | AF-A0A7D4BZH6-F1-model_v4 Uncharacterized protein | 23 | 1.13E-06 | Tenuifilum thalassicum |
| 467 | AF-H1SB90-F1-model_v4 Chitinase-like protein | 24.5 | 2.34E-06 | Cupriavidus basilensis OR16 |
| 468 | AF-A0A840BFW7-F1-model_v4 Putative chitinase | 24 | 3.46E-06 | Niveibacterium umoris |
| 469 | AF-A0A7X1TW46-F1-model_v4 Chitinase | 20 | 8.74E-06 | Betaproteobacteria bacterium |
| 470 | AF-A0A346YKY4-F1-model_v4 Chitinase | 23.4 | 3.29E-06 | Neorhizobium sp. SOG26 |
| 471 | AF-A0A370H5R6-F1-model_v4 Putative chitinase | 24 | 3.81E-06 | Nocardia mexicana |
| 472 | AF-A0A519LH17-F1-model_v4 Peptidase_M23 domain-containing protein | 23.3 | 1.30E-06 | Flavobacterium sp. |
| 473 | AF-A0A5N5VKT7-F1-model_v4 Uncharacterized protein | 17.9 | 1.06E-05 | Mycolicibacterium mucogenicum DSM 44124 |
| 474 | AF-A0A484GEW4-F1-model_v4 Uncharacterized protein | 48.8 | 5.63E-06 | Candidatus Schmidhempelia bombi str. Bimp |
| 475 | AF-A0A495FX70-F1-model_v4 Putative chitinase | 21.1 | 1.73E-05 | Paraburkholderia sediminicola |
| 476 | AF-A0A736I4T9-F1-model_v4 Uncharacterized protein | 23.6 | 9.25E-07 | Salmonella enterica subsp. houtenae serovar 44:z36[z38]:- |

| | | | | |
|---|---|---|---|---|
| 477 | AF-A0A1B8R6J9-F1-model_v4 Phage endolysin | 21.9 | 3.78E-05 | Rhizobium leguminosarum bv. trifolii |
| 478 | AF-A0A5Y3Q770-F1-model_v4 Uncharacterized protein | 26 | 1.44E-06 | Salmonella enterica subsp. arizonae |
| 479 | AF-A0A1X0J2B1-F1-model_v4 Peptidase_M15_4 domain-containing protein | 21.3 | 1.06E-05 | Mycobacteroides saopaulense |
| 480 | AF-A0A7G5CV67-F1-model_v4 Uncharacterized protein | 24.3 | 1.43E-05 | Ewingella americana |
| 481 | AF-A0A257IX82-F1-model_v4 Fibronectin type-III domain-containing protein | 23.5 | 9.71E-07 | Cytophagaceae bacterium BCCC1 |
| 482 | AF-A0A7W9DH42-F1-model_v4 Putative chitinase | 24 | 2.01E-05 | Janthinobacterium sp. S3M3 |
| 483 | AF-A0A1V5ITE7-F1-model_v4 Chitinase class I | 22.1 | 3.81E-06 | Bacteroidetes bacterium ADurb.BinA104 |
| 484 | AF-A0A1I5Z243-F1-model_v4 Predicted chitinase | 20.5 | 4.90E-07 | Pseudarcicella hirudinis |
| 485 | AF-A0A838A087-F1-model_v4 Uncharacterized protein | 19.5 | 3.63E-06 | Mycobacteroides sp. LB1 |
| 486 | AF-A0A4Q3GEM4-F1-model_v4 Glycoside hydrolase family 19 protein | 23.9 | 2.32E-05 | Hyphomicrobiales bacterium |
| 487 | AF-A0A2S7FMZ0-F1-model_v4 Glyco_hydro_19_cat domain-containing protein | 28 | 4.87E-06 | Pseudomonas oleovorans |
| 488 | AF-A0A0P6VL12-F1-model_v4 PG_binding_1 domain-containing protein | 22.4 | 2.23E-06 | Prosthecomicrobium hirschii |
| 489 | AF-A0A7Y4PAF8-F1-model_v4 Basic endochitinase | 18 | 5.92E-06 | Corynebacterium ulcerans |
| 490 | AF-A0A246GFV0-F1-model_v4 DUF5675 domain-containing protein | 19.9 | 1.30E-06 | Flavobacterium columnare |
| 491 | AF-A0A482IP72-F1-model_v4 Chitinase | 19.9 | 8.74E-06 | Cupriavidus metallidurans |
| 492 | AF-A0A4C3GAJ5-F1-model_v4 Uncharacterized protein | 23.2 | 1.12E-05 | Escherichia coli |
| 493 | AF-A0A1M5NSK9-F1-model_v4 Putative chitinase | 17.2 | 7.55E-06 | Bradyrhizobium erythrophlei |
| 494 | AF-A0A5C7PXF4-F1-model_v4 Uncharacterized protein | 17.9 | 1.12E-05 | Desulfurellales bacterium |
| 495 | AF-A0A2T4VQQ4-F1-model_v4 Uncharacterized protein | 18.8 | 8.33E-06 | Vitiosangium sp. GDMCC 1.1324 |
| 496 | AF-A0A0X8WX03-F1-model_v4 Chitinase class I | 25.6 | 9.64E-06 | Leptolyngbya sp. O-77 |
| 497 | AF-A0A4R3EE59-F1-model_v4 Putative chitinase | 23.7 | 1.01E-05 | Bosea sp. BK604 |
| 498 | AF-A0A821EZP2-F1-model_v4 Hypothetical protein | 23.3 | 6.85E-06 | Rotaria sp. Silwood2 |
| 499 | AF-A0A4S5EDN9-F1-model_v4 Glyco_hydro_19_cat domain-containing protein | 18.2 | 5.11E-06 | Rhodococcus qingshengii |
| 500 | AF-A0A2A3DCW4-F1-model_v4 PG_binding_1 domain-containing protein | 21.3 | 9.64E-06 | Mesorhizobium sp. WSM3860 |
| 501 | AF-B1WPV6-F1-model_v4 Uncharacterized protein | 19.1 | 2.01E-05 | Crocosphaera subtropica ATCC 51142 |
| 502 | AF-A0A318UV35-F1-model_v4 Putative chitinase | 21.9 | 1.23E-05 | Pedobacter nutrimenti |
| 503 | AF-A0A3S5YGX0-F1-model_v4 Chitinase | 26.6 | 2.84E-06 | Salmonella enterica subsp. arizonae serovar 18:z4,z23:- str. CVM N26626 |
| 504 | AF-A0A085WWN3-F1-model_v4 Membrane-bound lytic murein transglycosylase D | 21.9 | 9.64E-06 | Hyalangium minutum |
| 505 | AF-A0A3M4KBA7-F1-model_v4 Prophage PSPPH01, chitinase | 22.5 | 6.21E-06 | Pseudomonas syringae pv. delphinii |

| | | | | |
|---|---|---|---|---|
| 506 | AF-G0HAJ7-F1-model_v4 Uncharacterized protein | 17.5 | 1.50E-05 | Corynebacterium variabile DSM 44702 |
| 507 | AF-A0A239GZY3-F1-model_v4 Putative chitinase | 20.1 | 7.19E-06 | Rhodococcus kyotonensis |
| 508 | AF-A0A426QY17-F1-model_v4 Glyco_hydro_19_cat domain-containing protein | 19.8 | 2.71E-06 | Rhodococcus sp. Eu-32 |
| 509 | AF-A0A6L3SZ36-F1-model_v4 PG_binding_1 domain-containing protein | 25.1 | 1.82E-05 | Methylobacterium soli |
| 510 | AF-A0A365XXD7-F1-model_v4 Uncharacterized protein | 23.9 | 3.63E-06 | Chitinophaga flava |
| 511 | AF-A0A077ZTD2-F1-model_v4 Glycoside hydrolase family protein | 23.5 | 5.37E-06 | Stylonychia lemnae |
| 512 | AF-A0A356KH37-F1-model_v4 Peptidoglycan-binding protein | 20 | 1.73E-05 | Planctomycetota bacterium |
| 513 | AF-A0A6A5BSH0-F1-model_v4 Glyco_hydro_19_cat domain-containing protein | 21.8 | 6.52E-06 | Naegleria fowleri |
| 514 | AF-A0A2W5VAY2-F1-model_v4 Uncharacterized protein | 24.1 | 2.01E-05 | Archangium gephyra |
| 515 | AF-A0A379SD19-F1-model_v4 Predicted chitinase | 21 | 3.27E-05 | Salmonella enterica |
| 516 | AF-A0A2G8MYJ5-F1-model_v4 Chitinase | 20 | 1.29E-05 | Pseudomonas sp. 382 |
| 517 | AF-A0A4U1I5T0-F1-model_v4 Peptidoglycan-binding protein | 17.2 | 2.11E-05 | Trinickia sp. 7GSK02 |
| 518 | AF-A0A5R1MUZ7-F1-model_v4 Basic endochitinase | 17.5 | 5.92E-06 | Corynebacterium silvaticum |
| 519 | AF-A0A3M1PG24-F1-model_v4 Glycoside hydrolase family 19 protein | 23.7 | 6.21E-06 | Cyanobacteria bacterium J069 |
| 520 | AF-A0A1C2DKX9-F1-model_v4 Glyco_hydro_19_cat domain-containing protein | 27.2 | 4.00E-06 | Stutzerimonas xanthomarina |
| 521 | AF-A0A1I0E9E3-F1-model_v4 Predicted chitinase | 25.4 | 8.74E-06 | Nitrosomonas marina |
| 522 | AF-W5XWT2-F1-model_v4 Uncharacterized protein | 17.6 | 1.23E-05 | Corynebacterium casei LMG S-19264 |
| 523 | AF-A0A4R5P3Y7-F1-model_v4 Uncharacterized protein | 19.8 | 3.63E-06 | Mycobacteroides franklinii |
| 524 | AF-L8M6U3-F1-model_v4 Putative chitinase | 19.9 | 2.11E-05 | Xenococcus sp. PCC 7305 |
| 525 | AF-A0A4V2NWL7-F1-model_v4 N-acetylmuramoyl-L-alanine amidase domain-containing protein | 24.6 | 2.34E-06 | Flaviaesturariibacter flavus |
| 526 | AF-A0A4R7ABN9-F1-model_v4 Putative chitinase | 20.1 | 3.60E-05 | Mycobacterium sp. OK889 |
| 527 | AF-A0A4R1L9F7-F1-model_v4 Putative chitinase | 22.2 | 2.32E-05 | Pusillimonas sp. YR330 |
| 528 | AF-A0A085WGA7-F1-model_v4 Membrane-bound lytic murein transglycosylase D | 22.8 | 1.17E-05 | Hyalangium minutum |
| 529 | AF-A0A3D0PCH5-F1-model_v4 VanY domain-containing protein | 17.7 | 2.44E-05 | Synechococcales bacterium UBA8647 |
| 530 | AF-A0A521FIZ6-F1-model_v4 Predicted chitinase | 20.9 | 2.71E-06 | Chryseobacterium rhizoplanae |
| 531 | AF-A0A226GZZ9-F1-model_v4 Uncharacterized protein | 23.2 | 2.71E-06 | Flavobacterium hercynium |
| 532 | AF-A0A316EER0-F1-model_v4 Putative chitinase | 20 | 1.57E-05 | Arcicella aurantiaca |
| 533 | AF-A0A084XW03-F1-model_v4 Putative chitinase | 18.8 | 3.43E-05 | Candidatus Accumulibacter vicinus |
| 534 | AF-A0A4R6SJA9-F1-model_v4 Putative chitinase | 22.1 | 8.33E-06 | Labedaea rhizosphaerae |
| 535 | AF-A0A6L7GBC7-F1-model_v4 Peptidoglycan-binding protein | 21.6 | 2.46E-06 | Pseudooceanicola sp. GBMRC 2024 |

| | | | | |
|---|---|---|---|---|
| 536 | AF-A0A1P8UWJ1-F1-model_v4 Putative chitinase | 22.9 | 4.17E-05 | Salipiger abyssi |
| 537 | AF-A0A1J5H5H1-F1-model_v4 Uncharacterized protein | 19.5 | 1.91E-05 | Oscillatoriales cyanobacterium CG2_30_44_21 |
| 538 | AF-A0A1Y6C0F2-F1-model_v4 Putative chitinase | 23.1 | 2.69E-05 | Pseudogulbenkiania subflava DSM 22618 |
| 539 | AF-A0A7W9UFL5-F1-model_v4 Putative chitinase | 21.4 | 8.74E-06 | Nocardia transvalensis |
| 540 | AF-A0A1B7L387-F1-model_v4 EF-hand domain-containing protein | 19.8 | 6.85E-06 | Mangrovibacter phragmitis |
| 541 | AF-A0A2P5LIT0-F1-model_v4 LysM domain-containing protein | 22 | 3.97E-05 | Methylobacter sp. |
| 542 | AF-A0A168W0Y5-F1-model_v4 Uncharacterized protein | 17.4 | 1.36E-05 | Phormidium willei BDU 130791 |
| 543 | AF-A0A800M4X9-F1-model_v4 Glycoside hydrolase family 19 protein | 23.9 | 6.85E-06 | Bacteroidetes bacterium |
| 544 | AF-T2GCV8-F1-model_v4 Putative baseplate assembly protein W | 21.3 | 1.12E-05 | Megalodesulfovibrio gigas DSM 1382 = ATCC 19364 |
| 545 | AF-A0A7C9Q0W9-F1-model_v4 Glyco_hydro_19_cat domain-containing protein | 19.6 | 5.07E-05 | Oscillatoria sp. SIO1A7 |
| 546 | AF-A0A4Y7Z1F9-F1-model_v4 Uncharacterized protein | 24.2 | 4.41E-06 | Escherichia coli |
| 547 | AF-I3PCM7-F1-model_v4 Putative lysozyme glycoside hydrolase family protein | 19.5 | 3.27E-05 | Variovorax sp. HH01 |
| 548 | AF-A0A3A8PYN9-F1-model_v4 Glycoside hydrolase family 19 protein | 22.6 | 9.18E-06 | Corallococcus aberystwythensis |
| 549 | AF-A0A5S4EJY3-F1-model_v4 Phage endolysin | 19.9 | 1.23E-05 | Candidatus Accumulibacter phosphatis |
| 550 | AF-A0A1Q6BRZ1-F1-model_v4 Peptidase_C39_2 domain-containing protein | 19.9 | 3.11E-05 | Corynebacterium glutamicum |
| 551 | AF-A0A1I3IQ17-F1-model_v4 Uncharacterized protein | 21.9 | 4.87E-06 | Halpernia frigidisoli |
| 552 | AF-W6RS59-F1-model_v4 Putative chitinase | 22.7 | 5.07E-05 | Rhizobium favelukesii |
| 553 | AF-A0A7Z1V637-F1-model_v4 Glycoside hydrolase family 19 | 22.2 | 2.69E-05 | Azospirillum brasilense |
| 554 | AF-A0A285UXT9-F1-model_v4 Putative chitinase | 22.7 | 5.59E-05 | Rhizobium subbaraonis |
| 555 | AF-D2VMQ6-F1-model_v4 Predicted protein | 21.8 | 1.17E-05 | Naegleria gruberi |
| 556 | AF-A0A679JU29-F1-model_v4 PG_binding_1 domain-containing protein | 16.1 | 4.41E-06 | Hyphomicrobium sp. ghe19 |
| 557 | AF-A0A1M6PHN5-F1-model_v4 Predicted chitinase | 18.8 | 9.64E-06 | Chryseobacterium polytrichastri |
| 558 | AF-D2VBA0-F1-model_v4 Predicted protein | 19.2 | 8.74E-06 | Naegleria gruberi |
| 559 | AF-A0A2N3XZ01-F1-model_v4 Putative chitinase | 18.4 | 4.41E-06 | Saccharopolyspora spinosa |
| 560 | AF-A0A7R7AMQ5-F1-model_v4 Uncharacterized protein | 23.8 | 5.92E-06 | Mesorhizobium sp. 113-1-2 |
| 561 | AF-A0A7W0UYF9-F1-model_v4 Peptidoglycan-binding protein | 22.5 | 2.01E-05 | Pyrinomonadaceae bacterium |
| 562 | AF-A0A1W9I729-F1-model_v4 PG_binding_1 domain-containing protein | 28.1 | 2.11E-05 | Proteobacteria bacterium HN_bin10 |
| 563 | AF-K1RJM4-F1-model_v4 Lysozyme | 22.1 | 4.83E-05 | human gut metagenome |
| 564 | AF-A0A7V8ZJI7-F1-model_v4 Glycoside hydrolase family 19 protein | 26.6 | 8.26E-05 | Solirubrobacterales bacterium |

| | | | | |
|---|---|---|---|---|
| 565 | AF-A0A6A5BVE6-F1-model_v4 Glyco_hydro_19_cat domain-containing protein | 21 | 9.64E-06 | Naegleria fowleri |
| 566 | AF-A0A2I8QMF6-F1-model_v4 Glycoside hydrolase | 21.5 | 1.50E-05 | Enterobacteriaceae bacterium ENNIH2 |
| 567 | AF-A0A1W9HC14-F1-model_v4 Uncharacterized protein | 20.1 | 9.18E-06 | Proteobacteria bacterium SG_bin6 |
| 568 | AF-A0A1H8J8K5-F1-model_v4 Putative chitinase | 23.5 | 4.20E-06 | Nitrosomonas marina |
| 569 | AF-A0A3D9AZT6-F1-model_v4 Uncharacterized protein | 20.7 | 3.63E-06 | Chryseobacterium piscium |
| 570 | AF-A0A370I0Y1-F1-model_v4 Putative chitinase | 23.7 | 2.32E-05 | Nocardia pseudobrasiliensis |
| 571 | AF-A0A1B1ALW6-F1-model_v4 Uncharacterized protein | 28.1 | 2.23E-06 | Candidatus Viadribacter manganicus |
| 572 | AF-A0A1S1LIL2-F1-model_v4 Chitinase | 20 | 2.44E-05 | Mycobacteroides chelonae |
| 573 | AF-A0A6H1TWP9-F1-model_v4 Peptidase_C39_2 domain-containing protein | 18.4 | 2.56E-05 | Oxynema aestuarii AP17 |
| 574 | AF-A3YYS2-F1-model_v4 Putative peptidoglycan binding domain 1 | 16.8 | 3.97E-05 | Synechococcus sp. WH 5701 |
| 575 | AF-A0A817M124-F1-model_v4 Hypothetical protein | 23.4 | 2.32E-05 | Rotaria sp. Silwood2 |
| 576 | AF-A0A7D7V7T3-F1-model_v4 Uncharacterized protein | 18.8 | 1.82E-05 | Gordonia rubripertincta |
| 577 | AF-A0A1Y0G6Y8-F1-model_v4 Uncharacterized protein | 25 | 5.11E-06 | Sulfuriferula sp. AH1 |
| 578 | AF-A0A7I7R8A7-F1-model_v4 Bacteriophage protein | 20.6 | 3.60E-05 | Mycolicibacter minnesotensis |
| 579 | AF-A0A847SJR9-F1-model_v4 DUF4280 domain-containing protein | 23.6 | 1.24E-06 | Chitinophaga eiseniae |
| 580 | AF-L7UBP9-F1-model_v4 Class I chitinase | 19.5 | 9.57E-05 | Myxococcus stipitatus DSM 14675 |
| 581 | AF-A0A1L6LFA3-F1-model_v4 Membrane-bound lytic murein transglycosylase D | 21.1 | 1.57E-05 | Minicystis rosea |
| 582 | AF-A0A1I5X9C9-F1-model_v4 Putative chitinase | 22.7 | 9.64E-06 | Parafilimonas terrae |
| 583 | AF-A0A1E4DMI0-F1-model_v4 Uncharacterized protein | 18.8 | 2.82E-05 | Methylobacterium sp. SCN 67-24 |
| 584 | AF-A0A375YXX6-F1-model_v4 Peptidase_C39_2 domain-containing protein | 20.2 | 3.11E-05 | Mycobacterium shimoidei |
| 585 | AF-A0A5B1B106-F1-model_v4 Uncharacterized protein | 20.9 | 7.19E-06 | Aquimarina sp. RZ0 |
| 586 | AF-A0A380CEM2-F1-model_v4 Predicted chitinase | 26 | 7.55E-06 | Sphingobacterium spiritivorum |
| 587 | AF-A0A6P0YX59-F1-model_v4 Peptidoglycan-binding protein | 17 | 6.17E-05 | Merismopedia sp. SIO2A8 |
| 588 | AF-Q1CVS9-F1-model_v4 Chitinase, class I | 20.6 | 9.64E-06 | Myxococcus xanthus DK 1622 |
| 589 | AF-A0A8B2BSU1-F1-model_v4 Uncharacterized protein | 20.8 | 6.85E-06 | Prevotella melaninogenica |
| 590 | AF-A0A1M4W6B2-F1-model_v4 Predicted chitinase | 20.8 | 6.85E-06 | Chryseobacterium vrystaatense |
| 591 | AF-A0A1V4XWR3-F1-model_v4 Uncharacterized protein | 24.6 | 1.82E-05 | Syntrophus sp. PtaB.Bin001 |
| 592 | AF-A0A3A8GR70-F1-model_v4 Glycoside hydrolase family 19 protein | 23.2 | 1.36E-05 | Corallococcus sp. AB011P |
| 593 | AF-A0A6M0A8T4-F1-model_v4 Glycoside hydrolase family 19 protein | 17.9 | 1.12E-05 | Okeania sp. SIO3B5 |
| 594 | AF-A0A0M6Y9U5-F1-model_v4 Putative chitinase | 19.7 | 1.23E-05 | Roseibium aggregatum |

| | | | | |
|---|---|---|---|---|
| 595 | AF-A0A6A5B8Y5-F1-model_v4 SH3b domain-containing protein | 19.5 | 1.23E-05 | Naegleria fowleri |
| 596 | AF-A0A444VY26-F1-model_v4 Glycoside hydrolase | 21.6 | 1.36E-05 | Flavobacterium anhuiense |
| 597 | AF-A0A1L7NPW5-F1-model_v4 Lytic enzyme | 26.3 | 1.91E-05 | Pseudomonas putida |
| 598 | AF-A0A840MZV1-F1-model_v4 Putative chitinase | 22.6 | 4.83E-05 | Chitinivorax tropicus |
| 599 | AF-A0A1B3NLT1-F1-model_v4 Peptidoglycan-binding 1 domain protein | 18.3 | 1.12E-05 | Bosea sp. RAC05 |
| 600 | AF-A0A7G9QMZ5-F1-model_v4 Uncharacterized protein | 22.2 | 1.44E-06 | Pedobacter roseus |
| 601 | AF-A0A101CQZ3-F1-model_v4 DUF5675 domain-containing protein | 23 | 6.52E-06 | Flavobacteriaceae bacterium CRH |
| 602 | AF-A0A2A2B7U1-F1-model_v4 LysM domain-containing protein | 23.1 | 1.17E-05 | Psychrobacter sp. JB193 |
| 603 | AF-A0A3C0XAC4-F1-model_v4 Glyco_hydro_19_cat domain-containing protein | 23.6 | 1.57E-05 | Stenotrophomonas sp. |
| 604 | AF-A0A2E9AX19-F1-model_v4 Uncharacterized protein | 21.6 | 2.21E-05 | Pseudomonas sp. |
| 605 | AF-A0A504BUZ6-F1-model_v4 Glycoside hydrolase family 19 protein | 21.4 | 8.68E-05 | Mesorhizobium sp. B2-1-3A |
| 606 | AF-A0A227ND01-F1-model_v4 Uncharacterized protein | 24 | 1.57E-05 | Flavobacterium araucananum |
| 607 | AF-A0A7Z0CY35-F1-model_v4 Putative chitinase | 17.8 | 1.23E-05 | Microlunatus terrae |
| 608 | AF-A0A3N2F1S8-F1-model_v4 Putative chitinase | 20.2 | 6.52E-06 | Chryseobacterium nakagawai |
| 609 | AF-K8GP44-F1-model_v4 Putative chitinase | 21 | 1.22E-04 | Leptolyngbyaceae cyanobacterium JSC-12 |
| 610 | AF-A0A522R073-F1-model_v4 Uncharacterized protein | 22.3 | 7.93E-06 | Chitinophagaceae bacterium |
| 611 | AF-A0A1M5BB02-F1-model_v4 TIGR02594 family protein | 20.8 | 2.46E-06 | Pedobacter caeni |
| 612 | AF-A0A3D6EHQ1-F1-model_v4 Uncharacterized protein | 16.3 | 1.35E-04 | Holophagaceae bacterium |
| 613 | AF-A0A427K5Y4-F1-model_v4 Glyco_hydro_19_cat domain-containing protein | 20.4 | 2.56E-05 | Erwinia sp. 198 |
| 614 | AF-A0A3M1P6I9-F1-model_v4 Glycoside hydrolase family 19 protein | 18.8 | 1.23E-05 | Bacteroidetes bacterium |
| 615 | AF-A0A848A737-F1-model_v4 Glycoside hydrolase family 19 protein | 19.6 | 8.33E-06 | Verrucomicrobia bacterium |
| 616 | AF-A0A820AHM2-F1-model_v4 Hypothetical protein | 21.4 | 2.44E-05 | Rotaria magnacalcarata |
| 617 | AF-A0A542HJN9-F1-model_v4 Putative chitinase | 21.3 | 3.43E-05 | Phycicoccus sp. SLBN-51 |
| 618 | AF-A0A3A8P171-F1-model_v4 LysM peptidoglycan-binding domain-containing protein | 21.1 | 2.96E-05 | Corallococcus llansteffanensis |
| 619 | AF-A0A1X0A4E3-F1-model_v4 Peptidase_M15_4 domain-containing protein | 18.4 | 6.17E-05 | Mycobacterium aquaticum |
| 620 | AF-U3A488-F1-model_v4 Uncharacterized protein | 21.3 | 9.57E-05 | Vibrio proteolyticus NBRC 13287 |
| 621 | AF-A0A411DQR3-F1-model_v4 Peptidase_M23 domain-containing protein | 20.8 | 9.18E-06 | Chryseobacterium indologenes |
| 622 | AF-A0A2T1LT79-F1-model_v4 Glyco_hydro_19_cat domain-containing protein | 19.7 | 7.87E-05 | Aphanothece hegewaldii CCALA 016 |
| 623 | AF-A0A845XAK1-F1-model_v4 Uncharacterized protein | 16.1 | 6.80E-05 | Spirulina sp. SIO3F2 |

| | | | | |
|---|---|---|---|---|
| 624 | AF-A0A4Q7TH75-F1-model_v4 Putative chitinase | 20.4 | 1.65E-05 | Bradyrhizobium sp. BK707 |
| 625 | AF-S3IX33-F1-model_v4 EF-hand domain-containing protein | 19.8 | 9.18E-06 | Cedecea davisae DSM 4568 |
| 626 | AF-A0A2S8ADZ0-F1-model_v4 LysM domain-containing protein | 21.9 | 5.11E-06 | Apibacter adventoris |
| 627 | AF-A0A846ADR9-F1-model_v4 Uncharacterized protein | 22.3 | 3.43E-05 | Leptolyngbya sp. SIO4C5 |
| 628 | AF-A0A1X9Z1V6-F1-model_v4 Uncharacterized protein | 21 | 8.33E-06 | Sphingobacteriaceae bacterium GW460-11-11-14-LB5 |
| 629 | AF-A0A2R7PY95-F1-model_v4 DNA primase | 29.3 | 5.59E-05 | Stenotrophomonas sp. HMWF022 |
| 630 | AF-A0A814Z504-F1-model_v4 Hypothetical protein | 21.7 | 4.17E-05 | Adineta steineri |
| 631 | AF-A0A1I5YYG4-F1-model_v4 Predicted chitinase | 25.5 | 8.74E-06 | Pseudarcicella hirudinis |
| 632 | AF-A0A143PKM9-F1-model_v4 Putative family GH19 chitinase | 19.9 | 2.69E-05 | Luteitalea pratensis |
| 633 | AF-A0A1I2X2J3-F1-model_v4 Putative chitinase | 20 | 2.01E-05 | Paracoccus aminovorans |
| 634 | AF-A0A1Y0FVK0-F1-model_v4 Uncharacterized protein | 18.4 | 5.87E-05 | Cellvibrio sp. PSBB006 |
| 635 | AF-A0A1G6UCX6-F1-model_v4 Chitinase class I | 16.2 | 9.18E-06 | Williamwhitmania taraxaci |
| 636 | AF-A0A1G6MV18-F1-model_v4 Predicted chitinase | 19.8 | 1.36E-05 | Variovorax sp. CF079 |
| 637 | AF-A0A1Q9VXV0-F1-model_v4 Peptidase_C39_2 domain-containing protein | 22.1 | 5.07E-05 | Corynebacterium sp. CNJ-954 |
| 638 | AF-A0A6C2YT43-F1-model_v4 Uncharacterized protein | 18.3 | 2.96E-05 | Tuwongella immobilis |
| 639 | AF-A0A554UDY6-F1-model_v4 Uncharacterized protein | 21.3 | 6.47E-05 | Mycobacterium sp. KBS0706 |
| 640 | AF-A0A0G2ZNT5-F1-model_v4 Membrane-bound lytic murein transglycosylase D | 20.4 | 7.14E-05 | Archangium gephyra |
| 641 | AF-A0A4R1CED4-F1-model_v4 Peptidase_M15_4 domain-containing protein | 18.7 | 9.11E-05 | Nocardioides jejuensis |
| 642 | AF-A0A2N8QLD6-F1-model_v4 Uncharacterized protein | 20.2 | 6.17E-05 | Paraburkholderia fungorum |
| 643 | AF-A0A2U1JXG8-F1-model_v4 Glycoside hydrolase | 24.5 | 1.01E-04 | Flavobacterium laiguense |
| 644 | AF-A0A6N6ZW30-F1-model_v4 Putative chitinase | 21.2 | 9.11E-05 | Rhodospirillaceae bacterium |
| 645 | AF-A0A814PAW5-F1-model_v4 Hypothetical protein | 19.9 | 7.87E-05 | Rotaria sordida |
| 646 | AF-A6UAM1-F1-model_v4 Chitinase-like protein | 21.8 | 1.01E-05 | Sinorhizobium medicae WSM419 |
| 647 | AF-A0A0Q6D2X3-F1-model_v4 Uncharacterized protein | 24.4 | 1.36E-05 | Methylobacterium sp. Leaf456 |
| 648 | AF-A0A7K0DEU4-F1-model_v4 Glyco_hydro_19_cat domain-containing protein | 20.7 | 3.11E-05 | Nocardia macrotermitis |
| 649 | AF-A0A3R9YZH7-F1-model_v4 Glyco_hydro_19_cat domain-containing protein | 21.3 | 1.73E-05 | Variovorax sp. DXTD-1 |
| 650 | AF-A0A2E8PLW7-F1-model_v4 Uncharacterized protein | 19.9 | 8.26E-05 | Rhodobacteraceae bacterium |
| 651 | AF-A0A1M5G7P6-F1-model_v4 Predicted chitinase | 21.7 | 5.92E-06 | Chryseobacterium vrystaatense |
| 652 | AF-A0A0A0M6Q0-F1-model_v4 Chitinase | 21 | 3.43E-05 | Lysobacter defluvii IMMIB APB-9 = DSM 18482 |
| 653 | AF-A0A0A8NWI3-F1-model_v4 Uncharacterized protein | 22.9 | 5.07E-05 | Xenorhabdus nematophila str. Websteri |
| 654 | AF-A0A2S8ZFW8-F1-model_v4 Peptidase_M23 domain-containing protein | 19.8 | 6.52E-06 | Chryseobacterium sp. MYb7 |

| | | | | |
|---|---|---|---|---|
| 655 | AF-A0A7K0DVV4-F1-model_v4 Glyco_hydro_19_cat domain-containing protein | 22.4 | 4.83E-05 | Nocardia aurantia |
| 656 | AF-A0A107FN43-F1-model_v4 Uncharacterized protein | 23.1 | 1.28E-04 | Burkholderia ubonensis |
| 657 | AF-G1VIN7-F1-model_v4 Uncharacterized protein | 22.4 | 3.27E-05 | Prevotella sp. C561 |
| 658 | AF-A0A7Y0H8X4-F1-model_v4 Uncharacterized protein | 22 | 7.14E-05 | Glaciimonas sp. |
| 659 | AF-A0A0W0YRP2-F1-model_v4 Uncharacterized protein | 18.6 | 5.92E-06 | Legionella shakespearei DSM 23087 |
| 660 | AF-A0A5C7W4X4-F1-model_v4 Glyco_hydro_19_cat domain-containing protein | 19.9 | 4.17E-05 | Ottowia sp. |
| 661 | AF-A0A1H7QTQ6-F1-model_v4 Putative chitinase | 20.1 | 1.35E-04 | Rhodococcus maanshanensis |
| 662 | AF-A0A7U5YL47-F1-model_v4 Uncharacterized protein | 24.4 | 4.38E-05 | Salmonella enterica subsp. diarizonae serovar 48:i:z |
| 663 | AF-E5XQC8-F1-model_v4 Uncharacterized protein | 20.5 | 1.23E-05 | Segniliparus rugosus ATCC BAA-974 |
| 664 | AF-A0A6A5BXB1-F1-model_v4 SH3b domain-containing protein | 19 | 2.01E-05 | Naegleria fowleri |
| 665 | AF-A0A2K8W0N2-F1-model_v4 Uncharacterized protein | 23.2 | 2.96E-05 | Dickeya solani RNS 08.23.3.1.A |
| 666 | AF-A0A1L3ZTC9-F1-model_v4 Glyco_hydro_19_cat domain-containing protein | 22.9 | 1.72E-04 | Tardibacter chloracetimidivorans |
| 667 | AF-A0A1G5UCC5-F1-model_v4 Putative chitinase | 27.1 | 4.60E-05 | Sphingomonas sp. NFR15 |
| 668 | AF-A0A3G6R136-F1-model_v4 Peptidase_M23 domain-containing protein | 29.2 | 2.82E-05 | Chryseobacterium shandongense |
| 669 | AF-A0A291RWC6-F1-model_v4 Glyco_hydro_19_cat domain-containing protein | 23.1 | 7.87E-05 | Nocardia terpenica |
| 670 | AF-A0A3D9URJ3-F1-model_v4 Putative chitinase | 23.4 | 2.56E-05 | Xenorhabdus cabanillasii |
| 671 | AF-A0A4R1NF17-F1-model_v4 Putative chitinase | 22.5 | 6.80E-05 | Sodalis ligni |
| 672 | AF-A0A5C1B466-F1-model_v4 Glycoside hydrolase | 19.6 | 2.30E-04 | Bosea sp. F3-2 |
| 673 | AF-A1B8K2-F1-model_v4 Peptidoglycan-binding domain 1 protein | 19.1 | 1.49E-04 | Paracoccus denitrificans PD1222 |
| 674 | AF-A0A110AZC3-F1-model_v4 Putative chitinase | 18.1 | 5.07E-05 | Mucilaginibacter gotjawali |
| 675 | AF-E2CHK7-F1-model_v4 Gene 10 protein | 21.3 | 1.01E-04 | Roseibium sp. TrichSKD4 |
| 676 | AF-A0A2G2ATW9-F1-model_v4 EF-hand domain-containing protein | 14.9 | 5.87E-05 | Sulfurimonas sp. |
| 677 | AF-A0A6L5FPK7-F1-model_v4 Uncharacterized protein | 20.3 | 2.32E-05 | Hyphomicrobiales bacterium |
| 678 | AF-A0A2N5AE67-F1-model_v4 Peptidase C1 | 20.5 | 5.87E-05 | Klebsiella variicola |
| 679 | AF-A0A420CG58-F1-model_v4 Uncharacterized protein (TIGR02594 family) | 17.7 | 1.29E-05 | Chryseobacterium sp. AG363 |
| 680 | AF-A0A6C2E0Z7-F1-model_v4 Uncharacterized protein | 20.5 | 5.33E-05 | Pseudanabaena sp. UWO310 |
| 681 | AF-A0A0A4ADK1-F1-model_v4 Glycoside hydrolase | 20.2 | 4.83E-05 | Erwinia typographi |
| 682 | AF-A0A1G6M265-F1-model_v4 Putative chitinase | 20 | 6.17E-05 | Sphingomonas sp. YR710 |
| 683 | AF-A0A7D6E0C4-F1-model_v4 M15 family metallopeptidase | 18.2 | 1.28E-04 | Mycobacterium gordonae |
| 684 | AF-A0A659LE29-F1-model_v4 Uncharacterized protein | 26.9 | 5.87E-05 | Escherichia sp. E2586 |

| | | | | |
|---|---|---|---|---|
| 685 | AF-A0A3A1WDF6-F1-model_v4 Glycoside hydrolase family 19 protein | 21.6 | 1.17E-05 | Acidovorax cavernicola |
| 686 | AF-A0A6I4WUY6-F1-model_v4 Glyco_hydro_19_cat domain-containing protein | 21.8 | 9.57E-05 | Rhodococcus rhodochrous |
| 687 | AF-A0A813NGZ8-F1-model_v4 Hypothetical protein | 19.1 | 3.60E-05 | Adineta ricciae |
| 688 | AF-A0A6G9AIZ2-F1-model_v4 Uncharacterized protein | 16 | 2.56E-05 | Spirosoma aureum |
| 689 | AF-A0A2W6F323-F1-model_v4 Uncharacterized protein | 21 | 7.87E-05 | Pseudonocardiales bacterium |
| 690 | AF-A0A5F1UIG0-F1-model_v4 Chitinase | 19.2 | 9.11E-05 | Escherichia coli |
| 691 | AF-A0A162Y446-F1-model_v4 Uncharacterized protein | 21.4 | 2.30E-04 | Aquimarina aggregata |
| 692 | AF-A0A3G8M9H7-F1-model_v4 Uncharacterized protein | 21.8 | 1.90E-04 | Methylocystis rosea |
| 693 | AF-A0A246GEZ6-F1-model_v4 Uncharacterized protein | 21.2 | 2.11E-05 | Flavobacterium columnare |
| 694 | AF-A0A6C2D4A5-F1-model_v4 Glycoside hydrolase family 19 protein | 23.8 | 8.26E-05 | Zoogloea oleivorans |
| 695 | AF-A0A4P6H3I9-F1-model_v4 Uncharacterized protein | 19.3 | 3.27E-05 | Mesorhizobium sp. Pch-S |
| 696 | AF-A0A0K1E680-F1-model_v4 Uncharacterized protein | 19.8 | 1.22E-04 | Chondromyces crocatus |
| 697 | AF-A0A1S7TA89-F1-model_v4 Peptidoglycan-binding domain-containing protein | 23.2 | 7.49E-05 | Agrobacterium fabacearum CFBP 5771 |
| 698 | AF-A0A848L6Y9-F1-model_v4 LysM peptidoglycan-binding domain-containing protein | 21.1 | 2.96E-05 | Pyxidicoccus fallax |
| 699 | AF-A0A1Y2SJI6-F1-model_v4 Triacylglycerol lipase | 21.7 | 1.01E-04 | Xenorhabdus vietnamensis |
| 700 | AF-A0A009F2I3-F1-model_v4 Chitinase class I family protein | 19.9 | 2.09E-04 | Acinetobacter baumannii 348935 |
| 701 | AF-A0A0N1BGC7-F1-model_v4 Glyco_hydro_19_cat domain-containing protein | 19.2 | 4.60E-05 | Blastomonas sp. AAP25 |
| 702 | AF-A0A0D5A628-F1-model_v4 Phage endolysin | 18.3 | 1.49E-04 | Rhodococcus sp. B7740 |
| 703 | AF-A0A813UZ72-F1-model_v4 Hypothetical protein | 19.7 | 1.35E-04 | Adineta ricciae |
| 704 | AF-A0A1I3IQ90-F1-model_v4 Uncharacterized protein | 20.5 | 2.32E-05 | Halpernia frigidisoli |
| 705 | AF-A0A3N5LHL6-F1-model_v4 Chitinase | 19 | 2.56E-05 | Ignavibacteriae bacterium |
| 706 | AF-A1B829-F1-model_v4 Peptidoglycan-binding domain 1 protein | 19.1 | 1.28E-04 | Paracoccus denitrificans PD1222 |
| 707 | AF-A0A814F7X5-F1-model_v4 Hypothetical protein | 20 | 6.17E-05 | Adineta ricciae |
| 708 | AF-A0A816Y303-F1-model_v4 Hypothetical protein | 21.4 | 1.06E-04 | Rotaria magnacalcarata |
| 709 | AF-A0A2A2E430-F1-model_v4 Uncharacterized protein | 19.9 | 2.19E-04 | Pseudomonas sp. PIC25 |
| 710 | AF-A0A2N5JRY8-F1-model_v4 Chitinase | 18.5 | 1.81E-04 | Cyanobacteria bacterium M5B4 |
| 711 | AF-A0A820A9W3-F1-model_v4 Hypothetical protein | 21.2 | 1.35E-04 | Rotaria magnacalcarata |
| 712 | AF-A0A2N3HRE8-F1-model_v4 Uncharacterized protein | 17.3 | 2.19E-04 | Labilibaculum manganireducens |
| 713 | AF-A0A1G8IEF5-F1-model_v4 Predicted chitinase | 17 | 1.56E-04 | Pseudomonas panipatensis |
| 714 | AF-A0A4Q2ZPD2-F1-model_v4 DUF4280 domain-containing protein | 19.3 | 1.57E-05 | Sphingobacteriales bacterium |

| | | | | |
|---|---|---|---|---|
| 715 | AF-N6VJK2-F1-model_v4 Uncharacterized protein | 24.7 | 9.57E-05 | Bartonella vinsonii subsp. berkhoffii str. Tweed |
| 716 | AF-A0A7Y7NM63-F1-model_v4 Glycoside hydrolase | 19.7 | 1.99E-04 | Geobacteraceae bacterium |
| 717 | AF-A0A4U8SCF5-F1-model_v4 Uncharacterized protein | 20.8 | 5.87E-05 | Helicobacter sp. MIT 05-5294 |
| 718 | AF-A0A4P6FHZ6-F1-model_v4 Glycoside hydrolase | 20.1 | 3.27E-05 | Sphingosinicella sp. BN140058 |
| 719 | AF-A0A547Q6C0-F1-model_v4 Glycoside hydrolase family 19 protein | 23.6 | 5.59E-05 | Paenimaribius caenipelagi |
| 720 | AF-A0A009KM83-F1-model_v4 Chitinase class I family protein | 19.1 | 1.57E-05 | Acinetobacter baumannii 146457 |
| 721 | AF-A0A7S9NFX5-F1-model_v4 Uncharacterized protein | 19.7 | 1.29E-05 | Campylobacter concisus |
| 722 | AF-A0A840WWV7-F1-model_v4 Putative chitinase | 19.8 | 3.43E-05 | Sphingomonas sp. R3G7C |
| 723 | AF-A0A285XCQ1-F1-model_v4 Putative chitinase | 20.3 | 1.56E-04 | Ensifer adhaerens |
| 724 | AF-F3B087-F1-model_v4 Uncharacterized protein | 26.4 | 2.32E-05 | Lachnospiraceae oral taxon 107 str. F0167 |
| 725 | AF-A0A3N0UDN0-F1-model_v4 Uncharacterized protein | 24 | 1.99E-04 | Lonsdalea populi |
| 726 | AF-E8V8T8-F1-model_v4 Glycoside hydrolase family 19 | 19.4 | 3.60E-05 | Terriglobus saanensis SP1PR4 |
| 727 | AF-A0A7W6H9C9-F1-model_v4 Putative chitinase | 23 | 1.64E-04 | Aurantimonas endophytica |
| 728 | AF-A0A2U3N492-F1-model_v4 Chitinase class I | 21.9 | 1.22E-04 | Acinetobacter stercoris |
| 729 | AF-B0VVF0-F1-model_v4 Glyco_hydro_19_cat domain-containing protein | 22.2 | 2.42E-04 | Acinetobacter baumannii SDF |
| 730 | AF-A0A7G1KTC2-F1-model_v4 Glyco_hydro_19_cat domain-containing protein | 21.9 | 5.33E-05 | Nocardia wallacei |
| 731 | AF-A0A814RWY9-F1-model_v4 Hypothetical protein | 20.2 | 1.28E-04 | Adineta ricciae |
| 732 | AF-T5LSC8-F1-model_v4 Uncharacterized protein | 19.1 | 6.17E-05 | Helicobacter bilis ATCC 43879 |
| 733 | AF-I4VWR8-F1-model_v4 Peptidoglycan binding domain/papain family cysteine protease | 20.4 | 1.16E-04 | Rhodanobacter spathiphylli B39 |
| 734 | AF-A0A4Q9XNV5-F1-model_v4 PG_binding_1 domain-containing protein | 12.8 | 4.14E-04 | Bowmanella sp. JS7-9 |
| 735 | AF-A0A8B3IP40-F1-model_v4 Peptidoglycan-binding protein | 18.5 | 1.56E-04 | Brucella anthropi |
| 736 | AF-A0A7T8U7Y3-F1-model_v4 Uncharacterized protein | 21.9 | 5.33E-05 | Chryseobacterium indologenes |
| 737 | AF-A0A172XU88-F1-model_v4 Uncharacterized protein | 18.1 | 3.58E-04 | Chryseobacterium glaciei |
| 738 | AF-A0A828S2D7-F1-model_v4 Chitinase class I family protein | 19.8 | 1.49E-04 | Escherichia coli STEC_7v |
| 739 | AF-A0A418N584-F1-model_v4 Fibronectin type-III domain-containing protein | 19.1 | 3.27E-05 | Muricauda aequoris |
| 740 | AF-A0A5S9SAJ0-F1-model_v4 Uncharacterized protein | 19.3 | 1.81E-04 | Tenacibaculum maritimum |
| 741 | AF-F7XBP7-F1-model_v4 Chitinase-like protein | 21.1 | 1.06E-04 | Sinorhizobium meliloti SM11 |
| 742 | AF-A0A5C7YJZ2-F1-model_v4 Uncharacterized protein | 18.4 | 1.72E-04 | Mycolicibacterium mageritense |
| 743 | AF-A0A815PJ33-F1-model_v4 Hypothetical protein | 19.9 | 7.49E-05 | Adineta steineri |
| 744 | AF-A0A0N1FH71-F1-model_v4 Glyco_hydro_19_cat domain-containing protein | 17.5 | 1.01E-04 | Bosea vaviloviae |

| | | | | |
|---|---|---|---|---|
| 745 | AF-J4K9Y2-F1-model_v4 Cell wall-binding repeat protein | 23.7 | 2.32E-05 | Lachnoanaerobaculum sp. ICM7 |
| 746 | AF-A0A816QIX4-F1-model_v4 Hypothetical protein | 21.4 | 1.90E-04 | Rotaria magnacalcarata |
| 747 | AF-A0A1U9KQE6-F1-model_v4 Lysozyme | 21.6 | 1.73E-05 | Neoasaia chiangmaiensis |
| 748 | AF-A0A1B9L3G3-F1-model_v4 Glyco_hydro_19_cat domain-containing protein | 22.4 | 1.11E-04 | Gilliamella apicola |
| 749 | AF-A0A818JJA9-F1-model_v4 Hypothetical protein | 21.8 | 1.99E-04 | Rotaria socialis |
| 750 | AF-A0A7Y7V7U8-F1-model_v4 Uncharacterized protein | 17.5 | 7.87E-05 | Pseudomonas edaphica |
| 751 | AF-A0A2Z5V6X8-F1-model_v4 Chitinase class I family protein | 21.6 | 1.22E-04 | Candidatus Rickettsiella viridis |
| 752 | AF-A0A244E2Q8-F1-model_v4 Uncharacterized protein | 19.5 | 3.58E-04 | Variovorax sp. JS1663 |
| 753 | AF-A0A5M8PAE1-F1-model_v4 Peptidoglycan-binding protein | 23.4 | 2.30E-04 | Candidatus Tokpelaia sp. |
| 754 | AF-A0A135P869-F1-model_v4 Uncharacterized protein | 20.4 | 6.47E-05 | Agrobacterium bohemicum |
| 755 | AF-A0A3M1C0M8-F1-model_v4 Glyco_hydro_19_cat domain-containing protein | 21.4 | 4.38E-05 | Bacteroidetes bacterium |
| 756 | AF-A0A3N7ZY30-F1-model_v4 Uncharacterized protein | 18.5 | 3.24E-04 | Burkholderia sp. Bp9131 |
| 757 | AF-A8YDG4-F1-model_v4 Genome sequencing data, contig C292 | 22.6 | 5.07E-05 | Microcystis aeruginosa PCC 7806 |
| 758 | AF-A0A258ZJH5-F1-model_v4 Uncharacterized protein | 18.4 | 5.33E-05 | Gallionellales bacterium 24-53-125 |
| 759 | AF-A0A7G2TII5-F1-model_v4 Peptidoglycan-binding protein | 19.7 | 2.19E-04 | Salipiger sp. |
| 760 | AF-A0A4U1HDF6-F1-model_v4 Glyco_hydro_19_cat domain-containing protein | 21.4 | 7.87E-05 | Trinickia sp. 7GSK02 |
| 761 | AF-A0A7W6E9D7-F1-model_v4 Putative chitinase | 21.3 | 1.01E-04 | Aureimonas pseudogalii |
| 762 | AF-A0A1Q4SLJ9-F1-model_v4 Peptidase_C39_2 domain-containing protein | 18.3 | 2.30E-04 | Mycobacterium sp. SWH-M5 |
| 763 | AF-A0A432K1F2-F1-model_v4 Uncharacterized protein | 22.4 | 3.43E-05 | Bacteroidetes bacterium |
| 764 | AF-A0A2G6IIA3-F1-model_v4 Peptidoglycan-binding protein | 23.5 | 4.79E-04 | Rhodobacterales bacterium |
| 765 | AF-A0A7W0G196-F1-model_v4 Uncharacterized protein | 17.7 | 1.81E-04 | Acidobacteria bacterium |
| 766 | AF-A0A7Y6AY97-F1-model_v4 Uncharacterized protein | 16.7 | 1.16E-04 | Pseudomonas sp. C2B4 |
| 767 | AF-A0A2K8UQ98-F1-model_v4 Glycoside hydrolase | 20.6 | 5.55E-04 | Acinetobacter lwoffii |
| 768 | AF-A0A5P2QRQ8-F1-model_v4 Peptidoglycan-binding protein | 19.5 | 6.47E-05 | Paracoccus yeei |
| 769 | AF-B8ITL4-F1-model_v4 Uncharacterized protein | 18.6 | 1.01E-04 | Methylobacterium nodulans ORS 2060 |
| 770 | AF-A0A133ZF30-F1-model_v4 Cell wall-binding repeat protein | 20 | 7.87E-05 | Lachnoanaerobaculum saburreum |
| 771 | AF-A0A553DTW1-F1-model_v4 Uncharacterized protein | 20.8 | 2.30E-04 | Flavobacterium sp. ZT3R18 |
| 772 | AF-A0A418WDB1-F1-model_v4 Uncharacterized protein | 17.3 | 4.56E-04 | Oleomonas cavernae |
| 773 | AF-A0A8A9DHM3-F1-model_v4 C39 family peptidase | 20.1 | 1.06E-04 | Mycobacterium tuberculosis |
| 774 | AF-A0A1A9S3Z7-F1-model_v4 Glyco_hydro_19_cat domain-containing protein | 24.3 | 2.30E-04 | Eikenella sp. NML03-A-027 |
| 775 | AF-A5FHN3-F1-model_v4 Zoocin A peptidase family M23B | 18.5 | 4.60E-05 | Flavobacterium johnsoniae UW101 |

| | | | | |
|---|---|---|---|---|
| 776 | AF-A0A379DT67-F1-model_v4 Predicted chitinase | 14 | 7.14E-05 | Pragia fontium |
| 777 | AF-A0A365VPU4-F1-model_v4 Chitinase | 27.3 | 8.26E-05 | Pseudomonas sp. MWU12-2534b |
| 778 | AF-A0A538GR89-F1-model_v4 Uncharacterized protein | 17.9 | 1.81E-04 | Actinomycetia bacterium |
| 779 | AF-A0A816QYB0-F1-model_v4 Hypothetical protein | 15.3 | 9.11E-05 | Rotaria magnacalcarata |
| 780 | AF-A0A143PL75-F1-model_v4 Putative family GH19 chitinase | 20.7 | 2.19E-04 | Luteitalea pratensis |
| 781 | AF-A0A1G7UVY5-F1-model_v4 Predicted chitinase | 21.3 | 1.16E-04 | Paraburkholderia phenazinium |
| 782 | AF-A0A2T4ZIV9-F1-model_v4 Putative chitinase | 20.6 | 2.94E-04 | Phreatobacter oligotrophus |
| 783 | AF-A0A3D8I9W3-F1-model_v4 Uncharacterized protein | 27.3 | 7.87E-05 | Helicobacter didelphidarum |
| 784 | AF-A0A838L4L3-F1-model_v4 Glycoside hydrolase family 19 protein | 25.1 | 9.11E-05 | Sphingomonas chungangi |
| 785 | AF-A0A2D5FT31-F1-model_v4 Peptidoglycan-binding protein | 21.7 | 1.28E-04 | Oceanicaulis sp. |
| 786 | AF-A0A7U1GRE8-F1-model_v4 Uncharacterized protein | 22.7 | 2.80E-04 | Heyndrickxia vini |
| 787 | AF-A0A250KK30-F1-model_v4 Uncharacterized protein | 17.5 | 7.87E-05 | Prevotella melaninogenica |
| 788 | AF-A0A5J5LX70-F1-model_v4 Peptidase_M15_3 domain-containing protein | 21.9 | 1.90E-04 | Microcystis aeruginosa EAWAG127a |
| 789 | AF-A0A0Q0CEV6-F1-model_v4 EF hand domain protein | 25.1 | 1.81E-04 | Pseudomonas syringae pv. tomato |
| 790 | AF-A0A2V3J417-F1-model_v4 Acidic endochitinase SP2 | 18 | 9.57E-05 | Gracilariopsis chorda |
| 791 | AF-A0A7X5QHQ8-F1-model_v4 Glyco_hydro_19_cat domain-containing protein | 40.5 | 3.75E-04 | Photorhabdus cinerea |
| 792 | AF-A0A1T1ANS5-F1-model_v4 Uncharacterized protein | 22 | 2.42E-04 | Rhodoferax fermentans |
| 793 | AF-A0A814KB02-F1-model_v4 Hypothetical protein | 17 | 5.33E-05 | Adineta steineri |
| 794 | AF-A0A849VNP2-F1-model_v4 Glycoside hydrolase family 19 protein | 22.3 | 2.67E-04 | Phyllobacterium pellucidum |
| 795 | AF-A0A1A6BJJ9-F1-model_v4 Peptidase_M15_4 domain-containing protein | 18.8 | 6.75E-04 | Mycobacterium gordonae |
| 796 | AF-A0A348HIL9-F1-model_v4 Glycoside hydrolase, family 19 | 17.6 | 2.09E-04 | Zymobacter palmae |
| 797 | AF-A0A2T4M6D7-F1-model_v4 Uncharacterized protein | 22.4 | 1.64E-04 | Staphylococcus auricularis |
| 798 | AF-A0A814Y1S4-F1-model_v4 Hypothetical protein | 21.4 | 1.64E-04 | Rotaria sp. Silwood1 |
| 799 | AF-A0A539DJX9-F1-model_v4 Putative chitinase | 17.7 | 3.94E-04 | Alphaproteobacteria bacterium |
| 800 | AF-A0A2G1B1B8-F1-model_v4 Uncharacterized protein | 19 | 2.09E-04 | Vibrio splendidus |
| 801 | AF-A0A5E4W1D3-F1-model_v4 Glyco_hydro_19_cat domain-containing protein | 18.3 | 3.24E-04 | Pandoraea cepalis |
| 802 | AF-A0A6P1VQN1-F1-model_v4 Glyco_hydro_19_cat domain-containing protein | 16.1 | 2.80E-04 | Spirosoma endbachense |
| 803 | AF-A6UIU2-F1-model_v4 Chitinase-like protein | 20.9 | 1.28E-04 | Sinorhizobium medicae WSM419 |
| 804 | AF-A0A2N3HTT2-F1-model_v4 Uncharacterized protein | 18.7 | 3.75E-04 | Labilibaculum filiforme |
| 805 | AF-A7ILJ3-F1-model_v4 Glycoside hydrolase, family 19 | 20.1 | 3.58E-04 | Xanthobacter autotrophicus Py2 |
| 806 | AF-I0R4S1-F1-model_v4 Chitinase class I domain protein | 20.9 | 6.47E-05 | Lachnoanaerobaculum saburreum F0468 |

| | | | | |
|---|---|---|---|---|
| 807 | AF-A0A259Y5G5-F1-model_v4 Uncharacterized protein | 17.5 | 2.54E-04 | Rhodococcus sp. 06-621-2 |
| 808 | AF-A0A542GBM4-F1-model_v4 Putative chitinase | 22.5 | 2.67E-04 | Microbacterium sp. SLBN-1 |
| 809 | AF-A0A2E9AMK4-F1-model_v4 EF-hand domain-containing protein | 21.3 | 1.90E-04 | Salinisphaera sp. |
| 810 | AF-A0A2A2GQM1-F1-model_v4 Uncharacterized protein | 24.8 | 3.41E-04 | Helicobacter sp. TUL |
| 811 | AF-D6ZB17-F1-model_v4 Chitinase-like protein | 18.2 | 2.09E-04 | Segniliparus rotundus DSM 44985 |
| 812 | AF-A0A1H2S9B5-F1-model_v4 Predicted chitinase | 19 | 2.19E-04 | Albimonas donghaensis |
| 813 | AF-A0A815JS44-F1-model_v4 Hypothetical protein | 15.5 | 1.64E-04 | Rotaria magnacalcarata |
| 814 | AF-N9H5U8-F1-model_v4 Glyco_hydro_19_cat domain-containing protein | 19.1 | 1.71E-03 | Acinetobacter lwoffii ATCC 9957 = CIP 70.31 |
| 815 | AF-A0A2T4TJS8-F1-model_v4 Uncharacterized protein | 15.6 | 2.09E-04 | Prevotella sp. oral taxon 313 |
| 816 | AF-A0A4R2SPY3-F1-model_v4 Putative chitinase | 19.2 | 1.22E-04 | Cricetibacter osteomyelitidis |
| 817 | AF-A0A191YV54-F1-model_v4 Uncharacterized protein | 20.2 | 4.35E-04 | Pseudomonas silesiensis |
| 818 | AF-A0A3D8VEL7-F1-model_v4 Uncharacterized protein | 21.3 | 3.24E-04 | Lysobacter soli |
| 819 | AF-A0A7U0YBD2-F1-model_v4 Uncharacterized protein | 42.2 | 4.79E-04 | Klebsiella quasipneumoniae |
| 820 | AF-A0A1W6E0D5-F1-model_v4 Glyco_hydro_19_cat domain-containing protein | 18.7 | 4.56E-04 | Fibrella sp. ES10-3-2-2 |
| 821 | AF-A0A1B9M7W4-F1-model_v4 Glyco_hydro_19_cat domain-containing protein | 22.4 | 5.28E-04 | Gilliamella apicola |
| 822 | AF-A0A0N0V0H1-F1-model_v4 Peptidoglycan-binding protein | 14.1 | 6.75E-04 | Candidatus Magnetomorum sp. HK-1 |
| 823 | AF-A0A817EZ17-F1-model_v4 Hypothetical protein | 16.1 | 2.80E-04 | Rotaria sp. Silwood2 |
| 824 | AF-A0A7G7ETT1-F1-model_v4 Glyco_hydro_19_cat domain-containing protein | 19 | 2.30E-04 | Aeromonas jandaei |
| 825 | AF-A0A0D6AWQ0-F1-model_v4 Phage endolysin | 19.1 | 2.67E-04 | Geminocystis sp. NIES-3709 |
| 826 | AF-A0A1B0ZP20-F1-model_v4 Peptidoglycan-binding protein | 18.7 | 6.75E-04 | Phaeobacter gallaeciensis |
| 827 | AF-A0A819NU12-F1-model_v4 Hypothetical protein | 15.7 | 2.94E-04 | Rotaria sp. Silwood2 |
| 828 | AF-A0A2U9U0E8-F1-model_v4 Uncharacterized protein | 18.6 | 5.28E-04 | Methylobacterium sp. XJLW |
| 829 | AF-A0A8B2BSX0-F1-model_v4 Uncharacterized protein | 19.5 | 2.19E-04 | Prevotella melaninogenica |
| 830 | AF-A0A1G4VFR2-F1-model_v4 Predicted chitinase | 19 | 4.14E-04 | Mycolicibacterium fluoranthenivorans |
| 831 | AF-A3IWZ0-F1-model_v4 Lysozyme, putative | 25.1 | 1.01E-04 | Crocosphaera chwakensis CCY0110 |
| 832 | AF-A0A250KLN4-F1-model_v4 Uncharacterized protein | 18.3 | 2.54E-04 | Prevotella melaninogenica |
| 833 | AF-A0A1E4Q6W7-F1-model_v4 Glyco_hydro_19_cat domain-containing protein | 18.9 | 3.58E-04 | Rhodanobacter sp. SCN 68-63 |
| 834 | AF-A0A3N0WNJ5-F1-model_v4 Glyco_hydro_19_cat domain-containing protein | 19.4 | 9.57E-05 | Chryseobacterium sp. G0240 |
| 835 | AF-A0A2C9AR95-F1-model_v4 Predicted chitinase | 18.5 | 9.97E-04 | Burkholderia sp. YR290 |
| 836 | AF-A0A7C4VE81-F1-model_v4 Uncharacterized protein | 25.3 | 2.09E-04 | Proteobacteria bacterium |
| 837 | AF-A0A0P9G2T3-F1-model_v4 Endopeptidase | 21 | 1.10E-03 | Paenibacillus sp. A3 |

| | | | | |
|---|---|---|---|---|
| 838 | AF-A0A814HII2-F1-model_v4 Hypothetical protein | 18.1 | 3.41E-04 | Adineta ricciae |
| 839 | AF-A0A0P7ENS9-F1-model_v4 Uncharacterized protein | 19.7 | 4.35E-04 | Vibrio alginolyticus |
| 840 | AF-A0A1S8LDW0-F1-model_v4 N-acetylmuramoyl-L-alanine amidase CwlA | 18.6 | 3.58E-04 | Clostridium felsineum |
| 841 | AF-A0A2Z6UTU2-F1-model_v4 Uncharacterized protein | 23.2 | 1.35E-04 | Microcystis aeruginosa Sj |
| 842 | AF-A0A0H4BEX4-F1-model_v4 Uncharacterized protein | 18.5 | 5.03E-04 | Synechococcus sp. WH 8020 |
| 843 | AF-A0A3A1WI00-F1-model_v4 Peptidoglycan-binding protein | 17.4 | 1.71E-03 | Aureimonas flava |
| 844 | AF-A0A1V3JEL9-F1-model_v4 Uncharacterized protein | 20 | 1.63E-03 | Rodentibacter genomosp. 2 |
| 845 | AF-A0A484G9X4-F1-model_v4 Glyco_hydro_19_cat domain-containing protein | 22.8 | 1.05E-03 | Candidatus Schmidhempelia bombi str. Bimp |
| 846 | AF-A0A814KEG6-F1-model_v4 Hypothetical protein | 14.9 | 1.56E-04 | Rotaria sordida |
| 847 | AF-H1ZEB6-F1-model_v4 Uncharacterized protein | 18.5 | 1.11E-04 | Myroides odoratus DSM 2801 |
| 848 | AF-A0A858WV28-F1-model_v4 Uncharacterized protein | 18.2 | 6.42E-04 | Starkeya sp. ORNL1 |
| 849 | AF-D6Z8B3-F1-model_v4 Glycoside hydrolase family 19 | 19.1 | 3.41E-04 | Segniliparus rotundus DSM 44985 |
| 850 | AF-A0A653KIA7-F1-model_v4 EF-hand domain-containing protein | 17.3 | 9.49E-04 | Aeromonas salmonicida |
| 851 | AF-B6VN61-F1-model_v4 Chitinase class i family protein | 37.3 | 3.91E-03 | Photorhabdus asymbiotica subsp. asymbiotica ATCC 43949 |
| 852 | AF-A0A7T8YXX9-F1-model_v4 Uncharacterized protein | 29.4 | 5.03E-04 | Klebsiella pneumoniae |
| 853 | AF-A0A158BUG6-F1-model_v4 EF hand domain-containing protein | 21.7 | 8.20E-04 | Caballeronia catudaia |
| 854 | AF-A0A2U0WHH3-F1-model_v4 Putative chitinase | 20.4 | 1.28E-04 | Paraburkholderia sp. OV555 |
| 855 | AF-A0A1X0D8M1-F1-model_v4 Peptidase_C39_2 domain-containing protein | 17 | 1.27E-03 | Mycolicibacterium insubricum |
| 856 | AF-A0A0P9PSX5-F1-model_v4 Uncharacterized protein | 18.5 | 9.04E-04 | Pseudomonas syringae pv. delphinii |
| 857 | AF-A0A519KV85-F1-model_v4 Glycoside hydrolase family 19 protein | 16.5 | 3.58E-04 | Brevundimonas sp. |
| 858 | AF-A0A327NRP3-F1-model_v4 Uncharacterized protein | 19.8 | 1.65E-05 | Spirosoma telluris |
| 859 | AF-A0A419TRX3-F1-model_v4 Putative chitinase | 16.2 | 9.49E-04 | Collimonas fungivorans |
| 860 | AF-A0A7W8LHA3-F1-model_v4 Putative chitinase | 20.9 | 9.04E-04 | Paraburkholderia sp. HC6.4b |
| 861 | AF-A0A6P2ZV10-F1-model_v4 Calcium-binding protein | 22.4 | 8.20E-04 | Burkholderia contaminans |
| 862 | AF-A0A844WU65-F1-model_v4 Uncharacterized protein | 23.5 | 1.71E-03 | Gilliamella sp. Pas-s27 |
| 863 | AF-A0A8B2C520-F1-model_v4 Peptidoglycan DD-metalloendopeptidase family protein | 16.7 | 5.28E-04 | Prevotella jejuni |
| 864 | AF-A0A814ZZB6-F1-model_v4 Hypothetical protein | 15.7 | 2.54E-04 | Rotaria sp. Silwood1 |
| 865 | AF-A0A3D3MGG6-F1-model_v4 Uncharacterized protein | 19 | 1.27E-03 | Chryseobacterium sp. |
| 866 | AF-A0A6J4QT41-F1-model_v4 GH19 | 19.5 | 1.88E-03 | uncultured Rubrobacteraceae bacterium |
| 867 | AF-A0A2E3FA46-F1-model_v4 Peptidoglycan-binding protein | 32 | 1.34E-03 | Rhodobacteraceae bacterium |

| | | | | |
|---|---|---|---|---|
| 868 | AF-A0A7D7VSC2-F1-model_v4 Uncharacterized protein | 26.9 | 1.10E-03 | Flavobacteriaceae bacterium |
| 869 | AF-A0A7R7ZD40-F1-model_v4 EF-hand domain-containing protein | 19.5 | 1.71E-03 | Aeromonas caviae |
| 870 | AF-A0A1V3JFM4-F1-model_v4 Uncharacterized protein | 35.3 | 1.40E-03 | Rodentibacter myodis |
| 871 | AF-A0A7Y3X6G8-F1-model_v4 Chitinase | 21.5 | 3.24E-04 | Flavobacterium sp. CLA17 |
| 872 | AF-A0A815ALY3-F1-model_v4 Hypothetical protein | 16.5 | 4.35E-04 | Rotaria sp. Silwood1 |
| 873 | AF-A0A3M7QQS0-F1-model_v4 Chitinase | 18.7 | 2.40E-03 | Brachionus plicatilis |
| 874 | AF-A0A2D9KQK3-F1-model_v4 Uncharacterized protein | 19.6 | 1.98E-03 | Rickettsiales bacterium |
| 875 | AF-A0A806XLZ8-F1-model_v4 Uncharacterized protein | 19.7 | 7.08E-04 | Aeromonas hydrophila |
| 876 | AF-A0A7Z0L8T1-F1-model_v4 LysM peptidoglycan-binding domain-containing protein | 23.8 | 9.04E-04 | Gemella palaticanis |
| 877 | AF-A0A060BJA6-F1-model_v4 CAZy families CBM50\|GH19 protein | 17.1 | 9.42E-03 | uncultured Myxococcus sp. |
| 878 | AF-A0A1V3K3H7-F1-model_v4 Uncharacterized protein | 17.2 | 2.42E-04 | Rodentibacter pneumotropicus |
| 879 | AF-A0A818EHB9-F1-model_v4 Hypothetical protein | 16 | 3.94E-04 | Rotaria sp. Silwood1 |
| 880 | AF-A0A2W4UNG7-F1-model_v4 Uncharacterized protein | 31.2 | 4.99E-03 | Flavobacteriaceae bacterium |
| 881 | AF-A0A4U8S2F5-F1-model_v4 Glycoside hydrolase | 36 | 9.49E-04 | Helicobacter trogontum |
| 882 | AF-A0A839IBS2-F1-model_v4 Uncharacterized protein | 21.5 | 1.15E-03 | Aliivibrio sp. SR45-2 |
| 883 | AF-A0A0S8ZK03-F1-model_v4 Uncharacterized protein | 18 | 2.65E-03 | Pseudomonas sp. Leaf48 |
| 884 | AF-A0A4R7PA92-F1-model_v4 Chitinase class I | 15.9 | 2.07E-03 | Panacagrimonas perspica |
| 885 | AF-A0A819LNF6-F1-model_v4 Hypothetical protein | 18.2 | 6.75E-04 | Rotaria sp. Silwood1 |
| 886 | AF-A0A659LM02-F1-model_v4 Uncharacterized protein | 28.9 | 4.31E-03 | Escherichia sp. E2593 |
| 887 | AF-A0A380UI88-F1-model_v4 Chitinase | 21.6 | 2.78E-03 | Acinetobacter junii |
| 888 | AF-A0A0C1ERX9-F1-model_v4 Uncharacterized protein | 17.4 | 3.75E-04 | Kaistella jeonii |
| 889 | AF-A0A1H4XIM4-F1-model_v4 Putative chitinase | 25.7 | 2.65E-03 | Pseudomonas saponiphila |
| 890 | AF-A0A8B1B9M2-F1-model_v4 Uncharacterized protein | 22.9 | 2.78E-03 | Flavobacterium sp. CS20 |
| 891 | AF-A0A821PUA4-F1-model_v4 Hypothetical protein | 15.4 | 4.79E-04 | Rotaria socialis |
| 892 | AF-A0A2A9KEM7-F1-model_v4 Putative chitinase | 14.6 | 4.56E-04 | Collimonas sp. PA-H2 |
| 893 | AF-A0A1I5YMF8-F1-model_v4 Putative chitinase | 25.8 | 2.29E-03 | Pseudarcicella hirudinis |
| 894 | AF-A0A2X2X3E7-F1-model_v4 Predicted chitinase | 20.2 | 1.05E-03 | Chryseobacterium jejuense |
| 895 | AF-A0A1I5W7M0-F1-model_v4 Predicted chitinase | 18.4 | 7.81E-04 | Pseudarcicella hirudinis |
| 896 | AF-A0A817TJ42-F1-model_v4 Hypothetical protein | 16.2 | 5.28E-04 | Rotaria socialis |
| 897 | AF-A0A2A2GQ71-F1-model_v4 Uncharacterized protein | 18 | 1.15E-03 | Helicobacter sp. TUL |
| 898 | AF-A0A1Z8SBN4-F1-model_v4 Uncharacterized protein | 20.4 | 3.38E-03 | Planctomycetia bacterium TMED53 |
| 899 | AF-A0A816KCF6-F1-model_v4 Hypothetical protein | 14.1 | 1.27E-03 | Rotaria magnacalcarata |
| 900 | AF-A0A3P3W5I9-F1-model_v4 Uncharacterized protein | 21.4 | 4.79E-04 | Flavobacterium macacae |

| | | | | |
|---|---|---|---|---|
| 901 | AF-A0A3A1YCT5-F1-model_v4 Uncharacterized protein | 18.5 | 6.07E-03 | Capnocytophaga canis |
| 902 | AF-A0A069QI87-F1-model_v4 Peptidase, M23 family | 19.3 | 3.94E-04 | Prevotella loescheii DSM 19665 = JCM 12249 = ATCC 15930 |
| 903 | AF-I7GTM4-F1-model_v4 Uncharacterized protein | 29.5 | 1.05E-03 | Helicobacter cinaedi CCUG 18818 = ATCC BAA-847 |
| 904 | AF-A0A2T4TJZ7-F1-model_v4 Uncharacterized protein | 18.5 | 5.83E-04 | Prevotella sp. oral taxon 313 |
| 905 | AF-A0A6V7CDN3-F1-model_v4 Uncharacterized protein | 23.2 | 6.07E-03 | Xanthomonas hortorum pv. carotae |
| 906 | AF-A0A7C2EX51-F1-model_v4 Uncharacterized protein | 15.4 | 1.88E-03 | Blastocatellia bacterium |
| 907 | AF-A0A7W3FJL2-F1-model_v4 Peptidoglycan DD-metalloendopeptidase family protein | 21.1 | 2.40E-03 | Stenotrophomonas tumulicola |
| 908 | AF-A0A521QNZ5-F1-model_v4 Uncharacterized protein | 16.2 | 1.27E-03 | Reyranella sp. |
| 909 | AF-A0A7Y5LNG7-F1-model_v4 Glyco_hydro_19_cat domain-containing protein | 18.2 | 5.51E-03 | Myxococcales bacterium |
| 910 | AF-A0A522MSW4-F1-model_v4 Uncharacterized protein | 17.1 | 1.15E-03 | Paraburkholderia sp. |
| 911 | AF-A0A1Q2LFI1-F1-model_v4 Uncharacterized protein | 17.4 | 9.04E-04 | Helicobacter bilis |
| 912 | AF-A0A511F6Q1-F1-model_v4 Chitinase | 15.4 | 2.92E-03 | Methylorubrum extorquens |
| 913 | AF-A0A839VRP1-F1-model_v4 Hydroxyethylthiazole kinase | 17.7 | 4.79E-04 | Herbaspirillum sp. Sphag64 |
| 914 | AF-A0A0P4V6Z7-F1-model_v4 Glyco_hydro_19_cat domain-containing protein | 16.5 | 4.31E-03 | Leptolyngbya sp. NIES-2104 |
| 915 | AF-B1FQZ7-F1-model_v4 Uncharacterized protein | 21 | 1.71E-03 | Burkholderia ambifaria IOP40-10 |
| 916 | AF-A0A376EMZ3-F1-model_v4 Predicted chitinase | 17.8 | 5.51E-03 | Chryseobacterium carnipullorum |
| 917 | AF-A0A7X1FHY4-F1-model_v4 Glycoside hydrolase family 19 protein | 33.3 | 9.42E-03 | Pseudomonas sp. MSSRFD41 |
| 918 | AF-A0A2R7N6W0-F1-model_v4 Uncharacterized protein | 19.3 | 9.42E-03 | Chryseobacterium sp. HMWF028 |
| 919 | AF-A0A211ZQA2-F1-model_v4 Uncharacterized protein | 16.5 | 1.26E-02 | Inquilinus limosus |
| 920 | AF-A0A2T9Y5E9-F1-model_v4 Glyco_hydro_19_cat domain-containing protein | 19.1 | 1.55E-03 | Smittium simulii |
| 921 | AF-A0A0U3SWJ5-F1-model_v4 Putative endochitinase 3 | 17.8 | 6.69E-03 | Tachinaephagus zealandicus |
| 922 | AF-A0A817UUW6-F1-model_v4 Hypothetical protein | 16.2 | 2.29E-03 | Rotaria socialis |
| 923 | AF-A0A1I5YXZ1-F1-model_v4 Uncharacterized protein | 24.4 | 1.79E-03 | Pseudarcicella hirudinis |
| 924 | AF-A0A3C0GJM5-F1-model_v4 Glyco_hydro_19_cat domain-containing protein | 16.8 | 2.92E-03 | Maribacter sp. |
| 925 | AF-A0A846HPU2-F1-model_v4 Uncharacterized protein | 15.5 | 5.51E-03 | Hassallia byssoidea VB512170 |
| 926 | AF-A0A2T9ZAJ8-F1-model_v4 Glyco_hydro_19_cat domain-containing protein | 17.2 | 4.53E-03 | Smittium megazygosporum |
| 927 | AF-A0A562FNF0-F1-model_v4 Uncharacterized protein | 16.1 | 2.29E-03 | Aminobacter sp. J15 |
| 928 | AF-A0A851H659-F1-model_v4 Uncharacterized protein | 17.4 | 2.78E-03 | Aquitalea sp. LB_tupeE |
| 929 | AF-A0A443RY97-F1-model_v4 Glyco_hydro_19_cat domain-containing protein | 19.8 | 4.76E-03 | Leptotrombidium deliense |
| 930 | AF-A0A820SE06-F1-model_v4 Hypothetical protein | 25 | 1.39E-02 | Rotaria socialis |

| | | | | |
|---|---|---|---|---|
| 931 | AF-A0A2B7ZMX4-F1-model_v4 Uncharacterized protein | 25 | 1.20E-02 | Candidatus Nephrothrix sp. EaCA |
| 932 | AF-A0A1D1VPG4-F1-model_v4 Uncharacterized protein | 16.2 | 4.76E-03 | Ramazzottius varieornatus |
| 933 | AF-A0A2N7QV02-F1-model_v4 Uncharacterized protein | 17.1 | 5.51E-03 | Dyella sp. AD56 |
| 934 | AF-A0A511Q3S9-F1-model_v4 Chitinase | 13.4 | 5.51E-03 | Sphingomonas aquatilis NBRC 16722 |
| 935 | AF-A0A817ZY93-F1-model_v4 Hypothetical protein | 25.8 | 6.69E-03 | Rotaria socialis |
| 936 | AF-A0A814H2K5-F1-model_v4 Hypothetical protein | 19.4 | 7.38E-03 | Brachionus calyciflorus |
| 937 | AF-A0A223NX21-F1-model_v4 Chitinase | 14.2 | 3.04E-02 | Mucilaginibacter xinganensis |
| 938 | AF-A0A6A7RW67-F1-model_v4 Uncharacterized protein | 19.8 | 3.88E-02 | Candidatus Accumulibacter phosphatis |
| 939 | AF-A0A3C0AX61-F1-model_v4 Uncharacterized protein | 16.7 | 8.55E-03 | Flavobacterium sp. |
| 940 | AF-A0A443Q798-F1-model_v4 Glyco_hydro_19_cat domain-containing protein | 17.2 | 5.51E-03 | Dinothrombium tinctorium |
| 941 | AF-A0A5C7P5Z3-F1-model_v4 Uncharacterized protein | 17.4 | 2.18E-03 | Desulfurellales bacterium |
| 942 | AF-Q92UB6-F1-model_v4 Putative glycoside hydrolase | 28.1 | 1.46E-02 | Sinorhizobium meliloti 1021 |
| 943 | AF-A0A1C3XH69-F1-model_v4 Uncharacterized protein | 18 | 1.09E-02 | Bradyrhizobium shewense |
| 944 | AF-A0A7D7ZX30-F1-model_v4 Glyco_hydro_19_cat domain-containing protein | 29.8 | 1.26E-02 | Flavobacteriaceae bacterium |
| 945 | AF-A0A059ENZ3-F1-model_v4 Glyco_hydro_19_cat domain-containing protein | 14.4 | 9.89E-03 | Anncaliia algerae PRA109 |
| 946 | AF-A0A1D1VYW9-F1-model_v4 Glyco_hydro_19_cat domain-containing protein | 14.9 | 1.69E-02 | Ramazzottius varieornatus |
| 947 | AF-A6NY60-F1-model_v4 Putative peptidoglycan binding domain protein | 15.4 | 9.42E-03 | Pseudoflavonifractor capillosus ATCC 29799 |
| 948 | AF-A0A1D2N1Y0-F1-model_v4 Fibroblast growth factor receptor | 19 | 7.38E-03 | Orchesella cincta |
| 949 | AF-A0A3S3P3P2-F1-model_v4 Glyco_hydro_19_cat domain-containing protein | 18.4 | 1.09E-02 | Dinothrombium tinctorium |
| 950 | AF-A0A7W6ICU4-F1-model_v4 Putative chitinase | 16.7 | 1.61E-02 | Microvirga flocculans |
| 951 | AF-A0A3A9IVM8-F1-model_v4 DUF4280 domain-containing protein | 15.9 | 1.26E-02 | Anaerotruncus sp. 1XD22-93 |
| 952 | AF-A0A2B8A0M4-F1-model_v4 Uncharacterized protein | 26.2 | 3.52E-02 | Candidatus Nephrothrix sp. EaCA |
| 953 | AF-A0A256A0E5-F1-model_v4 Uncharacterized protein | 17.5 | 1.87E-02 | Flavobacterium aurantiibacter |
| 954 | AF-A0A6J0M4Y4-F1-model_v4 structural maintenance of chromosomes protein 3-like | 15.6 | 1.33E-02 | Raphanus sativus |
| 955 | AF-A0A375I2Q7-F1-model_v4 Chitinase | 12.3 | 3.19E-02 | Propionibacterium ruminifibrarum |
| 956 | AF-A0A5B8YLA5-F1-model_v4 Core-binding (CB) domain-containing protein | 24.5 | 1.78E-02 | Antarcticibacterium arcticum |
| 957 | AF-A0A846A811-F1-model_v4 S8 family serine peptidase | 14.1 | 1.26E-02 | Leptolyngbya sp. SIO4C5 |
| 958 | AF-A0A437AQH0-F1-model_v4 Endochitinase | 14.6 | 2.38E-02 | Tubulinosema ratisbonensis |
| 959 | AF-A0A2S1LK33-F1-model_v4 Uncharacterized protein | 15 | 7.75E-03 | Flavobacterium kingsejongi |
| 960 | AF-A0A1D2MHY3-F1-model_v4 Metalloendopeptidase | 17.3 | 1.96E-02 | Orchesella cincta |

| | | | | |
|---|---|---|---|---|
| 961 | AF-A0A443S307-F1-model_v4 Glyco_hydro_19_cat domain-containing protein | 16.2 | 2.38E-02 | Leptotrombidium deliense |
| 962 | AF-A0A2D5AIF7-F1-model_v4 Uncharacterized protein | 15.2 | 2.50E-02 | Verrucomicrobiales bacterium |
| 963 | AF-A0A4Q9LJ66-F1-model_v4 Putative chitinase | 17.3 | 1.78E-02 | Hamiltosporidium magnivora |
| 964 | AF-A0A4U8S2I0-F1-model_v4 Uncharacterized protein | 30.9 | 9.82E-02 | Helicobacter trogontum |
| 965 | AF-A0A1J5WNL5-F1-model_v4 Endochitinase | 18.3 | 2.06E-02 | Amphiamblys sp. WSBS2006 |
| 966 | AF-A0A2E8A917-F1-model_v4 Uncharacterized protein | 16.5 | 2.16E-02 | Marinobacter sp. |
| 967 | AF-A0A814L726-F1-model_v4 Hypothetical protein | 18.5 | 1.87E-02 | Didymodactylos carnosus |
| 968 | AF-A0A2T9ZAE4-F1-model_v4 Glyco_hydro_19_cat domain-containing protein | 16.6 | 1.96E-02 | Smittium megazygosporum |
| 969 | AF-L1P4X3-F1-model_v4 Uncharacterized protein | 16.7 | 2.76E-02 | Capnocytophaga sp. oral taxon 332 str. F0381 |
| 970 | AF-A0A1R0GP95-F1-model_v4 Endochitinase PR4 | 14.5 | 3.19E-02 | Smittium mucronatum |
| 971 | AF-A0A3M7Q777-F1-model_v4 Chitinase 4-like | 16.4 | 4.72E-02 | Brachionus plicatilis |
| 972 | AF-A0A6D2GAN9-F1-model_v4 Predicted chitinase | 14.9 | 7.75E-03 | Salmonella enterica subsp. salamae |
| 973 | AF-A0A2G5BL59-F1-model_v4 Lysozyme-like protein | 15.9 | 2.06E-02 | Coemansia reversa NRRL 1564 |
| 974 | AF-A0A7J9CLG7-F1-model_v4 Chitin-binding type-1 domain-containing protein | 14.4 | 1.54E-02 | Gossypium gossypioides |
| 975 | AF-A0A3E0C4S4-F1-model_v4 Putative chitinase | 13.8 | 3.70E-02 | Paraburkholderia sp. BL6669N2 |
| 976 | AF-A0A1H2ZVF6-F1-model_v4 Predicted chitinase | 17.5 | 2.38E-02 | Capnocytophaga granulosa |
| 977 | AF-A0A1Z9VEL6-F1-model_v4 Uncharacterized protein | 14.2 | 1.09E-02 | Gammaproteobacteria bacterium TMED281 |
| 978 | AF-B0XK59-F1-model_v4 Basic endochitinase CHB4 | 17.4 | 1.14E-01 | Culex quinquefasciatus |
| 979 | AF-A0A659AS86-F1-model_v4 Uncharacterized protein | 16.8 | 2.76E-02 | Vibrio parahaemolyticus |
| 980 | AF-A0A512J235-F1-model_v4 Uncharacterized protein | 12.6 | 1.32E-01 | Methylobacterium oxalidis |
| 981 | AF-A0A250F2N9-F1-model_v4 Uncharacterized protein | 15.5 | 3.04E-02 | Capnocytophaga sputigena |
| 982 | AF-A0A0R0M0W5-F1-model_v4 Putative chitinase | 16.3 | 6.64E-02 | Pseudoloma neurophilia |
| 983 | AF-A0A2D4BNH4-F1-model_v4 Uncharacterized protein | 14.9 | 2.15E-01 | Pythium insidiosum |
| 984 | AF-A0A2N3I860-F1-model_v4 Uncharacterized protein | 14.3 | 1.60E-01 | Raineya orbicola |
| 985 | AF-A0A0M4PV89-F1-model_v4 Uncharacterized protein | 15 | 2.48E-01 | Pseudonocardia sp. AL041005-10 |
| 986 | AF-A0A814PZA7-F1-model_v4 Hypothetical protein | 17.7 | 1.85E-01 | Brachionus calyciflorus |
| 987 | AF-A0A349DNN6-F1-model_v4 Uncharacterized protein | 14.8 | 2.76E-02 | Microscillaceae bacterium |
| 988 | AF-A0A437AM80-F1-model_v4 Endochitinase | 15 | 1.68E-01 | Tubulinosema ratisbonensis |
| 989 | AF-A1ZH11-F1-model_v4 Uncharacterized protein | 16.4 | 5.20E-02 | Microscilla marina ATCC 23134 |
| 990 | AF-A0A423XPF7-F1-model_v4 Uncharacterized protein | 10.7 | 3.17E-01 | Cytospora leucostoma |
| 991 | AF-A0A2H9TFJ6-F1-model_v4 Uncharacterized protein | 19.3 | 4.50E-02 | Paramicrosporidium saccamoebae |
| 992 | AF-R0MMH7-F1-model_v4 Acidic endochitinase SP2 | 17.2 | 1.60E-01 | Nosema bombycis CQ1 |
| 993 | AF-A0A7W5WSN6-F1-model_v4 Putative chitinase | 27.3 | 1.67E+00 | Rhizobium sp. BK393 |

| | | | | |
|---|---|---|---|---|
| 994 | AF-A0A0G2FKG1-F1-model_v4 Uncharacterized protein | 10.1 | 4.25E-01 | Diaporthe ampelina |
| 995 | AF-R1CQG6-F1-model_v4 Uncharacterized protein | 11.9 | 1.52E-01 | Emiliania huxleyi |
| 996 | AF-A0A1V3JU95-F1-model_v4 Uncharacterized protein | 22.7 | 1.19E+00 | Rodentibacter sp. Ppn85 |
| 997 | AF-A0A2P6TDK2-F1-model_v4 Class VII chitinase | 16.8 | 1.93E+00 | Chlorella sorokiniana |
| 998 | AF-A0A0Q6P000-F1-model_v4 Uncharacterized protein | 16.4 | 2.88E-01 | Devosia sp. Root105 |
| 999 | AF-A0A1G5ZXP1-F1-model_v4 Uncharacterized protein | 16.9 | 2.61E-01 | Sinorhizobium sp. NFACC03 |
| 1000 | AF-A1ZW01-F1-model_v4 Uncharacterized protein | 16.9 | 5.98E-01 | Microscilla marina ATCC 23134 |
